# Covalent Targeting of Splicing in T Cells

**DOI:** 10.1101/2023.12.18.572199

**Authors:** Kevin A. Scott, Hiroyuki Kojima, Nathalie Ropek, Charles D. Warren, Tiffany L. Zhang, Simon J. Hogg, Caroline Webster, Xiaoyu Zhang, Jahan Rahman, Bruno Melillo, Benjamin F. Cravatt, Jiankun Lyu, Omar Abdel-Wahab, Ekaterina V. Vinogradova

## Abstract

Despite significant interest in therapeutic targeting of splicing, few chemical probes are available for the proteins involved in splicing. Here, we show that elaborated stereoisomeric acrylamide chemical probe EV96 and its analogues lead to a selective T cell state-dependent loss of interleukin 2-inducible T cell kinase (ITK) by targeting one of the core splicing factors SF3B1. Mechanistic investigations suggest that the state-dependency stems from a combination of differential protein turnover rates and availability of functional mRNA pools that can be depleted due to extensive alternative splicing. We further introduce a comprehensive list of proteins involved in splicing and leverage both cysteine- and protein-directed activity-based protein profiling (ABPP) data with electrophilic scout fragments to demonstrate covalent ligandability for many classes of splicing factors and splicing regulators in primary human T cells. Taken together, our findings show how chemical perturbation of splicing can lead to immune state-dependent changes in protein expression and provide evidence for the broad potential to target splicing factors with covalent chemistry.

## Introduction

Small molecule probes are central to the drug discovery process and can serve as starting points for the development of new therapeutics. Unlike gene editing approaches, chemical probes enable studies of the role of both acute and long-term perturbation of protein function and can be easily applied to different cell types, including primary human cells, providing unique insights into cell-type specific roles of these proteins. Electrophilic compounds have emerged as a new class of covalent chemical probes that provide unprecedented opportunities for targeting proteins and sites on proteins that have been historically inaccessible using standard non-covalent small-molecule approaches[1–3]. Combining covalent probes with advanced mass-spectrometry (MS)- enabled chemical proteomic workflows such as activity-based protein profiling (ABPP) has been instrumental in global mapping of thousands of ligandable sites in proteins in native biological systems[4, 5]. However, more potent and selective chemical probes are still missing for the majority of proteins due to the time-consuming nature of target-focused ligand optimization efforts. Phenotypic screening using covalent electrophiles can serve as a complementary target-agnostic platform for the discovery of advanced small molecules with unique mechanisms-of-action, including protein degradation[6] and stabilization of mRNA-bound splicing factor complexes[7]. The use of covalent electrophiles further facilitates broad-scale target ID studies enabled by advanced chemical proteomic platforms (TMT-ABPP)[1, 8].

Splicing factor SF3B1 is a major component of the U2 spliceosome and is commonly mutated in hematopoietic cancers[9] and myelodysplastic syndromes[10]. As a result, SF3B1 has become an attractive target for therapeutic development for treating blood cancers and solid tumors[11] and is currently the subject of several clinical trials[12, 13]. Recently, a set of stereochemically defined tryptoline acrylamides was found to covalently engage SF3B1 at C1111, leading to remodeling of the spliceosome and altered splicing events in human cancer cells[14]. Here, we show that SF3B1 is a phenotypically relevant target of the tryptoline acrylamide EV96 found previously in a screen for inhibitors of T cell activation[6]. We further show that inhibition of SF3B1 with EV96 or natural product Pladienolide B (PladB) leads to loss of immune kinase ITK specifically in stimulated, but not quiescent T cells, and generally overlapping changes in global gene and protein expression. Our mechanistic investigations into the cell state-dependency of ITK loss suggest that it is likely due to a combination of differences in protein turnover rates and depletion of functional mRNA pools by alternative splicing. This finding, together with the lack of gross protein expression changes caused by chemical perturbation of SF3B1 in quiescent T cells, suggests that there may be unique opportunities for targeting splicing in the context of immunopathologies, which have remained largely unexplored to date.

Precursor messenger RNA splicing is a ubiquitous mechanism that generates multiple transcripts from a single gene, enabling a large increase in structural and functional complexity in the proteome. It has recently become clear that alternative splicing plays a central role in biological processes that regulate immune cell function, including the maintenance of quiescence, activation, and effector function in T cells[15–17]. Furthermore, many viruses require host splicing machinery for replication, and alternative splicing of canonical host proteins is often implicated in viral infection[18–20]. Abnormal splicing events can lead to cellular dysfunction and pathological states including autoimmune conditions[21, 22] and cancer[23]. Inspired by our findings herein that SF3B1 modulators[14] suppress T-cell activation, as well as the discovery of covalent ligands for other splicing factors like NONO[7], we decided to perform a more in-depth evaluation of the broad potential of targeting splicing with covalent electrophiles. We generated a comprehensive list of proteins that are involved in splicing, either directly or in a regulatory capacity. We further leveraged cysteine-directed TMT-ABPP data using scout fragments KB02 and KB05 from activated and quiescent T cells[6] to identify ligandable sites and annotated their functional relevance by integrating information from existing databases on specific protein domains harboring these sites. We finally generated and incorporated protein-directed TMT-ABPP data to expand the list of ligandable splicing factors and regulators and nominate specific targets for further ligand development efforts. To our knowledge, this represents the most exhaustive resource of ligandable splicing factors to date.

## RESULTS

### Investigating ITK as a direct target of EV96

Our recent study has shown that phenotypic screening of covalent electrophiles can serve as a powerful target-agnostic approach to identifying small molecules that suppress T cell activation by non-canonical mechanisms, including protein degradation (Fig. 1A)[6]. Among these, we evaluated a set of 4 stereoisomeric tryptoline-based acrylamides, one of which (EV96) inhibited T cell activation with high stereoselectivity (Fig. 1A-1B). Further investigation into the mechanism of suppression of T cell activation revealed a stereoselective decrease in the expression of a major immune kinase ITK[24] (Fig. 1A); however, the mechanism of this reduction in ITK remained unclear. Interestingly, ITK reductions occurred only under TCR-stimulating conditions, which motivated our interest in understanding the mechanism of this cell state-dependent pharmacology.

**Figure 1.**
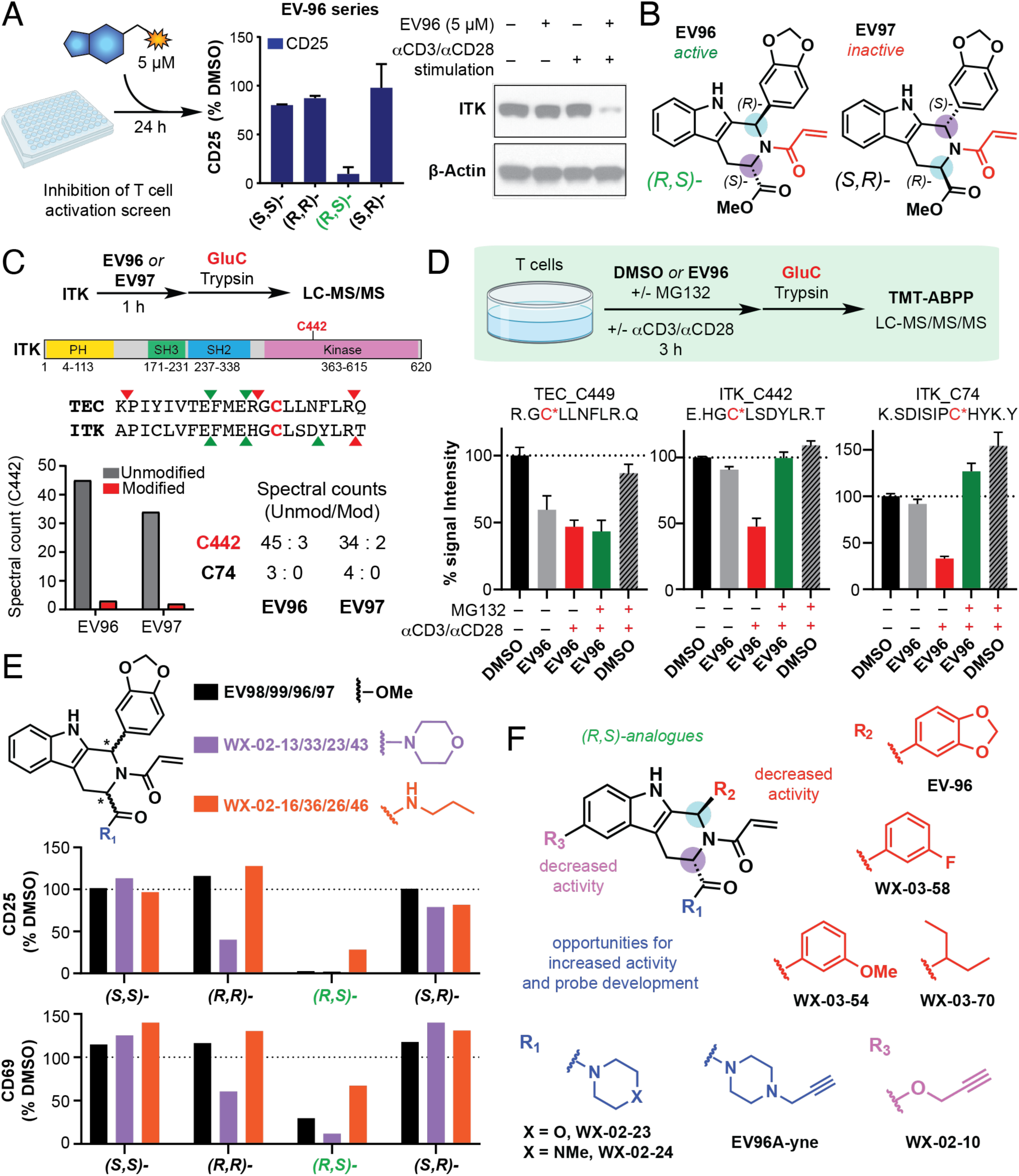
Structure-activity relationship study identifies EV96A-yne as a potent EV96-based chemical probe for target ID studies. (A) Elaborated tryptoline scaffold EV96 leads to stereoselective suppression of T cell activation and T cell state-dependent downregulation of ITK protein levels[1]. Primary human T cells were treated in 96-well plates with either compounds (5 µM) or DMSO control. Expression of early activation marker CD25 was measured using flow cytometry to determine the extent of suppression of T cell activation. The *(R,S)-*β-carboline scaffold EV96, but not its enantiomer EV97, or diastereomers EV98 and EV99, was active in suppressing T cell activation at 5 μM concentration, leading to simultaneous stereoselective and T cell state-dependent loss of ITK. (B) Structures of EV96 and EV97 with the electrophilic acrylamide highlighted in red and stereocenters in blue and purple. (C) *In vitro* evaluation of ITK_C442 engagement with active compound EV96 and inactive control EV97 using tandem trypsin/GluC protease digest. Alignment of ITK and TEC sequences, showing cysteine of interest in bold (red) and predicted trypsin (red) and GluC (green) cleavage sites. Full length ITK protein (40 nM) was treated with EV96 (1 µM) or EV97 (1 µM) for 1 h, followed by standard proteomic processing for LC-MS/MS analysis. Tandem trypsin/GluC digest allowed for quantification of the elusive peptide bearing C442. (D) *In situ* cell based evaluation of ITK_C442 engagement with EV96 using TMT-ABPP proteomic workflows with tandem trypsin/GluC protease digest. Primary human T cells were treated with DMSO or EV96 (5 µM) for 3 h with or without stimulation with αCD3/αCD28 antibodies. Addition of a proteasome inhibitor MG132 (10 µM) was used to test for loss of signal due to protein degradation. Tandem trypsin/GluC digest allowed for quantification of the elusive peptide bearing C442. Signal intensity is calculated based on the competition level for IA-DTB labeling compared to DMSO control. Bar graph representation of changes in signal intensities for two quantified ITK peptides and TEC_C449 peptide are shown, highlighting the differences between ligand engagement and protein degradation. (E) Amide analogues of EV96 show similar stereoselectivity of suppression of T cell activation. T cells were treated with 3 sets of stereoisomers (5 µM) with αCD3/αCD28 stimulation and suppression of T cell activation was measured using flow cytometry analysis of CD25 and CD69 expression following 24 h compound treatment. (F) Structure-activity relationship study leads to identification of potent analogues bearing alkyne handle for visualization and enrichment. T cells were treated with EV96 analogues at 2 different concentrations (5 µM and 2 µM) for 24 h. Suppression of T cell activation was measured by flow cytometry analysis of CD25 expression. Reduction of ITK levels was measured using Western blot analysis and correlated with suppression of T cell activation.

Our original study identified that EV96 stereoselectively engages another protein from the Tec kinase family, TEC, at a cysteine residue (C449)[6] that is conserved in ITK, initially suggesting that EV96 might also directly bind ITK. However, the tryptic peptide containing the corresponding cysteine in ITK (C442) was not observed in cysteine-directed ABPP experiments. To quantify ITK_C442 reactivity, we performed ABPP experiments using a tandem proteolysis protocol with trypsin and GluC proteases, which was expected to produce a shorter C442- containing peptide by introducing additional cut sites after aspartic and glutamic acid residues (Fig. 1C). Indeed, this strategy allowed us to quantify the peptide containing ITK_C442 both following *in vitro* treatment of purified ITK protein with EV96 and its inactive enantiomer (Fig. 1C), as well as cell-based treatment with and without αCD3/αCD28 stimulation (Fig. 1D). We did not observe direct engagement of ITK_C442 by EV96 in experiments performed with purified protein (Fig. 1C, Supplementary Tables S1A, S1B). For assessing the potential engagement of ITK by EV96 in stimulated T cells, we included control treatments with a proteasome inhibitor MG132 that prevented EV96-induced decreases in ITK protein. These experiments confirmed the expected direct engagement of TEC_C449 by EV96, which was observed in both control and MG132-treated T cells (Fig. 1D, Supplementary Table S2). In contrast, the EV96-induced decrease in ITK_C442 signals (as well as signals for other cysteine-containing peptides from ITK) was prevented by treatment with MG132 (Fig. 1D, S1A), indicating that ITK was likely not a direct target of EV96.

### Structure-activity relationship studies identify more potent EV96 analogues

The original target ID study was performed using competition assays with a broadly reactive iodoacetamide desthiobiotin (IA-DTB) chemical probe and analysis of cysteine-containing peptides following enrichment (cysteine-directed TMT-ABPP)[6]. This approach is often used for identification of targets without the need for derivatization of the original hit compounds, however it can overlook cysteine-containing peptides that do not ionize as well and are harder to detect using mass-spectrometry analysis[25]. We therefore wanted to perform additional competition-based experiments with alkyne-containing chemical probes based on the original hit compound to identify any targets that escaped our original detection methods (protein-directed TMT-ABPP)[14, 25].

To identify more potent EV96-based alkyne-containing chemical probes, we performed a focused structure-activity relationship (SAR) study using all four stereoisomeric compounds and the substitution of the ester group. This study confirmed stereospecificity of T cell suppression for the *(R,S)*-stereoisomers and nominated the methyl ester as a potential exit vector for further lead optimization and chemical probe development (Fig. 1E) with morpholine-substituted amides (WX-02-23) identified as the most potent suppressors of T cell activation. Additional study using *(R,S)*-stereoisomers revealed a tight SAR around the aryl group substituent and the tryptophan ring of the tryptoline scaffold (Fig. 1F, leading to decreased potency in both suppression of T cell activation and ITK abundance (Fig. S1B-S1F), which showed good correlation. Substitution of the morpholine group with a derivative *N-*methyl-piperidine group (WX-02-24) was further tolerated and allowed us to synthesize an alkynylated chemical probe EV96A-yne for further analyses (Fig. 1F, S2A-S2B).

### Identification of SF3B1 as a phenotypically relevant EV96 target

We next performed protein-directed ABPP experiments with the EV96A-yne probe (Fig. 2A, Supplementary Table S3), where we evaluated active compound EV96, inactive enantiomer EV97, and an inactive state ITK inhibitor PF-06465469 (henceforth, PF; Fig. 2B)[26], which is known to covalently engage ITK_C442, for effects on EV96A-yne enrichment of proteins. T cells were treated with the compounds for 3 h before EV96A-yne treatment for 2 h. Subsequent copper-catalyzed azide-alkyne cycloaddition (CuAAC) reaction with biotin azide and streptavidin enrichment followed by TMT-MS analysis allowed for identification and quantitation of EV96A-yne-labeled proteins. Three proteins were found to be stereospecifically competed by EV96 (>75% competition), but not EV97 that were not quantified in our original cysteine-directed ABPP experiments – AGL, MYO1G, and SF3B1. The same three targets were also identified in an experiment with Jurkat cells, which represent a constitutively activated T cell leukimia line (Fig S2C, Supplementary Table S4). ITK was not identified as a direct target of EV96A-yne in these protein-directed ABPP experiments. Stereospecific competition of EV96A-yne labeling of proteins matching the predicted molecular weight of SF3B1 (146 kDa) and MYO1G (116 kDa) was also visualized by gel-ABPP (CuAAC conjugation to a rhodamine azide reporter group followed by SDS-PAGE and in-gel fluorescence scanning) (Fig. 2C). Among the proteins stereoselectively liganded by EV96, SF3B1 was of particular interest, because, in a parallel study, we discovered that tryptoline acrylamides engaged C1111 in SF3B1 with a similar SAR to the T-cell suppression effects observed herein, leading to remodeling of the spliceosome and substantial alterations in splicing in human cancer cells [14]. SF3B1_C1111 is located on a nonproteotypic tryptic peptide[14] that is not typically quantified in cysteine-directed ABPP experiments. We noted that SF3B1_C1111 was conserved in mouse species (Fig. S2D) and further validated that the same state-dependent ITK loss phenotype was observed in mouse T cells isolated from mouse spleens (Fig. S2E).

**Figure 2.**
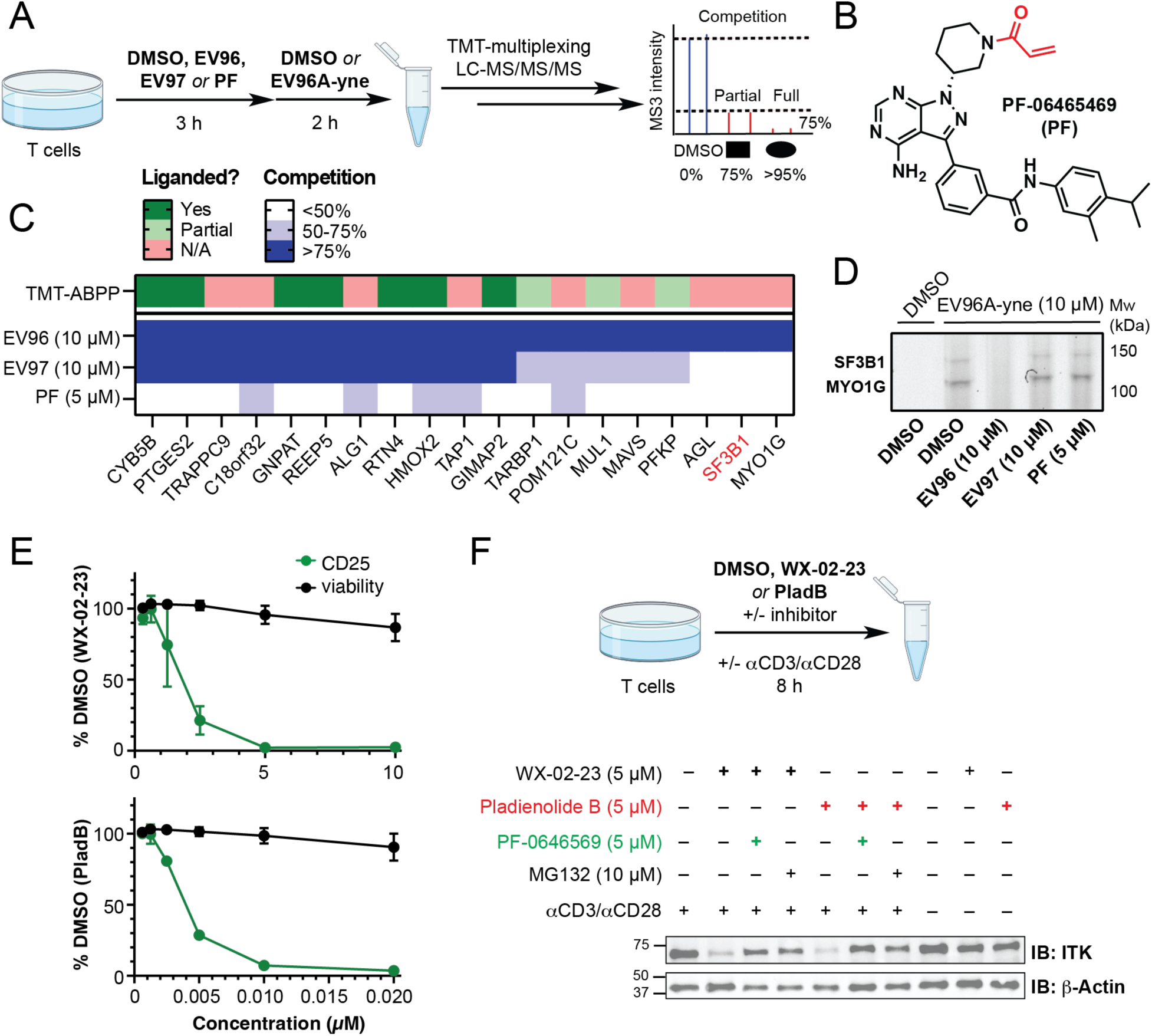
Identification of SF3B1 as the phenotypically-relevant target of EV96. (A) Schematic of a TMT-ABPP competition experiment in primary human T cells using EV96A-yne chemical probe (5 µM, 2 h) following pre-treatment with DMSO, EV96 (10 µM), EV97 (10 µM), and PF-06465469 (PF, 5 µM) for 3 h. (B) Structure of PF-06465469 (PF) – a published covalent inhibitor of cysteine 442 of IL2-inducible T-cell kinase (ITK_C442), which locks the kinase in its inactive confirmation. (C) Heatmap showing percent pulldown competition following compound treatment and evidence of ligandability with EV96 in a previous cysteine-directed TMT-ABPP study[1]. Data are average values from 2-3 independent experiments. (D) Gel-based ABPP visualization of EV96A-yne probe labeling and competition using copper-catalyzed azide-alkyne cycloaddition (CuAAC) reaction with rhodamine-azide. Competition for SF3B1 and MYO1G labeling was observed following pre-treatment with EV96 (10 µM), but not its inactive enantiomer EV97 (10 µM) or PF-06465469 (5 µM). (E) WX-02-23 and a known non-covalent SF3B1 inhibitor Pladienolide B showed the same concentration-dependent effects on suppression of T cell activation. Viability and CD25 expression were evaluated using flow cytometry following staining with near-IR live-DEAD dye and PE anti-CD25 antibody. Data are mean values ± SD from 2 independent experiments. (F) Both WX-02-23 (5 µM) and Pladienolide B (5 µM) lead to T cell state-dependent downregulation of ITK levels, which is rescued by the addition of proteasome inhibitor MG132 (10 µM) and pre-treatment with ITK_C442 inhibitor PF-06465469 (5 µM). Western blot showing ITK levels following inhibitor treatment for 8 h. Data are from a single experiment representative of three independent biological experiments.

The splicing factor SF3B1 is a key component of the U2 spliceosome and is frequently mutated in specific hematological cancers[27]. Despite the importance of splicing for physiology and disease, only a limited set of natural product-based chemical tools exist for studying this process, including pladienolide natural products, some of which are currently in clinical trials in patients with myelodysplastic syndromes (MDS), acute myeloid leukemia (AML) or chronic myelomonocytic leukemia[28, 29]. We found that pladienolide B (PladB) – a natural product that competitively binds with tryptoline acrylamides to SF3B1[14] – also suppressed T cell activation (Fig. 2E) and caused activation state-dependent loss of ITK (Fig. 2F) in human T cells. As reported previously for EV96[6], the decrease in ITK caused by WX-02-23 or PladB was attenuated by co-treatment with MG132 or the ITK inhibitor PF (Fig. 2F). The shared binding of EV96 and PladB to SF3B1 and similar pharmacological effects displayed by these compounds in T cells supported that SF3B1 modulation may underpin the state-dependent reductions in ITK.

### Structural modeling of EV96 binding to SF3B1

Covalent docking of active compounds EV96 and WX-02-23 using a co-crystal structure of the SF3B complex and PHF5A (PDB:7B91), provides a rationale for the observed SAR of SF3B1 engagement (Fig. S2F). In this model, the indole ring projects deep into the binding pocket, and the 3,4-methylenedioxoarene projects into a well-formed cavity. This is consistent with the observation that nearly any change to the 3,4-methylenedioxoarene results in a reduction of activity. The model also predicts a potential hydrogen-bond formation between the -NH of the indole and the backbone carbonyl of L1066 (Fig. S2F), which might contribute to stabilization of non-covalent interactions between EV96/WX-02-23 and SF3B1. Furthermore, while the binding pocket is tight around the aryl group substituent and tryptophan ring, there is a large hydrophobic cavity adjacent to the methyl ester of EV96 in this model, which is consistent with increased potency of WX-02-23 and can provide additional opportunities for further optimization of WX-02-23 potency and selectivity.

### WX-02-23 and PladB induce comparable global gene and protein level changes in T cells

To further compare WX-02-23 and PladB, we characterized global protein and transcript changes by MS-based proteomics (TMT-exp, Supplementary Table S5) and RNA sequencing (RNA-seq, Supplementary Table 2S-1), respectively (Fig. 3A). We compared protein expression changes across cells treated with DMSO, WX-02-23, WX-02-43, and PladB with and without αCD3/αCD28 stimulation by 16plex TMT-exp (each condition tested in duplicate, Fig. 3A–3C). This experiment confirmed state-dependent loss of ITK and several other proteins (PRF1, CBLB, IL27RA) induced by the more potent analogue WX-02-23, but not the inactive enantiomer WX-02-43 following 8 h treatment (Fig. 3B, left; Supplementary Table S5). Consistent with previous

**Figure 3.**
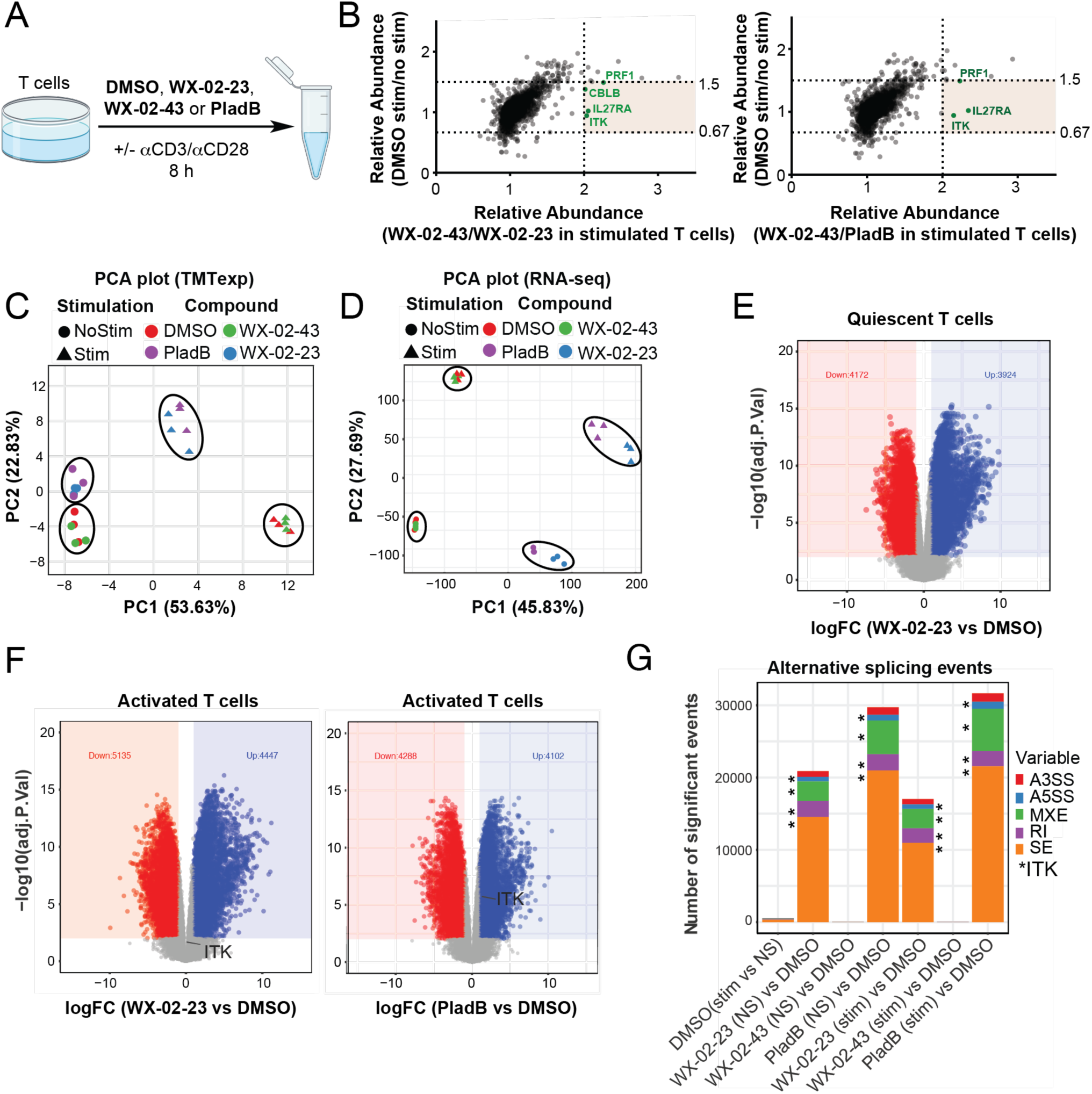
Comparison of global mRNA and protein expression changes following WX-02-23 and PladB treatment. (A) T cells were treated with DMSO, WX-02-23 (5 µM), WX-02-43 (5 µM), or PladB (5 µM) in the presence or absence of external stimulation and processed for RNA-seq or LC-MS/MS/MS analysis as part of a standard TMT 16-plex experiment. (B) Scatter plots showing selective downregulation of ITK and several other proteins following WX-02-23 and PladB treatment. (C-D) Principal component analysis of protein (C) and mRNA (D) abundance data obtained using unenriched proteomics and RNA-sequencing experiments comparing stimulated (triangle) and quiescent (circle) T cells, following treatment with DMSO (red), WX-02-23 (blue), WX-02-43 (green), and Pladienolide B (purple). Data are average values from n=3 biological replicates. Clustering of WX-02-23 and PladB conditions suggests similar global protein and mRNA abundance changes of the two SF3B1 inhibitors. (E-F) Volcano plots showing substantial (>2 log2 fold change) and statistically significant (FDR <0.05) gene expression changes in RNA-sequencing experiments of quiescent and activated T cells following WX-02-23 (5 µM) and PladB (5 µM) treatment for 8 h. Data are average values from n=3 biological replicates. Global gene expression changes indicate that SF3B1 is likely inhibited in both T cell states and ITK protein expression is regulated post-transcriptionally. (G) Alternative splicing events in quiescent and stimulated T cells following DMSO, WX-02-23 (5 µM), WX-02-43 (5 µM), or PladB (5 µM) treatment for 8 h. Splicing changes were identified from 3 biological replicates by rMATS with the requirement of ΔPSI > 0.2 compared to DMSO (NS) condition and FDR < 0.05. PSI – percent spliced in, FDR – false discovery rate, A3SS – alternative 3’ splice site, A5SS – alternative 5’ splice site, MSE – mutually-exclusive exon, RI – retained intron, SE – skipped exon. Asterisk denotes ITK alternative splicing events in different treatment groups. (See Supplementary Methods for details).

Western blot data, the same state-dependent decrease in ITK levels was observed in proteomic data of PladB treated samples (Fig. 3B, right). Principal component analysis (PCA) of protein (Fig. 3C) and gene (Fig. 3D) expression data further suggested concordant global effects following WX-02-23 and PladB treatment, as these conditions clustered together in both stimulated (triangles) and quiescent (circles) T cells, while WX-02-43 treatment clustered together with DMSO treated samples. Notably, changes at the gene expression level were much more pronounced than at the protein level following treatment with either WX-02-23 or PladB (Fig. 3E, 3F). Proteomic changes in quiescent T cells largely clustered together regardless of treatment (DMSO/WX-02-23/WX-02-43/PladB). Of note, we observed global transcriptional changes following compound treatment of both quiescent and activated cells, which indicates that SF3B1_C1111 engagement occurred in both cell states. ITK transcripts were not significantly altered in abundance following either WX-02-23 or PladB treatment of activated cells (Fig. 3F). Additionally, changes in global mRNA levels and protein levels induced by PladB and WX-02-23 were highly correlated in activated T cells (Pearson’s correlation coefficient of 0.847 and 0.927, respectively) (Fig. S3A, S3B).

Considering the major role of SF3B1 in regulating mRNA splicing, we investigated alternative splicing events produced by treatment with WX-02-23, PladB, and WX-02-43 with and without αCD3/αCD28 stimulation. Both WX-02-23 and PladB, but not WX-02-43, induced a substantial number of alternative splicing events in both activated and quiescent T cells (Fig. 3G, Supplementary tables 2S-2), including exon skipping (ES), mutually exclusive exons (MXE), retained introns (RI), and alternative 3’ and 5’ splice sites (A3SS, A5SS). Notably, ITK mRNA was alternatively spliced in T cells treated with WX-02-23 or PladB, showing multiple skipped exons, a retained intron, alternative 5’ splice site, and a mutually exclusive exon in the presence or absence of αCD3/αCD28 stimulation (Fig. 3G, 4A). These ITK splicing effects were observed in both quiescent and stimulated T cells (Fig. 3G).

**Figure 4.**
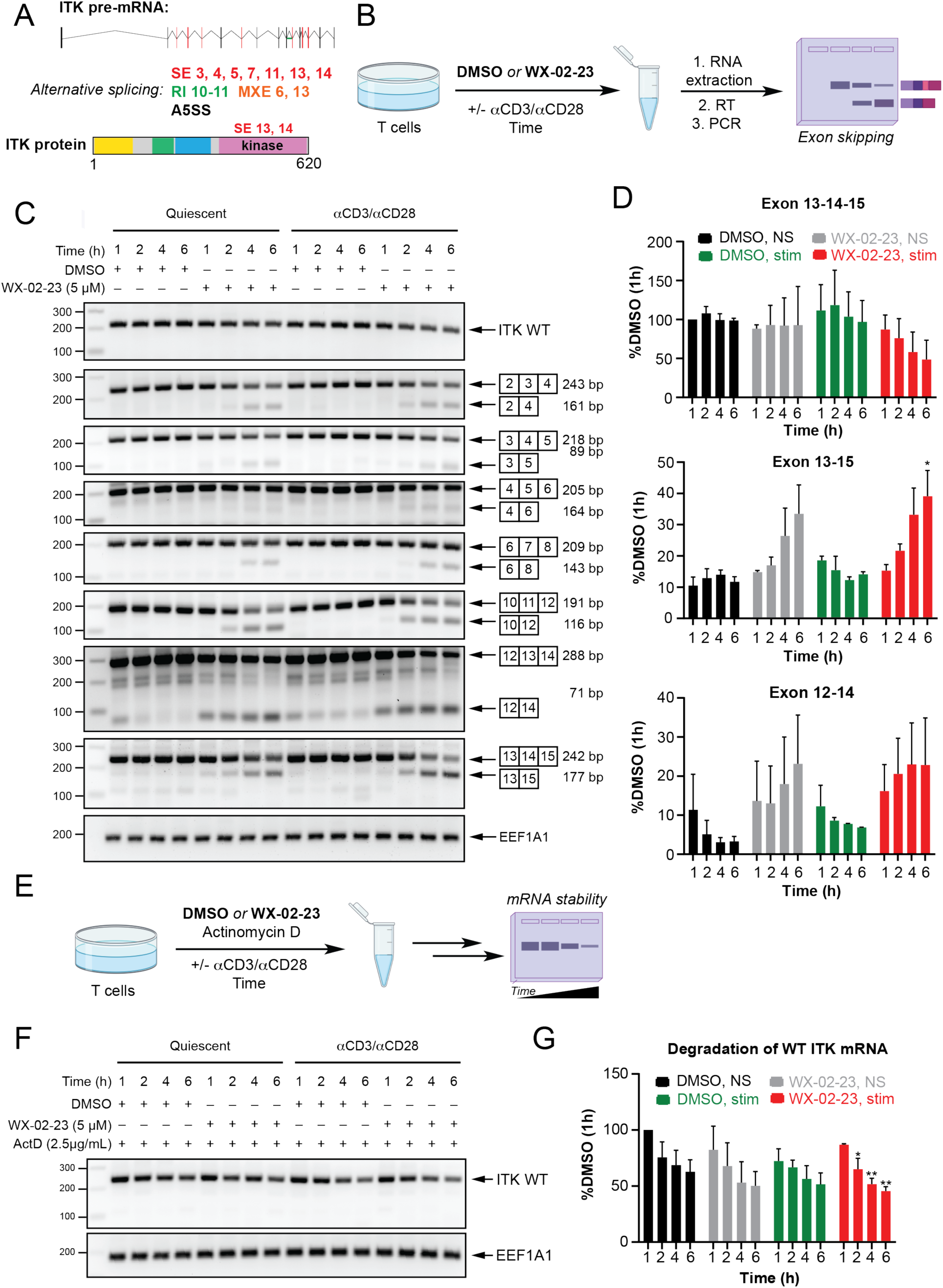
SF3B1 inhibitors induce significant alternative splicing events in ITK mRNA. (A) Schematic of ITK alternative splicing events identified from RNA-sequencing data. Alternative splicing events were considered significant with PSI > 0.2 and FDR < 0.05 identified by rMATs. Major exon skipping events were observed in the region corresponding to the kinase domain in the translated protein (exons 13, 14). (B) Workflow for evaluation of ITK splicing changes by RT-PCR. Primary human T cells were treated with DMSO or WX-02-23 (5 µM) for 1-6 hours with or without stimulation with αCD3/αCD28 antibodies, RNA was extracted and reverse transcribed, and PCR was used to generate amplified fragments using primers designed to span exons that were observed to be spliced out by RNA-sequencing. (C) RT-PCR analysis of alternative splicing of ITK mRNA induced by WX-02-23 in quiescent T cells versus stimulated T cells. Total RNA was isolated from quiescent and stimulated T cells treated with DMSO or WX-02-23 (5 µM) for the indicated times (1, 2, 4 or 6 h) and reverse transcribed to cDNA. PCR primers were designed to detect exon skipping events of exons 3, 4, 5, 7, 11, 13 and 14 identified from RNA-seq data. RT-PCR was performed with primers that bind to exons on both sides of the target exon. PCR products were separated by agarose gel (2%) electrophoresis and visualized by ethidium bromide staining. Eukaryotic translation elongation factor 1 alpha 1 (EEF1A1) served as a loading control. (D) Bar graphs showing changes in ITK exons 13 and 14 levels following compound treatment. Data are presented as the mean percentage of DMSO-treated quiescent cells (1 h) ± SD; n = 2/group. *, p < 0.05 compared to corresponding 1 h treatment using Dunnett’s multiple comparisons test (E-F) Effect of WX-02-23 on ITK mRNA stability in quiescent T cells versus stimulated T cells. (E) Workflow for evaluation of ITK mRNA stability by RT-PCR. (F) The decay rates of ITK mRNA were assessed using RT-PCR in quiescent and stimulated T cells treated with DMSO or WX-02-23 (5 µM) following transcription inhibition by actinomycin D (Act D, 2.5 µg/mL) for the indicated times (1, 2, 4 and 6 h). Total RNA was isolated, RT-PCR analysis was performed, and PCR products were visualized by ethidium bromide staining. Eukaryotic translation elongation factor 1 alpha 1 (EEF1A1) served as a loading control. (G) Bar graph showing degradation of WT ITK mRNA following compound treatment. Data are presented as the mean percentage of DMSO-treated quiescent cells (1 h) ± SD; n = 2/group. Average values from 2 biological replicates are shown. *, p < 0.05; **, p < 0.01 compared to corresponding 1 h treatment using Dunnett’s multiple comparisons test.

### WX-02-23 and PladB induce ITK mRNA alternative splicing

RNA-sequencing results suggested that ITK mRNA is alternatively spliced following compound treatment with the most substantial splicing events observed in the kinase domain of the protein (Fig. 4A, Supplementary Tables 2S-2). To confirm exon skipping events using a complementary approach, we performed standard PCR of reverse transcribed RNA from T cells treated with WX-02-23 with or without αCD3/αCD28 stimulation for 1-6 h (Fig. 4B-4D). We designed primers beginning on exons adjacent to each exon skipping event observed in the RNA-seq study (e.g., primers starting on exons 13 and 15 for the splicing event Δexon14) and used them for visualization of exon skipping by standard PCR. Indeed, we could see alternative splicing events as early as 1 h of compound treatment in both quiescent and activated T cells and a corresponding gradual decrease of WT mRNA (Fig. 4C, S4A, S4B). Skipping of exons 11, 13, and 14 were the most significant and early exon skipping events observed, in line with the RNA-seq experiment. However, no cell state-dependent differences in the kinetics of the formation of corresponding alternatively spliced products were observed, suggesting that the state-dependent loss of ITK in activated T cells is unlikely to be exclusively caused by alternative splicing (Fig. 4D).

Upon TCR stimulation, T cells undergo a major shift in RNA splicing, translation, and degradation[30]. To test whether the state-dependent loss of ITK upon WX-02-23 treatment is due to altered RNA metabolism in activated T cells (e.g., higher rates of RNA degradation in activated cells), we investigated the half-life of ITK mRNA by inhibiting DNA transcription with Actinomycin D (ActD). Treatment of T cells with ActD for 1-6 h (Fig. 4E-4G, S4C) led to a time-dependent decrease in ITK mRNA levels, but this decrease was not further affected by αCD3/αCD28 stimulation or WX-02-23 treatment, suggesting an alternative mechanism underpinning the impact of spliceosome modulatory compounds on ITK protein in stimulated T cells.

### ITK protein turnover is faster in activated T cells

Having observed similar ITK mRNA turnover rates and alternative splicing in WX-02-23-treated quiescent and stimulated T cells, we hypothesized that loss of ITK protein specifically observed in WX-02-23-treated stimulated T cells might stem from the differences in protein turnover rates. Indeed, treatment of T cells with DMSO with or without αCD3/αCD28 stimulation in the presence of a translation inhibitor cycloheximide (CHX) confirmed that ITK is more rapidly degraded in activated T cells (Fig. 5A-5C), possibly due to increased interaction with E3 ligases that regulate ITK expression[31, 32] and in line with a previous study using pulse-chase experiments to determine the half-life of ITK in CD4^+^ T cells[26]. Notably, pre-treatment with PF led to a full rescue of ITK degradation in stimulated T cells (Fig. 5B, C), consistent with this compound’s effects on blocking EV96-induced ITK loss[6]. While it remains mechanistically unclear how the binding of PF protects ITK from stimulation-induced degradation and extends the half-life of ITK under T cell activation conditions (Fig. 5B, 5C, S5A), our findings are in line with previous observations by Zapf and co-workers [26].

**Figure 5.**
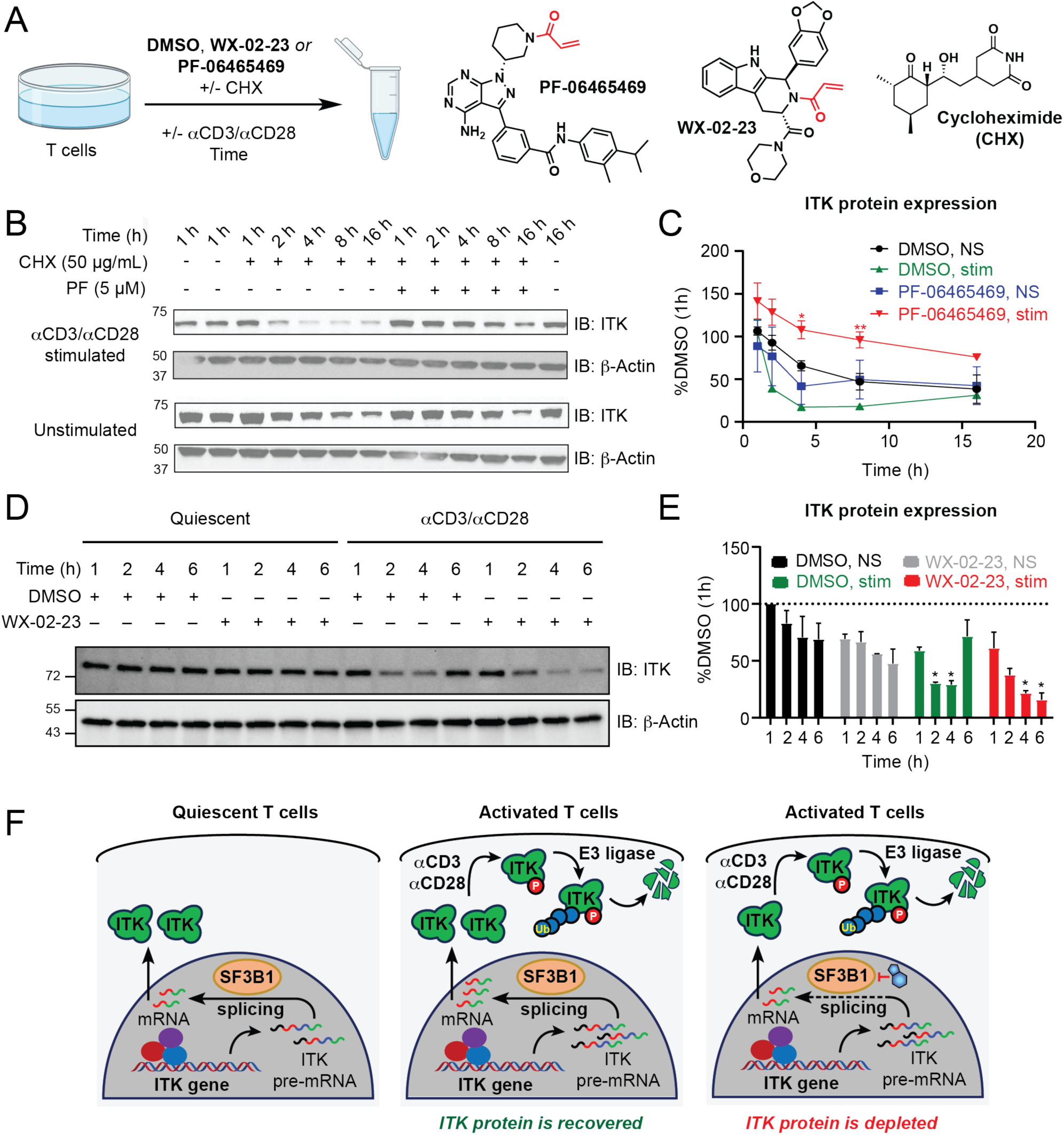
ITK stability and state-dependent reduction following SF3B1 inhibition. (A) Workflow for evaluation of time-dependent ITK protein expression in quiescent versus stimulated T cells, and the effect of PF-06465469 on ITK protein stability following translation inhibition by cycloheximide (50 µg/mL). (B and C) The decay rates of ITK protein were assessed using Western blot in quiescent and stimulated T cells treated with DMSO or PF-06465469 (5 µM) following translation inhibition by cycloheximide (50 µg/mL) for the indicated times (1, 2, 4, 8, and 16 h). Data are presented as the mean percentage of DMSO-treated quiescent cells (1 h) ± SEM; n = 3/group. *, p < 0.05, **, p < 0.01 PF-06465469 compared to DMSO treatment in stimulated T cells using Tukey’s multiple comparisons test. (D and E) Effect of T cell stimulation and WX-02-23 treatment on ITK protein expression. ITK protein expression levels were assessed using Western blot in quiescent T cells and stimulated T cells treated with DMSO or WX-02-23 (5 µM) for the indicated times (1, 2, 4 and 6 h). Data are presented as mean percentage of DMSO-treated quiescent cells (1 h) ± SD; n = 2/group. *, p < 0.05 compared to corresponding 1h treatment using Dunnett’s multiple comparisons test. (F) Schematic showing a plausible mechanism for state-dependency of ITK downregulation following inhibition of SF3B1 splicing factor in T cells. T cell activation leads to ITK phosphorylation and global structural re-arrangements, which can affect E3-ligase recruitment and protein turnover[2]. PF-06465469 locks ITK in its inactive conformation, thereby extending ITK half-life and leading to the observed “rescue” in Western blot experiments. Inhibition of SF3B1 leads to multiple alternative splicing events in ITK mRNA and reduction of functional ITK mRNA transcripts.

Considering the faster ITK turnover rates observed in experiments using CHX, we tested whether αCD3/αCD28 stimulation led to a decrease in ITK protein at shorter treatment times without the addition of translation inhibitor. Indeed, αCD3/αCD28 stimulation for 2 and 4 h led to a significant decrease in ITK levels, which was restored at longer treatment times (Fig. 5D, 5E), suggesting that there is a compensatory mechanism that replenishes the larger pools of depleted ITK under TCR stimulation conditions. Co-treatment with WX-02-23 under stimulating conditions led to a similar decrease in ITK levels at shorter timepoints, which was not restored at longer timepoints, likely due to depletion of WT mRNA pools, which can be used for replenishment of the degraded ITK (Fig 5D-5F, S5B).

To further test whether there were any cell state-dependent differences in the effect of WX-02-23 and PladB on the SF3B1 interacting partners, we performed immunoprecipitation-mass spectrometry (IP-MS) experiments using T cells treated with DMSO, WX-02-23, WX-02-43 or PladB in the presence or absence of αCD3/αCD28 stimulation (Fig. S5C, Supplementary Table S6). Similar to recent findings in 22RV1 cancer cells[14], WX-02-23 perturbed SF3B1 interactions with a number of proteins, including GPATCH11 and DHX15, while stabilizing interactions with another selection of proteins (e.g., DDX42 and DNAJ8C). WX-02-23 generally showed a similar profile to PladB with several notable exceptions, including decreases in HTATSF1 and DDX46, which further recapitulated the interaction profiles observed in cancer cells. However, no T cell state-dependent changes in WX-02-23 modified SF3B1 interactomes were observed.

### Ligandability of splicing factors and splicing regulators

The splicing factor SF3B1 has emerged as a key vulnerability in many cancers, including hematologic malignancies[33]. Identification of Pladienolide family of natural products with antitumor activity as inhibitors of SF3B1[28, 34] has enabled extensive studies into its role in cancer progression and nominated it as a target of interest for therapeutic development, leading to the synthesis of more advanced natural product analogues that are currently under clinical investigation[35]. Mutations in SF3B1 have also been associated with activation of proinflammatory signaling in myelodysplastic syndromes[36] and inhibition of SF3B1, or four other key components of the spliceosome (SF3A1, SF3A2, SF3A3, and EFTUD2) has been shown to reduce inflammatory cytokine production in LPS stimulated macrophages[37, 38]. Our study has revealed an unexpected mechanism, by which targeting of SF3B1 in T cells leads to suppression of T cell activation and a selective state-dependent downregulation of ITK protein due to alterations in the balance of production and degradation of ITK transcript and protein. Our findings thus point to an alternative therapeutic strategy for targeting T cells and T cell cancers that is complementary to existing mechanisms, such as direct ITK inhibitors or degraders[39]. Similar to protein degradation using bifunctional recruiters and molecular glues, perturbation of ITK splicing and the resulting impact on protein expression should address both the catalytic and scaffolding functions of the kinase[40–42], which are both important for suppression of T cell responses in T cell malignancies (Fig. 6A). This intriguing observation, together with the recent identification of small molecules targeting additional splicing factors/regulators, such as the spinal muscular atrophy drug risdiplam[43], covalent modulators of the RNA-binding protein NONO[7], and small-molecule protein degraders of RBM39[44], motivated us to perform a more in-depth study of splicing factors and splicing regulators that can be targeted with small-molecule electrophiles in the context of therapeutic discovery for cancer treatment and immune modulation[17, 22, 45–48]. To address this question, we first compiled an exhaustive list of proteins involved in splicing (Fig. 6B-6D, Supplementary Table S7)[49, 50]. Using CORUM[51] and Uniprot[52] databases, we curated a comprehensive list of proteins belonging to the spliceosome, known splicing factors, and splicing regulators, including protein kinases, phosphatases, mRNA polyadenylation enzymes, and methyl transferases (Fig. 6C). We further included RNA-binding prediction scores calculated in RBP2GO[53], as well as functional domain annotation from InterPro to facilitate identification of liganded cysteines in functionally relevant domains[54]. Of particular interest were ligandable cysteines annotated to be in short intrinsically disordered regions (IDRs), as a recent study by Mann and co-workers suggested that these domains have an increased rate of regulatory phosphorylation[55].

**Figure 6.**
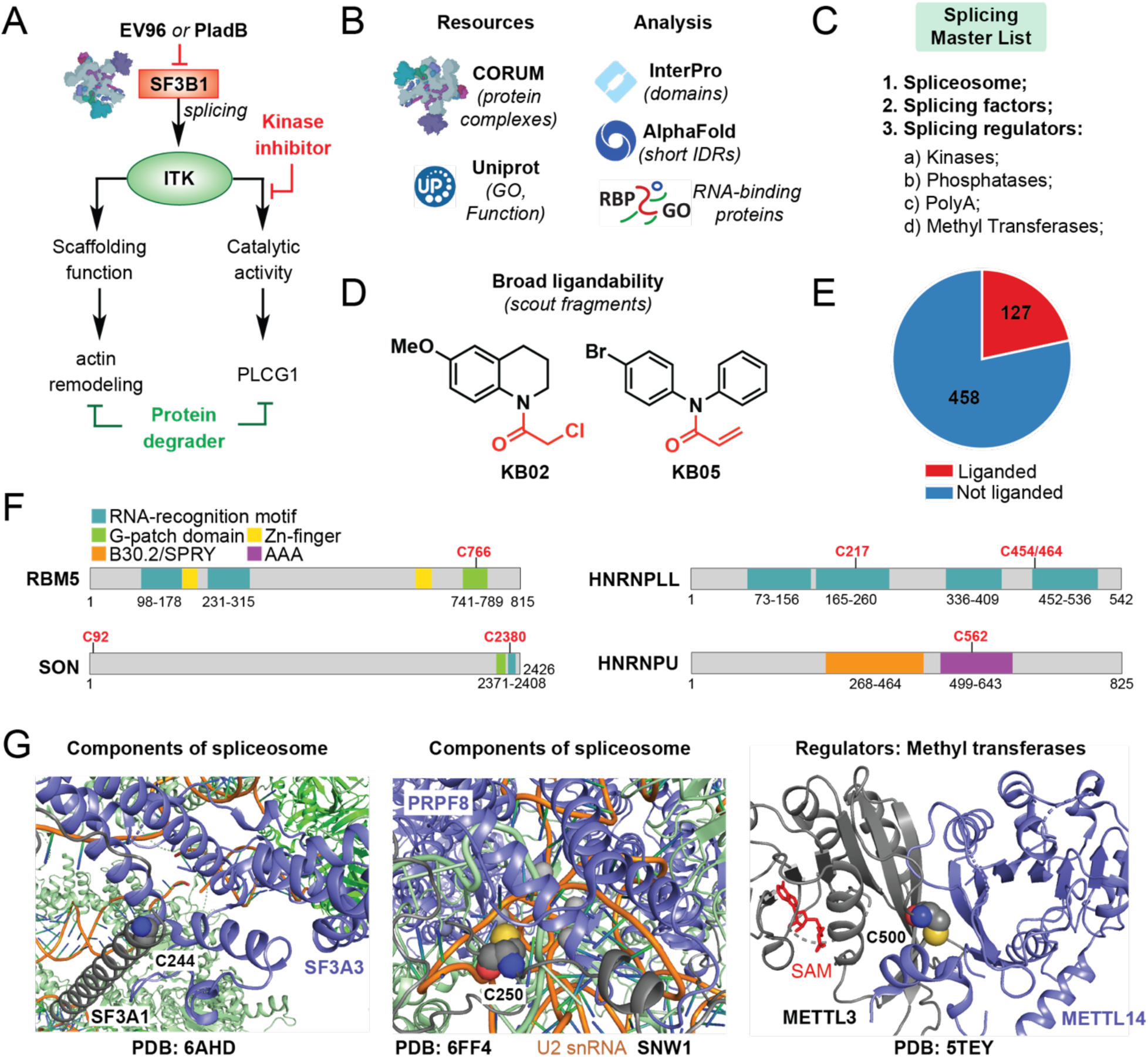
Generation of a splicing master list for identification of ligandable vulnerabilities in splicing factors and splicing regulators. (A) Conceptual differences between active site inhibitors, protein degraders, and post-transcriptional regulators of protein expression. Splicing regulators have the potential to offer new points of pharmacologic intervention, distinct from existing therapeutic modalities (B-D) Generation of the splicing factor master list. (B) Uniprot and CORUM databases were used for generation of a list of splicing factors, splicing regulators, and proteins present in associated complexes. The resulting shortlist of proteins was annotated using published data, including functional protein domains (InterPro), presence of short intrinsically disordered domains (IDRs[3]), and RNA-binding prediction (RBP2GO)[4]. (C) The resulting master list contains information on components of the spliceosome, annotated splicing factors, and splicing regulators, including kinases, phosphatases, polyadenylation factors, and methyl-transferases with reported roles in splicing. (D) Cysteine-directed TMT-ABPP broad ligandability data with scout fragments KB02 and KB05 in primary human T cells[1] was further integrated to identify cysteine residues amenable to covalent modification and ligand development. (E) Pie chart representation showing splicing master list proteins containing cysteine residues liganded by scout fragments, identified in cysteine-directed TMT-ABPP experiments. (F) Protein sequences of targets from the splicing master list identified using cysteine-directed TMT- ABPP platforms. Cysteine residues are considered “liganded”, if scout fragments KB02 or KB05 show >75% competition for reactivity of that site. Specifically, liganded sites were identified in RNA-recognition motifs (SON_C2380, HNRNPLL_C217, HNRNPLL_C454/464), G-patch domains (RBM5_C766), and AAA-domains (HNRNPU_C562). (G) Cryo-EM and crystal structures of representative splicing factor and splicing regulator targets of scout fragments identified using cysteine-directed TMT-ABPP platforms. Proteins of interest are shown in grey and ligandable cysteine residues are shown as spheres colored by elements. Left: Cryo-EM structure of human pre-catalytic spliceosome (B complex) (PDB: 6AHD) with ligandable cysteine in splicing factor SF3A1_C244 located at the protein-protein interaction surface with SF3A3; Middle: Cryo-EM Structure of human Bact spliceosome core (PDB: 6FF4) with ligandable cysteine in SNW1_C250 located at the protein-RNA and protein-protein (PRPF8) interface; Right: Crystal structure of human METTL3-METTL14 complex (PDB: 5TEY), with ligandable cysteine in **N6-adenosine-methyltransferase catalytic subunit** METTL3_C500 located at the protein-protein interaction surface with METTL14.

To complete the master list, we incorporated broad ligandability data that we recently collected using cysteine-directed TMT-ABPP mass spectrometry workflows with “scout” fragments KB02 and KB05 in activated and quiescent state T cells (Fig. 6D)[6]. “Scout” fragments are broadly reactive electrophiles that contain both an affinity element and a reactive group, which enables their use for broad profiling of sites in proteins that can be targeted with small-molecule electrophiles, making them useful tools for target and ligand discovery efforts[6, 56, 57].

The resulting list contained over 550 proteins with relevance to splicing, with over 100 proteins having one or more ligandable sites (Fig. 6E; see Supplementary Table S7 and Methods). Domain annotation using InterPro identified proteins with ligandable sites in functionally relevant domains (Fig 6F, S6A), including GPATCH domains and RNA recognition motifs (e.g., RBM5_C766, SON_C2380, HNRNPLL_C217,C454/C464), AAA-domains (e.g., HNRNPU_C562 and HNRNPUL1_C487), as well as IDRs adjacent to RRM domains (e.g., ESRP2_C237), suggesting the potential of targeting these sites for functional regulation of these proteins.

Heterogeneous nuclear ribonucleoprotein LL (HNRNPLL) is known to participate in CD44 and CD45 splicing, which are crucial for B cell to plasma cell differentiation and T cell activation, respectively[58, 59]. We identified two ligandable sites in HNRNPLL, located in two distinct RNA-binding domains, representing unique opportunities for targeting of distinct splicing events regulated by this splicing factor. Heterogeneous nuclear ribonucleoprotein U (hnRNPU) is similarly required for T cell activation and is involved in MALT1 alternative splicing[60]. Liganded cysteines in HNRNPU and its close homologue HNRNPUL1 are conserved and located in the AAA domain of these proteins. AAA domain of HNRNPU has been shown to regulate its oligomerization and play a role in chromatin decompaction[61], offering a starting point for developing chemical probes to study HNRNPU/HNRNPUL1 AAA function.

Another example includes RNA-binding protein NONO, with a ligandable site C145 located between two RNA-binding domains (Fig. S6A, S6B). Together with SFPQ and PSPC1, NONO is one of the three structurally similar members of the DBHS protein family, which play a role in a wide range of protein-protein and protein-nucleic acid interactions, regulating cellular processes ranging from transcriptional regulation, RNA processing and DNA repair[62]. Selective covalent electrophile ligands targeting NONO_C145 have been recently developed and shown to stabilize NONO interaction with mRNA, leading to unique “trapping” pharmacology that depends on the presence of NONO, but is not recapitulated by genetic disruption of NONO[7].

Further analysis of available crystal structures revealed that many of the ligandable sites in both splicing factors and splicing regulators are located at protein-protein or protein-nucleic acid interaction surfaces (Fig. 6G, S6B-S6E), including ligandable sites in known components of the spliceosome (SF3A1_C244, SNW1_C250; Fig. 6G), splicing factors (NONO_C145; Fig. S6B), and splicing regulators of mRNA methylation (METTL3_C500, METTL16_C253; Fig. 6G, S6C), protein phosphorylation (CDK12_C1009; Fig. S6D), and mRNA polyadenylation (CPSF4_C41; Fig. S6E). A number of these proteins have been shown to play key roles in acute and chronic inflammation and could represent unique opportunities for treatment of inflammatory disorders. For example, splicing factor 3 subunit 1 (SF3A1) is a splicing factor that participates in the assembly of 17S U2 snRNP and is responsible for regulating toll-like receptor (TLR) signaling[63]. The Ski-interacting protein (SNW1) has been shown to play a role as a transcriptional co-activator of NF-κB pathway[64] and the m^6^A writer protein METTL3 (a methyltransferase, or MTase) regulates T cell differentiation and homeostasis[65]. We detected C500 as a ligandable site in the METTL3 MT-A70 domain, which is located at the protein dimer interface with METTL14, known to bind substrate RNA[66], suggesting that targeting of this site could lead to functional outputs with modified m^6^A landscapes. Similarly, cleavage and polyadenylation specificity subunit CPSF4 was liganded at C41, which occurs within a zinc-finger region and may disrupt interaction between CPSF4 and CPSF1[67], affecting the polyadenylation landscapes (Fig. S6E).

### Scout fragment competition experiments with alkyne probes reveal high occupancy liganding sites

Cysteine-directed TMT-ABPP experiments (Fig. 7A, top) provide proteome-wide site-of-labeling information for covalent ligands without requiring chemical modifications to the ligands. However, as our current study, as well as other recent work[14, 25], highlighted, some ligandable cysteines may reside on non-proteotypic peptides and evade detection by cysteine-directed ABPP. Protein-directed ABPP experiments, where alkyne analogues of ligands are used to directly enrich protein targets, offer a complementary approach that exchanges knowledge of site-of-labeling for the quantification of many other (unlabeled) peptides in probe-modified proteins (Fig. 7A-7C, Supplementary Table S8)[14, 25]. We therefore decided to perform global evaluation of targets that can be both liganded and efficiently competed with scout fragments and corresponding scout fragment-based chemical probes. We envisioned that this approach would allow us to further advance our knowledge on the versatility of scout fragments and scout fragment-based chemical probes in drug discovery. Specifically, we sought to (1) uncover targets that might be overlooked by cysteine-directed TMT-ABPP; (2) validate direct target engagement with scout fragments; and (3) identify targets with high stoichiometry engagement, which can be nominated for future streamlined drug discovery efforts using scout fragment based chemical probes.

**Figure 7.**
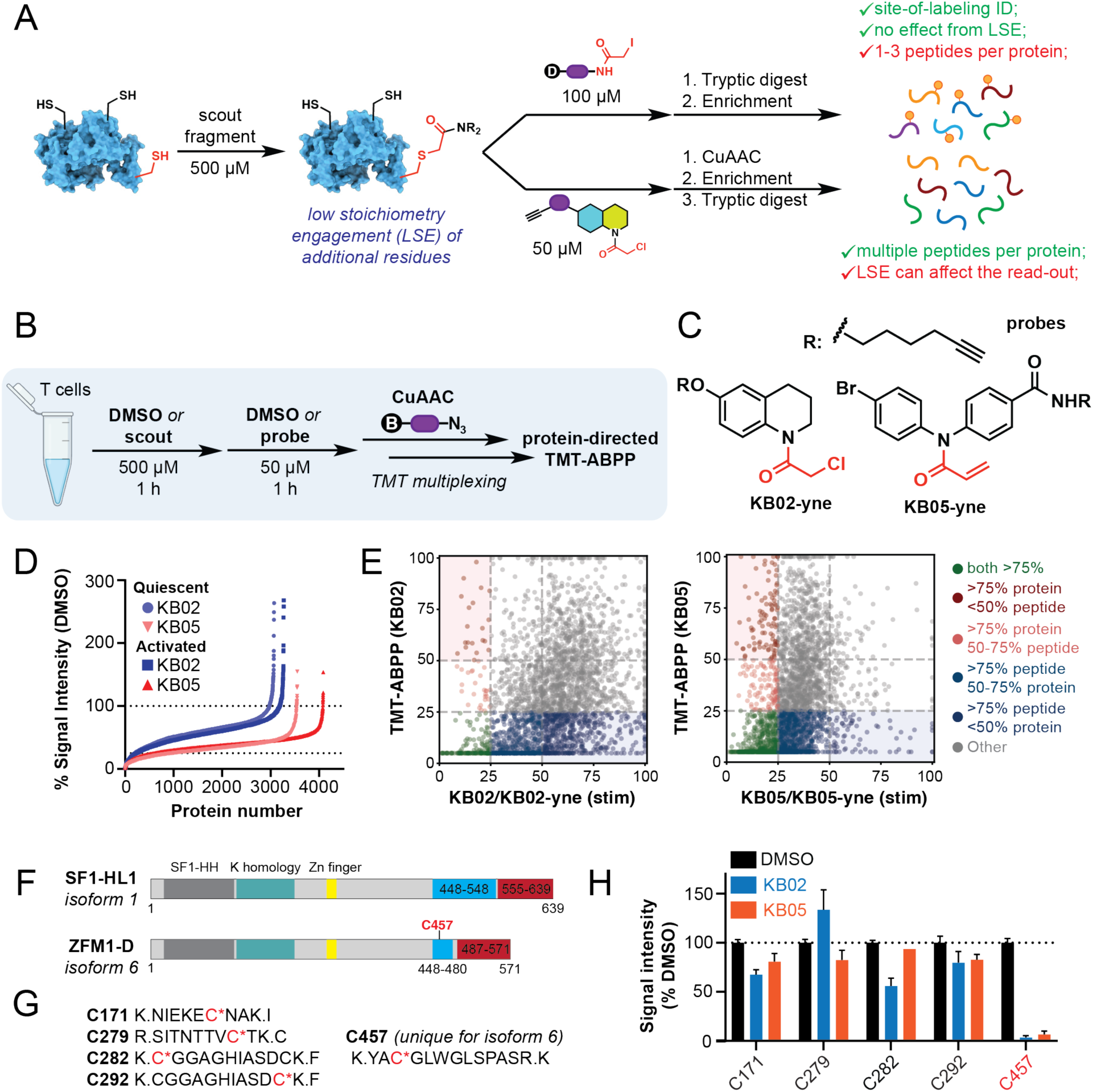
Protein-directed TMT-ABPP using scout fragment-based chemical probes. (A) Schematic showing the differences between cysteine-directed (top) and protein-directed (bottom) TMT-ABPP. Benefits and drawbacks of each of the complementary methods are listed. (B) Experimental workflow for quantitative proteomic experiments evaluating competition with scout fragments (KB02 and KB05) for cysteine reactivity against scout fragment-based alkyne containing chemical probes (KB02-yne and KB05-yne). Specifically, (1) Cell lysates from quiescent and αCD3/αCD28 stimulated T cells were treated with scout fragments (KB02 (500 µM), KB05 (500 µM)) or DMSO control for 1 h at room temperature, followed by scout fragment-based alkyne probes KB02-yne or KB05-yne (50 µM, 1 h); (2) Desthiobiotin enrichment handle was introduced using a copper-catalyzed azide-alkyne cycloaddition (CuAAC) reaction with biotin-azide chemical probe; and (3) resulting cell lysates were processed and analyzed by protein-directed TMT-multiplexed quantitative proteomics, where a 75% reduction in protein enrichment was interpreted as significant engagement and competition. See supplementary information for more details. (C) Structures of scout fragment-based alkyne containing chemical probes KB02-yne and KB05-yne. (D) Waterfall plot representation of competition ratios in the protein-directed TMT-ABPP competition experiments using KB02/KB02-yne and KB05/KB05-yne scout fragment and scout fragment-based alkyne probe pairs in quiescent and stimulated T cells. (E) Scatter plot representation showing comparison between cysteine-directed TMT-ABPP ligandability values[1] and protein-directed TMT-ABPP competition values. Proteins uniformly showing >75% ligandability and competition are nominated as highest priority targets for follow up studies and ligand development. Proteins uniquely liganded in cysteine-directed TMT-ABPP data are shown in blue. Proteins uniquely competed in protein-directed TMT-ABPP competition data are shown in red. (F-H) Protein-directed TMT-ABPP competition experiments reveal a previously overlooked isoform-specific ligandable cysteine in splicing factor SF1. (F) Schematic representation of sequence differences between isoform 1 (canonical) and isoform 6 (non-canonical) of SF1. (G) Left: Quantified cysteine-containing peptides belonging to both canonical and non-canonical SF1 isoforms; Right: Unique peptide belonging to non-canonical SF1 isoform – isoform 6 (ZFM1-D). (H) Bar graph showing signal intensities for quantified cysteine-containing peptides in SF1 isoforms 1 and 6 following treatment with DMSO (control), KB02 and KB05 (scout fragments). C457 is only present in isoform 6 and is engaged by both scout fragments.

We initiated the study by performing protein-directed TMT-ABPP competition experiments using KB02 and KB05 scout fragments at concentrations used in established *in vitro* broad ligandability studies (500 µM)[6, 56, 57] and alkyne-containing chemical probes KB02-yne and KB05-yne (50 µM, Fig. 7B-7C)[68]. We chose to test this chemical proteomic platform in T cells to compare this data with our previous results obtained using cysteine-directed TMT-ABPP experiments[6]. KB02 and KB05 scout fragments have been previously shown to be equally reactive in both cancer and immune proteomes[6, 56, 57], revealing ligandability (>75-80% target engagement) of 15-20% of identified sites. To our surprise, a much smaller portion of proteins enriched with KB02-yne probe was significantly competed with KB02 (>75%, 198 proteins in expanded and 228 in activated T cells), when compared to the competition levels with KB05 and KB05-yne (594 proteins in expanded and 955 in activated T cells, Fig. 7D). This stark difference suggests that low stoichiometry interactions with protein binding pockets outside of the main sites of ligandability identified using cysteine-directed TMT-ABPP approaches are likely more prevalent for smaller KB02 ligand compared to KB05 with larger surface area. Another interesting feature of protein-directed TMT-ABPP competition experiments with KB02 was a reproducible increase in protein enrichment for a number of targets, suggesting potential allosteric modulation that leads to enhanced KB02-yne probe binding, which has been previously observed with other covalent ligands.

To investigate the correlation between the new protein-directed TMT-ABPP data with scout fragments and scout fragment-based chemical probes and the cysteine-directed TMT-ABPP[6], we calculated the maximum competition value from all the quantified peptides corresponding to the protein of interest from our previous broad ligandability study in T cells and compared it to the protein-directed TMT-ABPP competition values (Fig. 7E, S7A, S7B). We identified proteins with high levels of competition measured using both chemical proteomic platforms, including XPO1, splicing factors NONO and HNRNPLL, as well as DNA and RNA binding protein SON (Fig. 7E, green dots). Most of these proteins contain one or two high stoichiometry labeling events and can be prioritized for future ligand discovery and functional follow up studies. We also identified proteins, which showed competition in the original peptide-level TMT-ABPP experiments, but lack of competition with scout fragment-based chemical probes, especially in the case of KB02/KB02-yne pair (Figure 7E, blue). One striking example was splicing factor HNRNPU, which was not competed by either of the scout fragments in the current protein-directed TMT-ABPP study. These examples suggest that there might be multiple low-stoichiometry engagement events in some proteins that can prevent the use of alkynylated scout fragment probes as tools for site-specific follow up studies.

### Analysis of scout fragment competition with alkyne probes reveals new ligandable sites, including sites with isoform-specific engagement

We were encouraged to see new ligandable proteins identified using protein-directed TMT-ABPP competition experiments (Fig. 7E, red dots), suggesting existence of previously unannotated ligandable sites in these proteins that were not detected by cysteine-directed ABPP (see Supplementary Data). One protein that caught our attention was splicing factor 1 (SF1), which showed competition in KB02/KB02-yne and KB05/KB05-yne experiments, but not in the previous cysteine-directed ABPP experiments despite all four cysteine-containing peptides in this protein (C171, C279, C282, and C292, Fig. 7F-7H) being quantified. SF1 is a key splicing factor involved in early stages of spliceosome assembly. It is known to mediate splice site recognition[69, 70] and mutations in the corresponding gene have been linked to tumorigenesis[71]. There are 10 potential SF1 isoforms, including the canonical SF1-HL, however contributions of each specific isoform to spliceosome assembly and splice site recognition have not been studied. Robust competition observed in our protein-directed TMT-ABPP data suggested that it is possible that we are reading out ligand engagement of a cysteine residue present in one of the non-canonical SF1 protein isoforms, which was not included in our original FASTA database for peptide mapping. We performed an additional search of our data to see, if any of the non-canonical isoforms containing cysteine residues contributed to the observed competition event. Indeed, we identified a cysteine (C457) uniquely found in isoform 6 of SF1 that was highly ligandable with both KB02 and KB05 (Fig. 7G-7H, Supplementary Table S9). Very little is known about functional differences between these isoforms and we expect identification of more advanced isoform-specific small-molecule inhibitors of this site will facilitate these studies.

## Discussion

Kinases are important signaling regulators that have both catalytic activity responsible for phosphorylation of substrates and scaffolding functions, which facilitate recruitment of substrates and larger complex assembly[40, 42]. There is a growing appreciation that both catalytic and scaffolding functions of kinases contribute to disease phenotypes, and, in such cases, inhibitors of these enzymes may fail to fully suppress pathology. Development of small-molecule protein degraders, including bifunctional recruiters and molecular glues, has been shown to overcome this limitation of enzymatic inhibitors. Indeed, a recent study by Wu and Gray labs has shown that bifunctional ITK kinase degraders can overcome therapeutic resistance in T cell lymphomas[39]. While the field of protein degradation has exploded over the last 10 years, one can envision development of alternative strategies affecting protein homeostasis, including the much less explored targeting of splicing with small molecules.

Identification of small-molecule chemical probes with state-dependent pharmacology[72] has great potential for the development of inhibitors that can preferentially target pathologic over physiologic functions of immune responses. Our serendipitous discovery of an electrophilic small molecule that leads to stereoselective, state-dependent loss of ITK through splicing inhibition underscores an underappreciated mechanism for achieving state-dependent pharmacology. Our extensive mechanistic investigations revealed that EV96 induces loss of ITK through inhibition of a major splicing factor SF3B1 and that the observed state-dependency is likely due to a combination of faster ITK protein turnover rates in activated vs quiescent T cells and depleted pools of functional mRNAs due to alternative splicing. The observed selectivity in regulation of protein homeostasis points to a broader potential of targeting splicing factors in immune cells for therapeutic purposes. Indeed, previous work has shown that partial gene knockdown of several key spliceosome components, including SF3B1, SF3A1, SF3A2, SF3A3, and EFTUD2 in macrophages reduces TLR-induced activation, but has no effect on macrophage viability or phagocytosis[37, 38], suggesting that there is underexplored potential of targeting proinflammatory pathways with splicing inhibitors without pronounced effects on the physiologic functions of immune cells. Similarly, our work has shown that SF3B1 inhibition with either previously described natural products (PladB) or newer synthetic tryptoline acrylamides affects T cell activation and IL2 production, without inducing global protein expression changes. Future work will show whether the EV96 tryptoline scaffold – which represents, to our knowledge, the first fully synthetic small molecule chemotype for liganding SF3B1 – has utility as a treatment for myelodysplastic disorders or solid tumors, similar to natural product SF3B1 modulators that are under current clinical investigation. One major question to be addressed is whether the lack of apparent cytotoxicity will translate into similarly low toxicity in other cell types and animal models. Pharmacological modulation of splicing is an exciting area for treatment of genetic disorders[16], cancer[23], and infectious disease[73]. Despite ongoing clinical trials investigating inhibition of splicing in cancers, and the approval of the gain of function splicing modulator risdiplam for treating spinal muscular atrophy[43], our ability to selectively target splicing with small moleclues is in its infancy. To begin to address this question, we have compiled a comprehensive list of proteins involved in splicing, including splicing factors, components of spliceosome, and proteins involved in post-translational regulation of splicing (Fig. 6C). We further leveraged broad ligandability data using cysteine reactive “scout” fragments and cysteine-directed TMT-ABPP platforms to assess proteome-wide potential for targeting splicing in T cells with small-molecule covalent electrophiles. This approach highlighted over 100 targets that can be pursued for future ligand discovery and drug development purposes. We also showed that functional domain annotation can be used to prioritize target selection in a way that maximizes the opportunity to ligand a protein in a domain associated with a biological outcome.

Cysteine-directed and protein-directed TMT-ABPP platforms are complementary approaches for identification of proteins and sites on proteins that can be targeted with small-molecule covalent electrophiles. Cysteine-directed TMT-ABPP platforms use broadly reactive chemical probes and analysis of single peptides that were directly engaged with the probe. Protein-directed TMT-ABPP platforms involve enrichment of proteins that form irreversible covalent bonds with chemical probes of interest and analysis of all the peptides, excluding the peptide that reacted with the probe. Protein-directed TMT-ABPP is most commonly used with advanced inhibitors, where introduction of corresponding alkyne-containing chemical probes allows for extensive characterization of target landscapes, enabling mechanism-of-action studies and identification of potential off-targets. Indeed, using protein-directed TMT-ABPP experiments with WX-02-23 and EV96A-yne probe allowed us to identify several new stereospecific targets of EV96, which were overlooked using cysteine-directed ABPP, including splicing factor SF3B1. We have further shown that protein-directed TMT-ABPP platforms that leverage broadly reactive scout fragments and scout fragment-based chemical probes can also be useful for identification of new targets, which might be overlooked using standard cysteine-directed TMT-ABPP approaches, including targets with isoform-specific engagement, like SF1_C457, which could open new grounds for studies of isoform-specific contributions of proteins to cell biology. Although the ligandability of specific splicing isoforms depends on the occurrence of unique cysteines, this result is a good demonstration of the idea that ligandability profiling is uniquely able to identify such isoforms in a target agnostic manner.

Comparison of cysteine-directed and protein-directed TMT-ABPP data provides unique information on the potential for the use of scout fragments and scout fragment-based chemical probes for drug discovery by gel-based ABPP, “function-first” biochemical assays, and crystallography studies, where high stoichiometry and site-specificity of engagement are crucial. With the growing interest in leveraging “scout” fragments for broad ligandability mapping of different protein classes and emerging drug discovery efforts, we anticipate that our scout fragment competition data will serve as a useful resource for the chemical biology and chemical immunology community to inform target and chemical probe selection for ligand discovery and functional follow up studies.

### Limitations of the study

Targeting core splicing factors, including SF3B1, is a promising therapeutic strategy for hematologic malignancies, however it might encounter limitations due to potential safety concerns and limited therapeutic index. Alternative strategies can include targeting accessory splicing factors and post-translational regulators. Post-translational modifications of splicing factors are known to regulate many processes, ranging from their subcellular localization, protein-protein interactions required for spliceosome assembly and efficient splicing. Indeed, therapeutic approaches to cancers with dysregulated splicing have now expanded to include inhibitors of arginine methylation (PRMT5 and type I PRMT enzymes), as well as phosphorylation (CLK and SRPK1)[74], which have also been included into our master splicing list.

While we have shown here that protein-directed ABPP can facilitate the discovery of proteins liganded by electrophilic fragments that may have been overlooked in cysteine-directed ABPP experiments, several important factors can complicate the interpretation of protein-directed ABPP data, including multiple low stoichiometry reactivity events on a protein by the alkyne probe and differences in the proteomic interaction landscape of alkyne vs non-alkyne electrophilic compounds. Nonetheless, as we have also shown recently with more elaborated stereochemically defined electrophilic compounds[25], we believe that the integrated use of cysteine- and protein-directed ABPP in evaluating electrophilic scout fragments constitutes a useful approach for identifying actionable covalent ligand binding events across the proteome. We would also like to note that only one treatment condition was used in our scout fragment-based protein-directed TMT-ABPP competition experiments. It is possible that varying probe concentration, labeling time and temperature, can highlight additional sites and proteins that can be studied using scout fragment-based probes.

### Significance

There is a growing appreciation that alternative splicing can play distinct roles in immune cell pathophysiology[15, 75, 76]. However, our understanding of the existing opportunities for targeting of splicing in the context of immunopathologies is limited. Here we report for the first time a small-molecule covalent electrophile that inhibits T cell activation and induces a T cell state-dependent loss of an important immune kinase, ITK, by targeting a major splicing factor SF3B1. The lack of observed cytotoxicity or gross protein expression changes associated with inhibition of splicing suggests that targeting of splicing could be a viable therapeutic strategy in immune-related disorders. To further investigate this opportunity, we generated a comprehensive list of splicing factors and splicing regulators, which can be targeted with small molecule covalent electrophiles based on peptide-level and protein-level TMT-ABPP experiments with scout fragments. This approach produced a valuable community resource, aimed at facilitating future target follow up and ligand development efforts.

## Supporting information

Chemistry SI

## Acknowledgements

We thank Jared Ramsey for technical assistance. This work was supported by Rockefeller University start-up funds (E.V.V.), Robertson Foundation (E.V.V.), The Achelis and Bodman Foundation (E.V.V.), NIH (T32GM136640-Tan to T.L.Z; R35CA231991 to B.F.C.; R00CA248715 to X.Z.), and Kimberly Lawrence-Netter Postdoctoral Fellowship (K.A.S.). E.V.V. is supported by Searle Scholarship Program. We thank the Flow Cytometry Resource Center and Genomics Resource Center at The Rockefeller University for help with acquiring flow cytometry and RNA-sequencing data.

## Author Contributions

Conceptualization: C.D.W., K.A.S., and E.V.V.; data curation, H.K., N.R., S.H., T.L.Z., J.L., and E.V.V.; formal analysis, H.K., N.R., S.H., J.L., T.L.Z., K.A.S., and E.V.V.; investigation: H.K., C.D.W., T.L.Z., K.A.S., C.W., X.Z., J.R., and E.V.V.; methodology: H.K., C.D.W., K.A.S., and E.V.V.; resources: B.M., B.F.C., J.L., O.A.-W., and E.V.V.; software, N.R., and S.H.; supervision: E.V.V.; writing – original draft, C.D.W., K.A.S., and E.V.V.; writing – review & editing, B.F.C., O.A.-W., and E.V.V. All authors read, edited, and approved the final manuscript.

## Supplementary Information

### (A) Supplementary Figures

**Supplementary figure 1.**
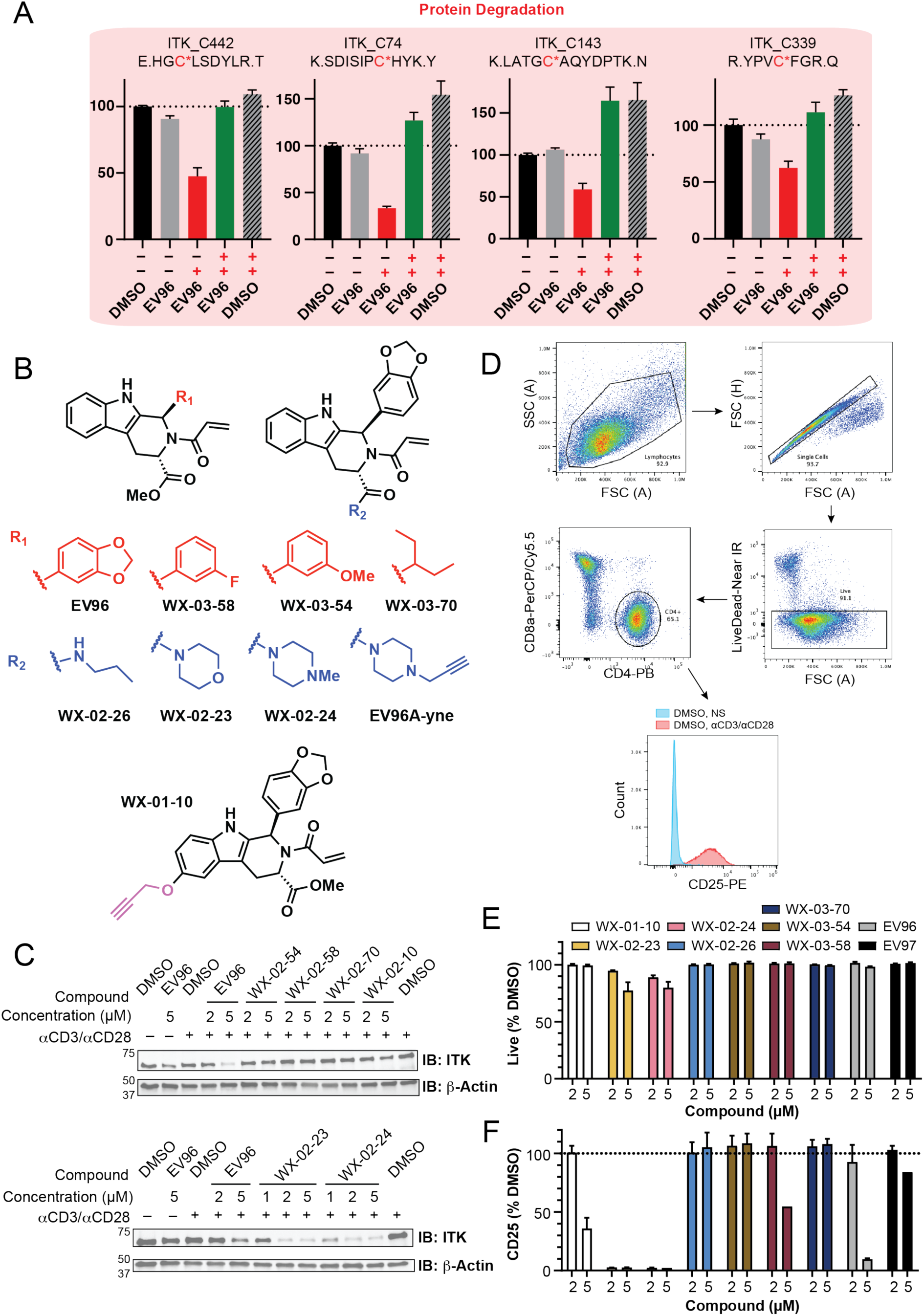
Evaluation of EV96 analogues for suppression of T cell activation and reduction of ITK levels. (A) *In situ* cell based evaluation of ITK_C442 engagement with EV96 using TMT-ABPP proteomic workflows with tandem trypsin/GluC protease digest. Bar graph representation of changes in signal intensities for all quantified ITK peptides, highlighting a clear loss of protein event. (B) Structures of EV96 and all EV96 analogues tested in the study. (C) Western blot analysis of ITK levels following T cell treatment with EV96 and EV96 analogues at 2 µM and 5 µM concentrations for 24 h. (D) The following FACS gating strategy was used for flow cytometry analysis of suppression of T cell activation using *(R,S)*-tryptoline analogues: (1) Lymphocyte selection by plotting FSC-A/FSC-A; (2) Doublet discrimination by plotting FSC-A/FSC-H; (3) Live cell selection by plotting SSC-A (y-axis) versus Live/Dead Near-IR (x-axis, logarithmic scale) and gating on the negative population; (4) Measurement of geometric mean of fluorescence intensity in the PE channel (PE-CD25 antibody). (E) Flow cytometry analysis of cell viability following treatment with EV96 and EV96-analogues at 2 µM and 5 µM concentrations for 24 h. (F) Flow cytometry analysis of CD25 levels following treatment with EV96 and EV96-analogues at 2 µM and 5 µM concentrations for 24 h. Little difference was observed in the effect of compound treatment on CD25 levels between CD4^+^ and CD8^+^ T cell subpopulations. As a result, pan-T cell CD25 values are reported for evaluation of compound efficiency.

**Supplementary figure 2.**
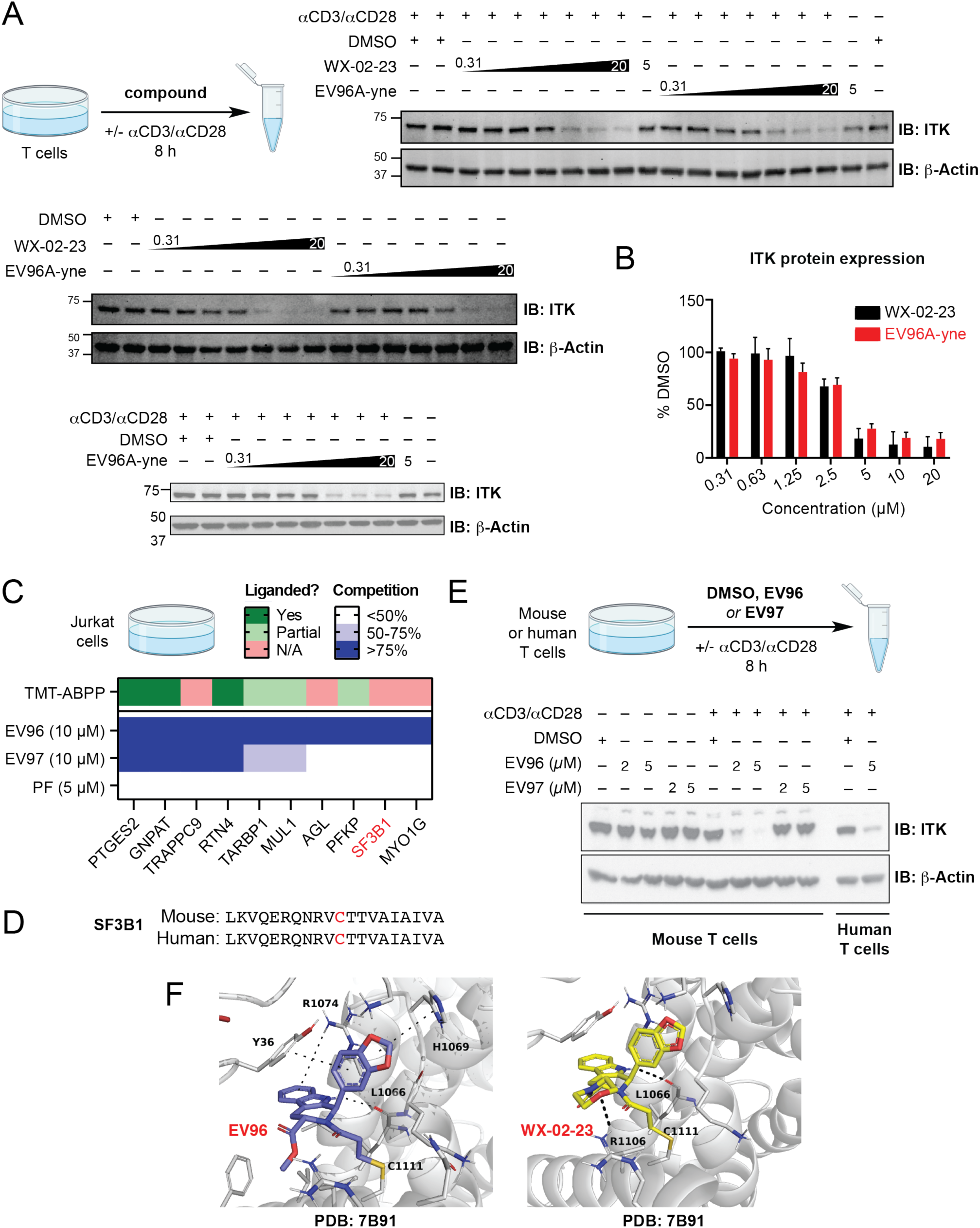
Identification of a potent chemical probe EV96A-yne for target ID studies. (A-B) Effect of EV96 analogues WX-02-23 and EV96A-yne on ITK levels in stimulated primary human T cells. (A) Western blot showing ITK levels following treatment with increasing concentrations of WX-02-23 and EV96A-yne for 8 h. (B) Bar graph showing mean percentage of DMSO-treated control cells ± SD following compound treatment for 8 h; n = 2-3/group. (C) Schematic of a protein-directed TMT-ABPP competition experiment in Jurkat cells using EV96A-yne chemical probe (5 µM, 2 h) following pre-treatment with DMSO, EV96 (10 µM), EV97 (10 µM), and PF-06465469 (PF, 5 µM) for 3 h. Heatmap showing percent pulldown competition following compound treatment and evidence of ligandability with EV96 in a previous cysteine-directed TMT-ABPP study. Data are average values from 2 independent experiments. (D) Alignment of human and mouse SF3B1 sequences showing conservation of C1111 between species[5]. (E) Treatment of both human and mouse primary T cells leads to cell state-dependent decrease in ITK levels. Western blot showing ITK levels following treatment with active compound EV96 (2 and 5 µM) and its inactive enantiomer EV97 (2 and 5 µM) for 8 h. (F) Docking model of active compounds EV96 and WX-02-23 covalently bound to SF3B1_C1111 (PDB: 7B91)[5]. Additional stabilizing interactions are highlighted using dashed lines.

**Supplementary figure 3.**
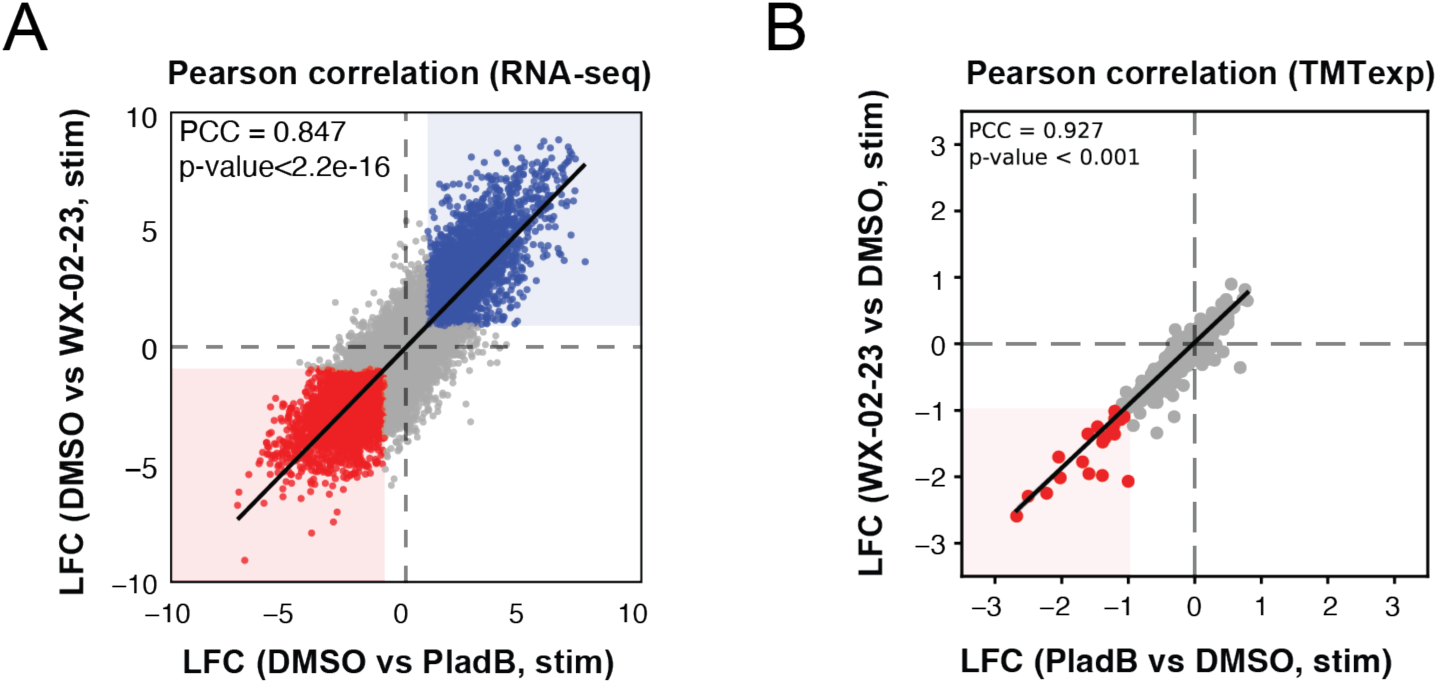
Comparison of global mRNA and protein expression changes following WX-02-23 and PladB treatment. (A-B) Scatter plots showing mRNA (A) and protein (B) abundance changes in stimulated T cells treated with WX-02-23 (5 µM) or PladB (5 µM) for 8 h. Data is from 3 biological replicates and is shown as average values of log_2_ fold changes. Pearson correlation coefficients (PCC > 0.75) suggest strong positive correlation of both mRNA and protein abundance changes induced by both compounds.

**Supplementary figure 4.**
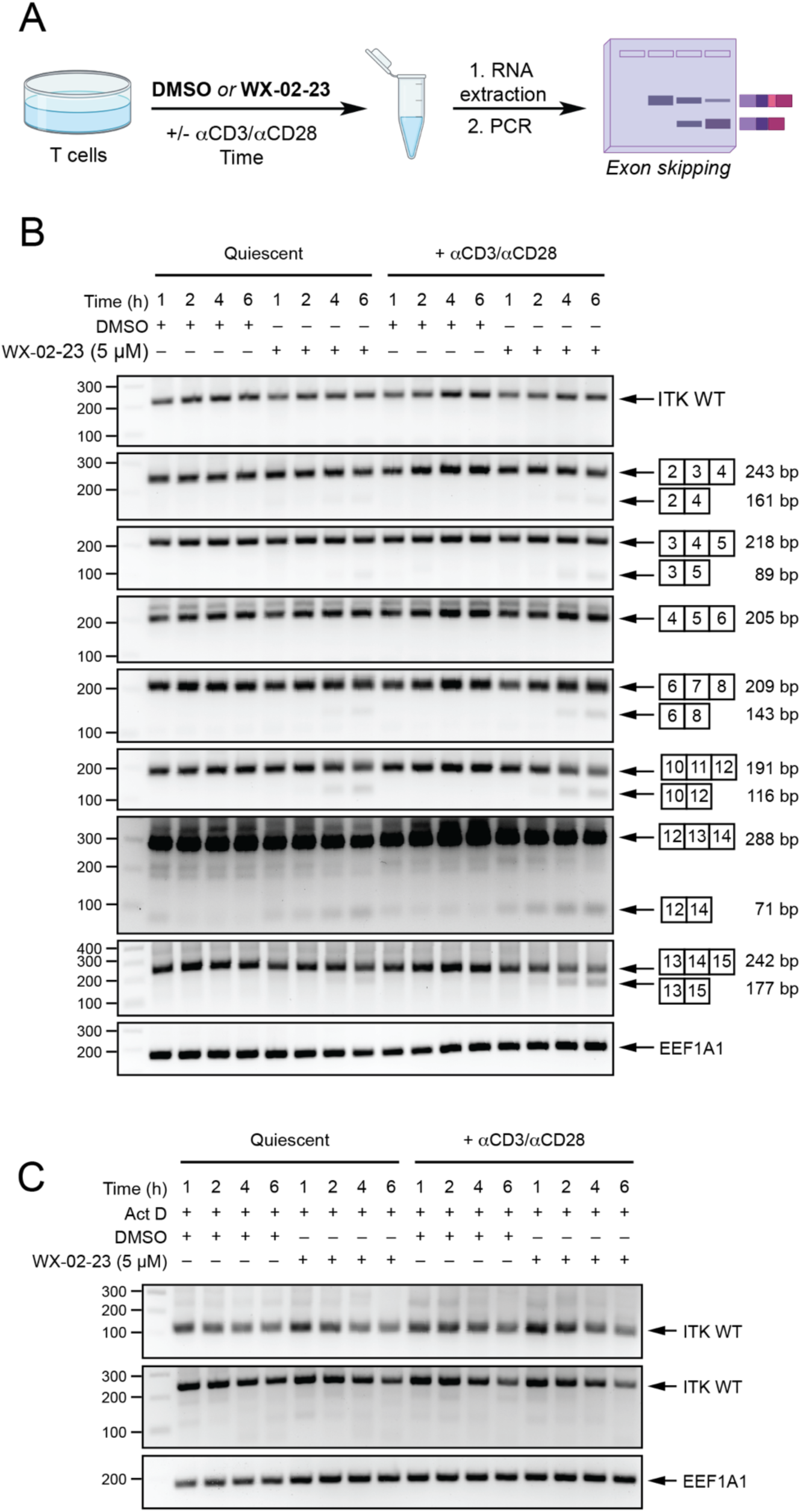
Contributions of alterntive splicing and WT ITK mRNA turnover to state-dependency of ITK protein level decrease. (A) Workflow for evaluation of ITK splicing changes by RT-PCR. Primary human T cells were treated with DMSO or WX-02-23 (5 µM) for 1-6 h with or without stimulation with αCD3/αCD28 antibodies, RNA was extracted and reverse transcribed, and PCR was used to generate amplified fragments using primers designed to span exons that were observed to be spliced out by RNA-sequencing. (B) Second replicate of RT-PCR analysis of alternative splicing of ITK mRNA induced by WX-02-23 in quiescent T cells versus stimulated T cells. Total RNA was isolated from quiescent T cells and stimulated T cells treated with DMSO or WX-02-23 (5 µM) for the indicated times (1, 2, 4 or 6 h) and reverse transcribed to cDNA. PCR primers were designed to detect exon skipping events of exon 3, 4, 5, 7, 11, 13 and 14 identified from RNA-seq data. RT-PCR was performed with primers that bind to exons on both sides of the target exon. PCR products were separated by agarose gel (2%) electrophoresis and visualized by ethidium bromide staining. Eukaryotic translation elongation factor 1 alpha 1 (EEF1A1) served as a loading control. (C) Second replicate of RT-PCR analysis evaluating effect of WX-02-23 on ITK mRNA stability in quiescent T cells versus stimulated T cells. The decay rates of ITK mRNA were assessed using RT-PCR in quiescent T cells and stimulated T cells treated with DMSO or EV96A (5 µM) following transcription inhibition by actinomycin D (Act D, 2.5 µg/mL) for the indicated times (1, 2, 4 and 6 h). Total RNA was isolated, RT-PCR analysis was performed, and PCR products were visualized by ethidium bromide staining. Eukaryotic translation elongation factor 1 alpha 1 (EEF1A1) served as a loading control.

**Supplementary figure 5.**
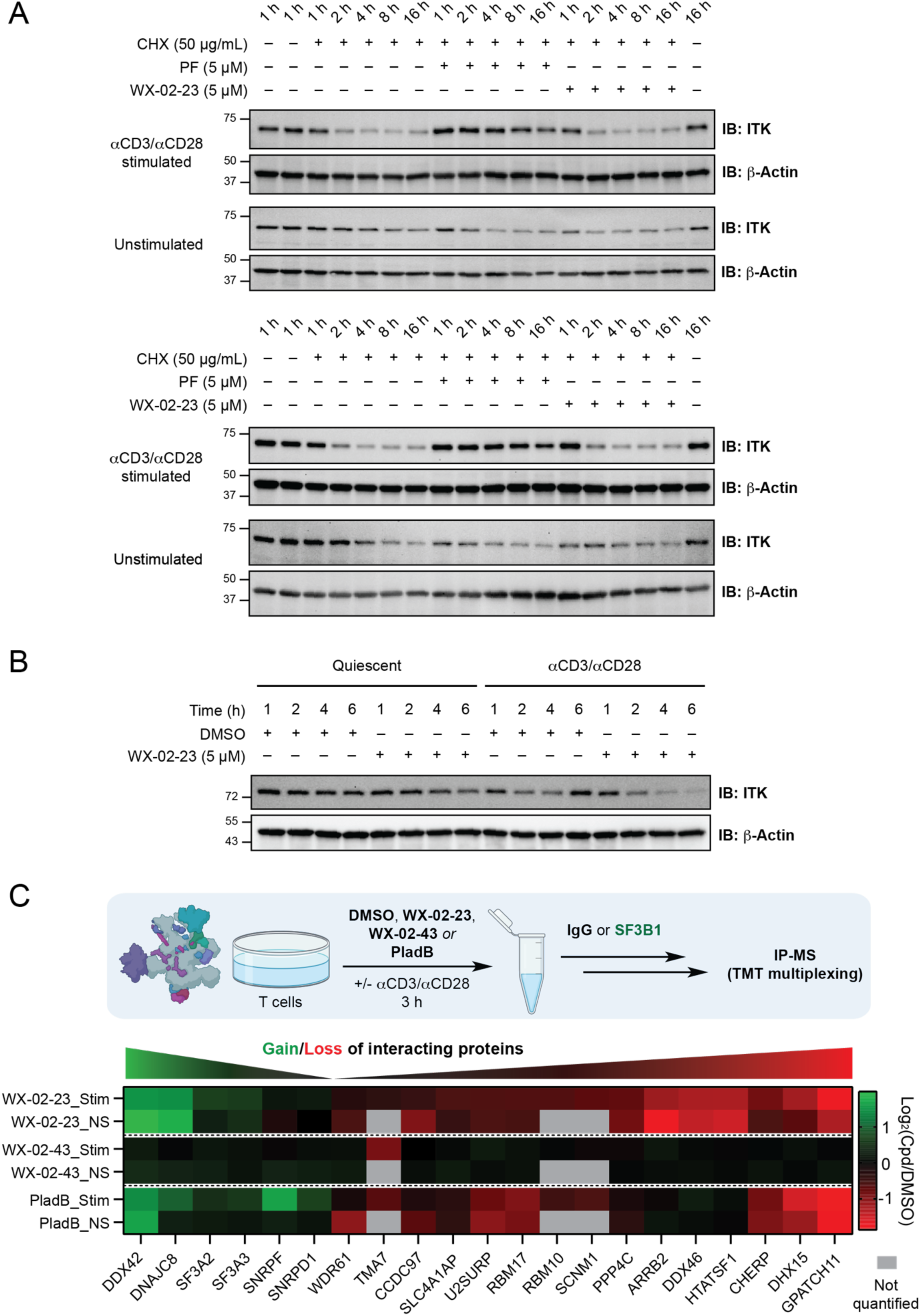
Contributions of ITK protein turnover and spliceosome composition to state-dependency of ITK protein level decrease. (A) Second and third experimental replicate showing ITK protein expression in quiescent T cells versus αCD3/αCD28 stimulated T cells, and the effect of PF-06465469 on ITK protein stability. The decay rates of ITK protein were assessed using Western blot in quiescent and stimulated T cells treated with DMSO, PF-06465469 (5 µM) or WX-02-23 (5 µM) following translation inhibition by cycloheximide (50 µg/mL) for the indicated times (1, 2, 4, 8, and 16 h). (B) Second experimental replicate showing the effect of T cell stimulation and WX-02-23 (5 µM) treatment on ITK protein expression. ITK protein expression was assessed using Western blot in quiescent T cells and stimulated T cells treated with DMSO or WX-02-23 (5 µM) for the indicated times (1, 2, 4, and 6 h). (C) Co-immunoprecipitation of proteins with SF3B1 antibody (CST #14434) following treatment of quiescent or stimulated T cells with DMSO, WX-02-23 (5 µM), WX-02-43 (5 µM), or Plad B (5 µM) for 3 h. Heatmap showing average log_2_(Cpd/DMSO) values of quantified proteins from 4-6 independent experiments (2-3 biological replicates) as representative values for compound-induced changes in SF3B1 interactomes in quiescent (NS) and stimulated (stim) T cells.

**Supplementary figure 6.**
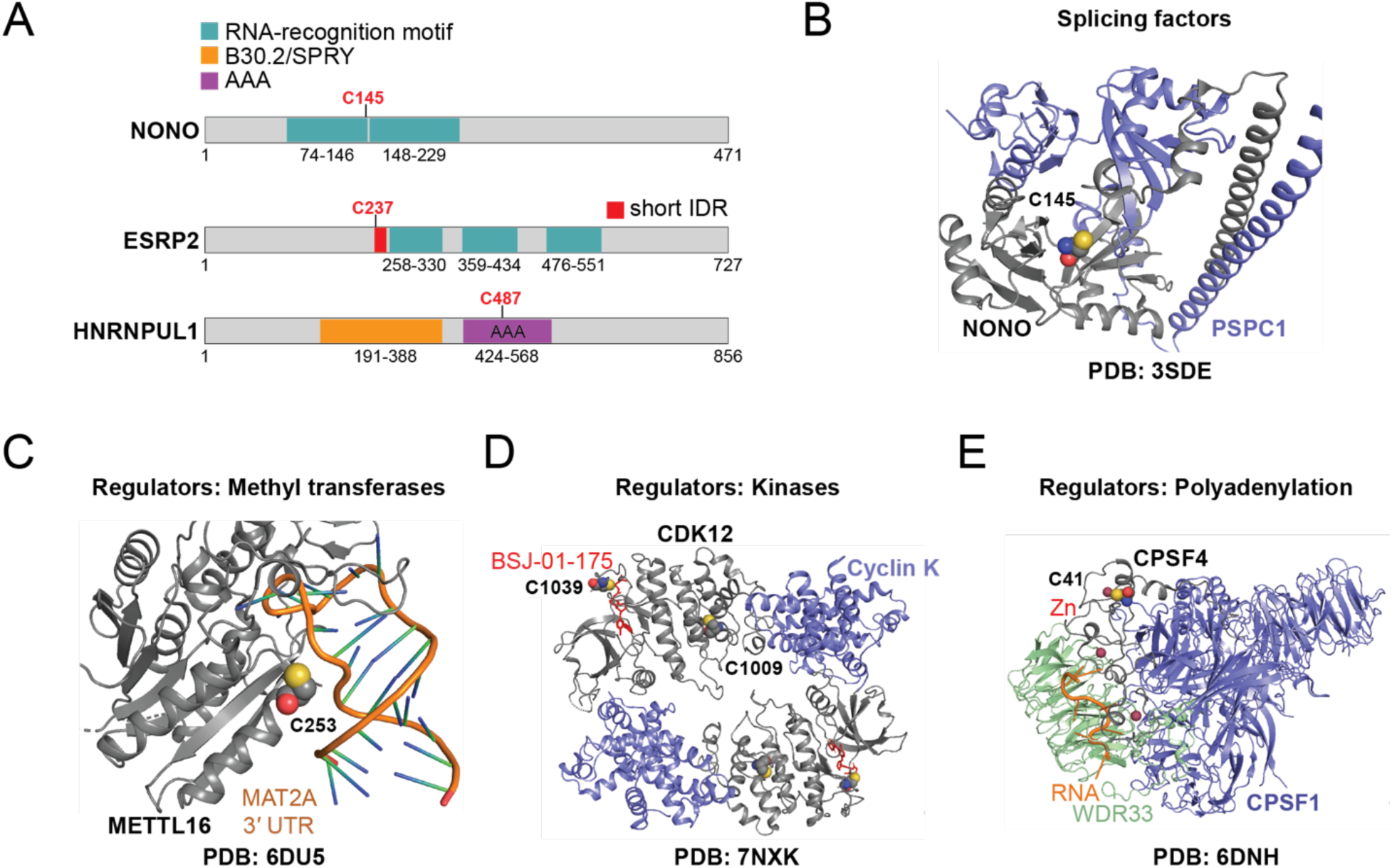
Ligandable cysteine residues from the splicing master list. (A) Protein sequences of targets from the splicing master list identified using cysteine-directed TMT-ABPP platforms. Cysteine residues are considered “liganded”, if scout fragments KB02 or KB05 show >75% competition for reactivity of that site. Specifically, liganded sites were identified in RNA-recognition motifs (NONO_C145), AAA-domains (HNRNPUL1_C487), and as part of short intrinsically disordered domains (ESRP2_C237). (B-E) Cryo-EM and crystal structures of representative splicing factor and splicing regulator targets of scout fragments identified using cysteine-directed TMT-ABPP platforms. Proteins of interest are shown in gray and the cysteine residues are shown as spheres colored by elements. (B) Crystal structure of a paraspeckle-protein heterodimer, PSPC1/NONO (PDB: 3SDE) with ligandable cysteine NONO_C145 in RNA-binding protein NONO located at the protein-protein interaction surface with PSPC1; (C) Crystal structure of METTL16 catalytic domain in complex with MAT2A 3’UTR hairpin 6 (PDB: 6DU5) with ligandable cysteine METTL16_C253 located at the protein-RNA interface; (D) Crystal structure of cyclin-dependent kinase CDK12/Cyclin K in complex with the allosteric covalent inhibitor BSJ-01-175 bound to CDK12_C1039 (PDB: 7NXK), with ligandable cysteine CDK12_C1030 located at the protein-protein interaction surface with Cyclin K; (E) Cryo-EM structure of human CPSF-160-WDR33-CPSF-30-PAS RNA complex (PDB: 6DNH) with ligandable cysteine in cleavage and polyadenylation specificity factor subunit CPSF4_C41 located as part of a CPSF4 Zn-finger.

**Supplementary figure 7.**
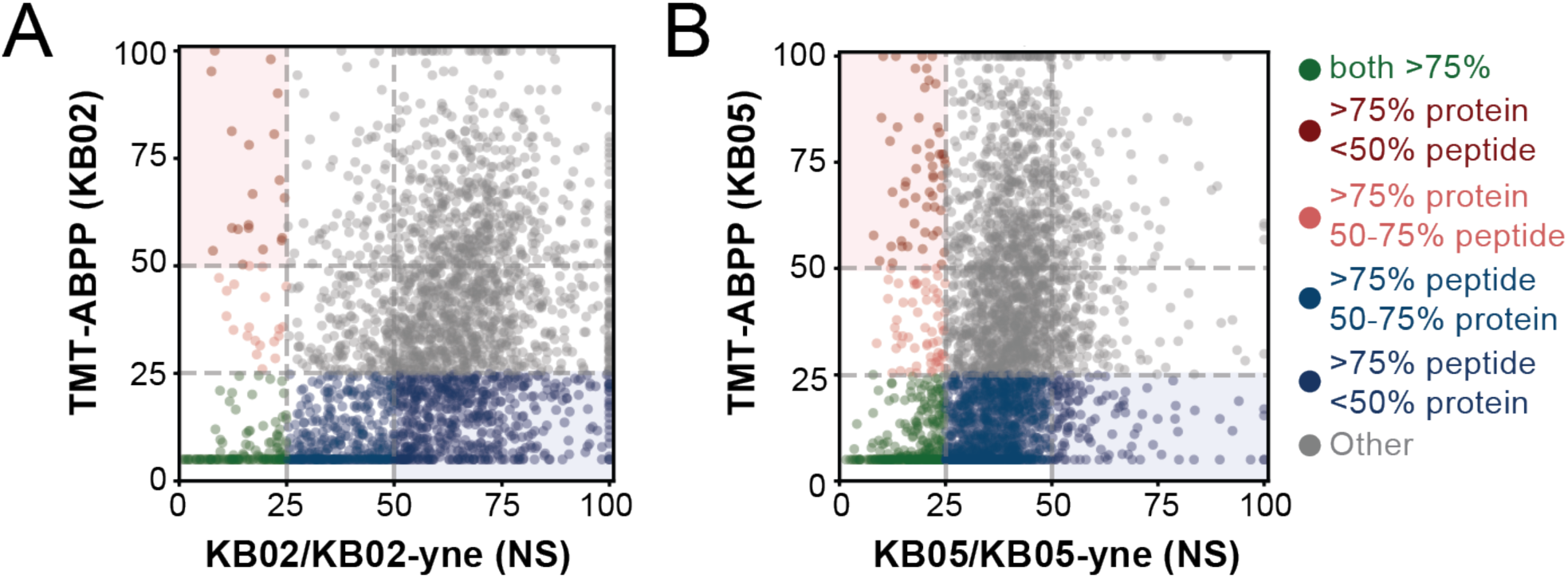
Comparison between cysteine-directed TMT-ABPP and protein-directed TMT-ABPP competition with scout fragments. (A-B) Scatter plot representation showing comparison between cysteine-directed TMT-ABPP ligandability values[1] and protein-directed TMT-ABPP competition values with KB02-yne (A) and KB05-yne (B) scout fragment-based chemical probes in quiescent T cells. Proteins uniformly showing >75% ligandability and competition are nominated as highest priority targets for follow up studies and ligand development. Proteins uniquely ligandaded in cysteine-directed TMT-ABPP data are shown in blue. Proteins uniquely competed in protein-directed TMT-ABPP competition data are shown in red.

### (B) Supplementary Data Set Legends

**Supplementary Data Set 1**

**Tab 1.** Table of contents for **Supplementary Dataset 1**.

**Tab 2 (Table S1A).** *In vitro* site-of-labeling experiment with GluC and Trypsin tandem digest showing labeling of ITK FL protein with EV96 WT ITK.

**Tab 3 (Table S1B).** *In vitro* site-of-labeling experiment with GluC and Trypsin tandem digest showing labeling of ITK FL protein with EV97 WT ITK.

**Tab 4 (Table S2).** *In situ* site-of-labeling experiment with multi protease digest showing percent signal intensity values of EV96, MG132 or EV96 and MG132-treated samples with or without T cell stimualtion normalized to DMSO_NS channels.

**Tab 5 (Table S3).** *In situ* TMT-ABPP competition experiment with EV96, EV97, and PF against EV96A-yne probe in expanded T cells.

**Tab 6 (Table S4).** *In situ* TMT-ABPP competition experiment with EV96, EV97, and PF against EV96A-yne probe in Jurkat cells.

**Tab 7 (Table S5).** TMT-exp proteomic data showing changes in protein expression following treatment with DMSO, EV96, EV97, or PladB with or without T cell stimulation.

**Tabs 8 (Tables S6).** TMT-exp proteomic data showing the effect of WX-02-23, WX-02-43, and PladB treatment on co-immunoprecipitation of proteins with SF3B1.

**Tab 9 (Table S7).** The splicing master list reference table was consolidated by combining proteins reported to be components of the spliceosome, splicing factors, or splicing regulators (kinases, phosphatases, methyl transferases, and proteins involved in polyadenylation.

**Tab 10 (Table S8).** *In vitro* TMT-ABPP competition experiment with scout fragments KB02 and KB05 against KB02-yne and KB05-yne chemical probes in expanded and activated T cells. Scout fragment ligandability data from Cell 2020 (PMID: 32730809) is also included, as well as the coloring annotation for figures 7E and S7.

**Tab 11 (Table S9).** TMT-ABPP broad ligandability data from Cell 2020 (PMID: 32730809) searched against SF1-isoform 6 FASTA database.

**Supplementary Dataset 2S**

**Tab 1.** Table of contents for **Supplementary Dataset 2S**.

**Tab 2 (Table 2S-1).** RNA-sequencing data for primary human T cells treated with DMSO, WX-02-23 (5 µM), WX-02-43 (5 µM), or PladB (5 µM) with or without T cell stimulation with αCD3/αCD28 antibodies.

**Tabs 3-9 (Table 2S-2A – 2S-2G).** Alternative splicing data for primary human T cells treated with DMSO, WX-02-23 (5 µM), WX-02-43 (5 µM), or PladB (5 µM) with or without T cell stimulation with αCD3/αCD28 antibodies.

**Supplementary Dataset 3**

Synthetic procedures and characterization.

### (C) Biological Methods

#### EXPERIMENTAL MODEL AND SUBJECT DETAILS

##### Cell lines

Jurkat cells (ATCC) were grown in RPMI media supplemented with 10% heat-inactivated FBS, *L*-glutamine (2 mM), penicillin (100 U/mL), and streptomycin (100 mg/mL) in 5% CO_2_ tissue culture incubators.

##### Mouse T Cell Isolation

Studies using primary mouse T cells were carried out following protocols approved by The Scripps Research Institute. Mouse T cells were isolated from mouse splenocytes using EasySep Mouse T Cell Isolation Kit (STEMCELL Technologies, negative selection) following manufacturer’s instructions. Mouse T cells were grown in RPMI media supplemented with 10% heat-inactivated FBS, *L*-glutamine (2 mM), penicillin (100 U/mL), streptomycin (100 mg/mL), and β-Mercaptoethanol (55 µM) in 5% CO_2_ tissue culture incubators.

##### T Cell Isolation

Studies using primary human T cells were carried out with samples from healthy donors following protocols approved by The Scripps Research Institute and The Rockefeller University Internal Review Board. Blood from de-identified healthy donors was obtained from Scripps Research Normal Blood Donor Services, NYBC, STEMCELL Technologies or Rockefeller Hospital. Peripheral blood mononuclear cells (PBMCs) were isolated using a Lymphoprep gradient (STEMCELL Technologies) following a slightly modified protocol compared to manufacturer’s instructions. Briefly, freshly isolated blood (25 mL) was layered on top of cold Lymphoprep (12.5 mL) in a 50 mL Falcon tube without disturbing the Lymphoprep layer. Tubes were spun down (930 *g*, 20 min, 23 °C) in a benchtop centrifuge with brake off. The plasma and Lymphoprep layers, containing PBMCs, were transferred to new Falcon tubes with 2-fold dilution with PBS and pelleted by centrifugation (520 *g*, 8 min, 4 °C). Cells were re-combined into one Falcon tube using PBS and pelleted (520 *g*, 8 min, 4 °C). Pan T cells were isolated from PBMCs using EasySep Human T Cell Isolation Kit (STEMCELL Technologies, negative selection) following manufacturer’s instructions.

#### STATISTICAL ANALYSIS

Statistical analysis was performed using GraphPad Prism (v9.3.1). Statistical significance is determined using either Dunnett’s multiple comparisons test or Tukey’s multiple comparisons test compared to respective control groups. The following statistical significance values are indicated in the figures and reported in the corresponding figure legends (*, p < 0.05; **, p < 0.01).

#### METHODS DETAILS

##### T Cell Activation

Non-tissue culture treated 6-well plates or 15 cm plates were pre-coated with αCD3 (5 µg/mL, BioXCell) and αCD28 (2 µg/mL, BioXCell) antibodies in PBS (2 mL/well for 6-well plates, 15 mL/plate for 15 cm plates) and incubated at 37 °C for 2 h or stored at 4 °C overnight in the dark, followed by incubation at 37 °C for 30 min. Plates were washed with PBS (5 mL/well or 10 mL/15 cm plate, 2 times) right before use. Freshly isolated T cells were resuspended in RPMI media (1×10^6^ cells/mL) supplemented with 10% heat-inactivated FBS, *L-*glutamine (2 mM), penicillin (100 U/mL), and streptomycin (100 mg/mL) [referred to as “T cell media”], plated onto pre-coated plates and incubated at 37 °C in a 5% CO_2_ incubator with or without compound treatment for the reported time.

##### T Cell Expansion

Non-tissue culture treated 6-well plates were pre-coated with αCD3 (1.5 µg/mL, BioXCell) in PBS (3 mL/well) and incubated at 37 °C for 2 h or stored at 4 °C overnight in the dark, and then incubated at 37 °C for 10-30 min before using. Plates were washed with PBS (5 mL/well, 2 times) and freshly isolated T cells (1×10^6^ cells/mL, T cell media supplemented with 1 µg/mL αCD28 antibody) were plated onto pre-coated plates and kept at 37 °C in a 5% CO_2_ incubator for 3 days. After this time, cells were combined into Falcon tubes, pelleted (520 *g*, 5 min, 4 °C), and washed with PBS (1 x 40 mL). Cells were then resuspended in fresh T cell media and kept at 37 °C in a 5% CO_2_ incubator for 12-14 days, splitting the cells with fresh media every 3-4 days. Cells were then pelleted (520 *g*, 5 min, 4 °C), washed with PBS and either resuspended in fresh T cell media for *in situ* treatments, or flash frozen and stored at −80 °C for *in vitro* treatments.

##### Compound screen for inhibitors of T cell activation (related to figures 1E, 1F, S1B-S1F)

Non-tissue culture treated flat bottom 96-well plates were pre-coated with αCD3 (5 µg/mL, BioXCell) and αCD28 (2 µg/mL, BioXCell) antibodies in PBS (100 µL/well) and kept at 4 °C overnight. The following day, plates were warmed to 37 °C for 1 h and washed with PBS (200 µL/well, 2 times) before use. Freshly isolated T cells were resuspended at 2×10^6^ cells/mL in T cell media and seeded in the pre-coated plates (100 µL/well). Compound stocks in DMSO (200x) were diluted to 2x in T cell media in a separate V-bottom plate and added to the pre-coated plate with T cells (100 µL/well, 1×10^6^ cells/mL final cell density). The outer wells of each plate were filled with PBS (150 µL/well) to avoid edge effect. The plates were then kept at 37 °C for 24 h. After 24 h, cells were transferred to a 96-well round bottom plate, spun down (520 *g*, 5 min, 4 °C) and further processed for flow cytometry analysis.

##### Flow cytometry analysis (related to figures 1F, 2E, S1D-S1F)

Flow cytometry analysis of T cell activation related to figure 1E was performed following previously described protocol[1]. Flow cytometry analysis of T cell activation related to figures 2E, S1D-S1F was performed following a modified protocol. Briefly, cells were washed once with PBS and resuspended in fixable near-IR LIVE/DEAD cell stain (Invitrogen, 1:4000 in PBS). Following incubation at ambient temperature for 20 min without light, cells were pelleted and washed with PBS. Cells were then resuspended in staining buffer (1% FBS in PBS, 25 µL/well) with Fc blockers (BioLegend, 1:200) and incubated on ice for 15 min. Following this incubation, 25 µL of antibody master mix (PB-CD4, PerCP-Cy5.5-CD8, and PE-CD25 antibodies, BioLegend, 1:200 in staining buffer) was added to each sample, and cells were stained on ice for additional 25 min. After the staining step, cells were pelleted and washed with PBS. Cells were then fixed in 4% PFA (Wako, 100 µL/well) on ice for 15 min, pelleted, washed with PBS, and resuspended in FACS buffer (1% FBS, 2 mM EDTA in PBS) for flow cytometry analysis. Flow cytometry analysis was performed on an Attune NxT Flow Cytometer (Thermo Fisher) with a CytKick Autosampler and the resulting data was analyzed using FlowJo software. Compensation was performed for each experiment with UltraComp eBeads (Invitrogen) prepared according to manufacturer’s instructions.

##### Western blot analysis (related to figures 2, 5, S2, and S5)

Expanded primary human T cells (8 x 10^6^ cells/treatment) were resuspended in fresh T cell media at 2 x 10^6^ cells/mL and treated with DMSO or test compounds for the indicated times at 37 °C in a 5% CO_2_ incubator. Following this incubation period, cells were pelleted (600 *g*, 5 min, 4 °C), washed with PBS, transferred to 1.5 mL Eppendorf tubes, flash-frozen, and stored at −80 °C until further analysis. On the day of the analysis, cell pellets were thawed on ice, resuspended in cold PBS containing protease inhibitor cocktail (Roche) and lysed by sonication (3 x 8 pulses, 60% duty cycle). Protein concentrations were adjusted to 1-2 mg/mL, 4x loading buffer was added, and the samples were heated at 95 °C for 7 min. Proteins were resolved using SDS-PAGE (4–20% Criterion™ TGX Stain-Free™ Protein Gel, BioRad) and transferred to 0.45 μm polyvinylidene fluoride membranes (Cytiva). The membranes were blocked with 5% milk or 5% BSA in Tris-buffered saline (20 mM Tris-HCl pH 7.6, 150 mM NaCl) with 0.1% tween 20 (TBS-T) buffer at room temperature for 30 min, washed 3 times with TBS-T, and incubated with primary antibodies in 5% BSA in TBS-T at 4 °C overnight. The membranes were washed 3 times with TBS-T, incubated with HRP-conjugated secondary antibodies in 5% BSA in TBS-T at room temperature for 1 h and washed 5 times with TBS-T. The immunoreactive bands were detected using Pierce^TM^ ECL Western Blotting Substrate (Thermo Fisher Scientific) on a ChemiDoc^TM^ MP Imaging System (Bio-Rad). Blots were quantified with densiometric analysis using ImageJ software (NIH).

##### RNA extraction and RT-PCR analysis (related to figures 4 and S4)

Expanded primary human T cells (8 x 10^6^ cells/treatment) were resuspended in T cell media at 2 x 10^6^ cells/mL and treated with DMSO or test compounds at 37 °C in a 5% CO_2_ incubator for the indicated times. Following this incubation period, the cells were pelleted (600 *g*, 5 min, 4 °C), washed with PBS, transferred to 1.5 mL Eppendorf tubes, flash-frozen, and stored at −80 °C until further analysis. Total RNA was extracted from cells using Direct-zol RNA MicroPrep Kit (Zymo research) according to manufacturer’s instructions, and RNA concentration was determined using NanoDrop. RNA (0.5 μg/treatment) was reverse-transcribed into complementary DNA (cDNA) using iScript cDNA Synthesis Kit (Bio-Rad) according to manufacturer’s instructions. RT-PCR reactions were set up with 10 ng cDNA per reaction and 0.5 µM forward and reverse primers along with Q5 High-Fidelity 2x Master Mix (New England Biolabs). Reactions were carried out on C1000 Touch^TM^ Thermal Cycler (Bio-Rad). The following PCR settings were used for the RT-PCR: 5 min at 94 °C; 35 cycles of 30 sec at 94 °C, 30 sec at 60 °C, and 2 min at 68 °C; 5 min at 72 °C. PCR products were separated on a 2% agarose gel (Lonza) and stained with ethidium bromide (Genesee Scientific). Relative gene expression was quantified with densiometric analysis using ImageJ software (NIH) and normalized using eukaryotic translation elongation factor alpha 1 (EEF1A1). Primers used for RT-PCR are listed in Key resources table.

##### In Gel Fluorescence (related to figure 2D)

Cell lysates prepared for mass spectrometry analysis of competition experiments with elaborated compounds were used for gel-based visualization of labeling and competition. Rhodamine was conjugated for visualization of modified proteins using copper-catalyzed azide-alkyne cycloaddition reaction (CuAAC). Reagents for the reaction were pre-mixed prior to their addition. Briefly, rhodamine-azide (1 µL, 1.25 mM stock in DMSO, 25 µM final concentration), copper sulfate (1 µL, 50 mM stock in water, 1 mM final concentration), tris(benzyltriazolylmethyl)amine ligand (TBTA; 3 µL, 1.7 mM stock in DMSO/*t*BuOH, 1:4, 100 µM final concentration), and tris(2-carboxyethyl)phosphine hydrochloride (TCEP; 1 µL, fresh 50 mM stock in water, 1 mM final concentration) were combined in an Eppendorf tube, vortexed and spun down. The reaction mixture (3 µL) was then added to each sample (25 µL) and the samples were incubated at ambient temperature for 1 h. Following the incubation period, samples were quenched with 9.3 µL 4x sample buffer and boiled at 95 °C for 5 min. Samples were then resolved on SDS-PAGE (4–20% Criterion™ TGX Stain-Free™ Protein Gel, BioRad) and analyzed using the appropriate fluorescence channel on a ChemiDoc^TM^ MP Imaging System (Bio-Rad).

##### Proteomic platforms: *in vitro* site-of-labeling (related to figure 1C, supplementary tables S1A and S1B)

###### Sample preparation

In a low binding 1.5 mL Eppendorf tube, full length ITK (Carnabio, 50 µL, 20 nM) was treated with EV96 (1 µL, 50 µM, 1 µM final concentration) or EV97 (1 µL, 50 µM, 1 µM final concentration) at ambient temperature for 1 h. Cold acetone was added (200 µL) and the samples were kept at −20 °C for 30 min to promote protein precipitation. The protein was pelleted (16,000 *g*, 10 min, 4 °C) and acetone was carefully aspirated without touching the pellet. The resulting precipitate was resuspended in 50 µL of 8M urea TEAB buffer containing DTT (2.2 g urea, 200 µL 1M TEAB pH 8.5, 2 mL H_2_O, 6.2 mg DTT) with sonication. The samples were heated at 65 °C for 15 min and allowed to cool down before the alkylation step. Iodoacetamide (2.5 µL, 400 mM, 20 mM final concentration) was then added and the samples were incubated for an additional 30 min at 37 °C. The sample was diluted with 400 µL TEAB buffer (50 mM, pH 8.5) to decrease urea concentration (0.9M) and incubated with GluC (1 µL, 0.5 µg/µL in H_2_O) for 2 h at 37 °C. Trypsin (4 µL, 0.25 µg/µL in trypsin buffer) and CaCl_2_ (5 µL, 100 mM, 1 mM final concentration) were then added and the digestion was allowed to proceed overnight at 37 °C. The following day the samples were acidified by the addition of formic acid (5 µL, 0.05% final concentration) and the solvent was removed using SpeedVac vacuum concentrator. The samples were then desalted using StageTips (100 µL). Briefly, the tips were activated and equilibrated (2 x 100 µL CH_3_CN, 3 x 100 µL 0.1% FA : 5% CH_3_CN : 95% H_2_O), the sample was re-dissolved in 100 µL buffer A (0.1% FA : 5% CH_3_CN : 95% H_2_O) and loaded on the tip, washed with buffer A (200 µL, 3 times) and eluted with buffer B (200 µL; 0.1% FA : 20% CH_3_CN : 80% H_2_O) into a new Eppendorf tube. The solvent was removed using a SpeedVac vacuum concentrator overnight.

###### Liquid chromatography tandem mass spectrometry analysis (LC-MS/MS)

Samples were re-suspended in buffer A (10 µL) and analyzed by liquid chromatography tandem mass spectrometry (7.5 µL injection) using Orbitrap Eclipse mass-spectrometer (Thermo Fisher) with UltiMate 3000 Series Rapid Separation LC system. Briefly, the peptides were eluted onto a capillary column (2 µm, 100 Å, 75µm x 25 cm) at 0.25 µL/min flow rate and separated using the following gradient: 5% LC-MS buffer B (CH_3_CN, 0.1% FA) in LC-MS buffer A (H_2_O, 0.1% FA) from 0-10 min, 5%–20% buffer B from 20-120 min, 20%–45% buffer B from 120-140 min, 45%–95% buffer B from 140-145 min, 95% buffer B from 145-147 min, 5% buffer B from 147-149 min, 95% buffer B from 149-151 min, 5% buffer B from 151-160 min) [“standard 160 min gradient”].

Downstream analysis included the following scan sequence: (1) MS1 master scan (Orbitrap analysis, resolution 240,000, 400−1600 m/z, RF lens 40%, normalized AGC Target 250%, automatic maximum injection time, profile mode) with enabled dynamic exclusion (repeat count 1, duration 60 s); (2) Selection of top ten ions for MS2 analysis; (3) MS2 analysis, which consisted of quadrupole isolation (isolation window 0.7) of precursor ion followed by collision-induced dissociation (CID) in the ion trap (fixed collision energy mode, normalized collision energy 30%, maximum CID activation time 10 ms, Activation Q 0.25). The MS2 files were extracted from the raw files using RAW Converter (version 1.1.0.22; available at http://fields.scripps.edu/rawconv/), uploaded to Integrated Proteomics Pipeline (IP2), and searched using the ProLuCID algorithm (publicly available at http://fields.scripps.edu/downloads.php) using a reverse concatenated, non-redundant variant of the Human UniProt database (release-2016_07). Cysteine residues were searched with a static modification for carboxyamidomethylation (+57.02146 Da) and a dynamic modification for EV96/EV97 (+402.1580 Da). Peptides were required to be at least 5 amino acids long, and to have at least one tryptic terminus. ProLuCID data was filtered through DTASelect (version 2.0) to achieve a peptide false-positive rate below 1%.

###### Data processing and analysis

Peptide lists were downloaded from Integrated Proteomics Pipeline and the following filters were applied: removal of non-unique peptides. The final data is reported in Supplementary Tables S1A and S1B.

##### Proteomic platforms: *in situ* cysteine-directed site-of-labeling TMT-ABPP (related to figure 1D, supplementary table S2)

###### Sample preparation

Two 10-plex experiments with primary human T cells from different biological donors were processed (5 experimental conditions in duplicate for each donor). Expanded T cells were pelleted (600 *g*, 5 min, 4 °C) and re-suspended in fresh T cell media at 1 x 10^6^ cells/mL for compound treatment. T cells were treated with EV96 (10 µM), EV96 (10 µM) and MG132 (10 µM), MG132 (10 µM) or DMSO control in 15 cm plates (25 mL/plate, 50 x 10^6^ cells/treatment) with or without αCD3/αCD28 stimulation for 3 h. For treatment conditions that required T cell activation, plates were pre-coated with 15 mL PBS containing αCD3 (5 µg/mL, BioXCell) and αCD28 (2 µg/mL, BioXCell) overnight at 4 °C or for 2 h at 37 °C and washed with PBS (7 mL, 3 times) prior to treatment. Following treatment, cells were transferred into Falcon tubes (50 mL) and pelleted (600 *g*, 5 min, 4 °C). Cells were then transferred into new low binding 1.5 mL Eppendorf tubes using 1 mL cold PBS, pelleted (600 *g*, 5 min, 4 °C), flash frozen in liquid nitrogen, and stored at −80 °C until the day of analysis.

On the first day of the mass spectrometry protocol, cell pellets were thawed on ice for 10 min, and cells were lysed by sonication (3 x 8 pulses, 60% duty cycle) in ice cold PBS (560 µL/treatment). Protein concentrations were measured using a standard DC assay (Bio-Rad) and normalized to 1-2 mg/mL. Normalized cell lysates were transferred to new low binding Eppendorf tubes (500 µL/tube), iodoacetamide desthiobiotin (Santa Cruz Biotechnologies) was added to each tube and the mixture was incubated at ambient temperature for 1 h. Following the incubation period, samples were transferred on ice and the proteins were precipitated using MeOH/CHCl_3_ precipitation. First, cold MeOH (500 µL) and CHCl_3_ (150 µL) were added to each Eppendorf tube and the proteins were pelleted by centrifugation (10,000 *g*, 10 min). The solvent was aspirated without disturbing the protein disk. At this point, the samples can be stored overnight at −80 °C before proceeding with the rest of the protocol. Additional MeOH (400 µL) was added, and the samples were sonicated to break down protein disks. Proteins were pelleted once again (10,000 *g*, 10 min), and re-solubilized in 8M urea TEAB buffer containing DTT (90 µL; 2.2 g urea, 6.2 mg DTT, 2 mL H_2_O, 200 µL 1M TEAB buffer, pH 8.5) with sonication. The samples were then heated at 65 °C for 20 min on a heating block and cooled down (5 min) before the alkylation step. Iodoacetamide (10 µL, 500 mM freshly made solution in H_2_O, 50 mM final concentration) was then added and the samples were incubated at 37 °C for an additional 30 min.

###### Multi protease digestion and streptavidin enrichment

The samples were diluted with 600 µL TEAB buffer (50 mM, pH 8.5) to reduce the urea concentration. GluC (2 µL, 0.5 µg/µL in H_2_O), trypsin/LysC (6 µL, 0.25 µg/µL in trypsin/LysC buffer), and CaCl_2_ (8 µL, 100 mM, 1 mM final concentration) were then added and the digestion was allowed to proceed overnight at 37 °C with shaking. The following day, pre-washed streptavidin-agarose beads were added to each sample (25 µL compact beads, resuspended in 50 mM TEAB buffer, pH 8.5, containing 150 mM NaCl and 0.2% NP40 (enrichment buffer); 300 µL/sample) and the mixture was rotated for 3 h at ambient temperature. Briefly, for a 10-plex experiment, streptavidin-agarose beads (550 µL, 50% slurry) were washed with wash buffer (50 mM TEAB buffer, pH 8.5, containing 150 mM NaCl and 0.1% NP40; 2 times x 10 mL) and re-suspended in 3500 mL of enrichment buffer. Following the enrichment step, the beads were pelleted (2000 *g*, 1 min), transferred to BioSpin columns and washed extensively (3 x 1 mL of wash buffer, 3 x 1 mL PBS, 3 x 1 mL water). The peptides were eluted into a new Eppendorf tube by the addition of 50% CH_3_CN : H_2_O buffer, containing 0.1% FA. The solvent was removed by SpeedVac vacuum concentrator.

###### TMT labeling

The resulting protein digest was re-suspended in 30% CH_3_CN in water, containing 0.1% FA (100 µL/sample), with sonication and used for TMT labeling. Briefly, TMT tags (3 µL/sample, 20 µg/µL in dry CH_3_CN) were added to the corresponding samples (“channels”), the samples were vortexed, spun down and incubated for 1 h at ambient temperature. The tags were quenched by the addition of NH_2_OH (3 µL, 5% in H_2_O), vortexed, and incubated for 15 min. The samples were acidified by the addition of formic acid (5 µL/sample) and combined in a new Eppendorf tube. The solvent was removed overnight using SpeedVac vacuum concentrator and the final sample was desalted on a SepPak (C18, 50 mg) column. Briefly, the column was activated (2 x 1 mL CH_3_CN) and equilibrated (3 x 1 mL Buffer A; 0.1% FA : 5% CH_3_CN : 95% H_2_O), the sample was re-dissolved in 1 mL buffer A and loaded on the column, washed with buffer A and eluted with buffer B (0.1% FA : 20% CH_3_CN : 80% H_2_O) into a new Eppendorf tube. The solvent was removed overnight using SpeedVac vacuum concentrator and the sample was high pH fractionated using HPLC fractionator.

###### High pH HPLC fractionation (general protocol for all LC-MS/MS/MS workflows)

Desalted sample was re-suspended in Buffer A (500 µL) and fractionated into a 96 deep-well plate using HPLC (Agilent). The peptides were eluted onto a Zorbax column (Extend-C18, 3.5 µm, 4.6 x 250 mm) at 0.5 mL/min flow rate and separated using the following gradient: 100% buffer C from 0-2 min, 0%–13% buffer D from 2-3 min, 13%–50% buffer D from 3-60 min, 50%–80% buffer D from 60-70 min, 100% buffer D from 71-75 min, 100%–0% buffer D from 75-76 min, 100% buffer C from 76-85 min, 0%–13% buffer D from 85-88 min, 13%–80% buffer D from 88-90 min, 80% buffer D from 90-95 min, 100% buffer B from 96-101 min, 0%–13% buffer D from 101-104 min, 13%–80% buffer D from 104-106 min, 80% buffer D from 106-111 min, and 80%–0% buffer D from 111-112 min (buffer C: 10 mM aqueous NH_4_HCO_3_; buffer D: CH_3_CN). The receiving plate contained 10 µL 20% formic acid in each well to acidify eluting peptides and the collection was set to a time-dependent mode in the following order: A1-A12, B1-B12, C1-C12, D1-D12, E1-E12, F1-F12, G1-G12, H1-H12. The solvent was removed overnight using SpeedVac vacuum concentrator and the sample was re-suspended in Buffer B and concatenated by combining wells from each column (e.g., A1-H1) into a total of 12 fractions. The solvent was removed overnight using SpeedVac vacuum concentrator and each fraction was re-suspended in Buffer A (10 µL) for LC-MS/MS/MS analysis.

###### TMT liquid chromatography mass-spectrometry (TMT LC-MS/MS/MS, general protocol for all LC-MS/MS/MS workflows)

Samples were analyzed by liquid chromatography tandem mass spectrometry using standard 160 min gradient. Downstream analysis included the following scan sequence: (1) MS1 master scan (Orbitrap analysis, resolution 120,000, 400−1600 m/z, RF lens 40%, normalized AGC Target 250%, automatic maximum injection time, profile mode) with enabled dynamic exclusion (repeat count 1, duration 60 s); (2) Selection of top ten precursor ions for MS2/MS3 analysis; (3) MS2 analysis, which consisted of quadrupole isolation (isolation window 0.7) of precursor ion followed by collision-induced dissociation (CID) in the ion trap (Standard AGC, normalized collision energy 35%, maximum injection time 120 ms); (4) Real-time search (RTS) and synchronous precursor selection (SPS) enabled selection of up to 20 MS2 fragment ions for MS3 analysis; (5) Fragmentation of MS3 precursors by HCD and analysis using Orbitrap (collision energy 55%, normalized AGC Target 500%, maximum injection time 118 ms, 60,000 resolution). For MS3 analysis, MS isolation window was set at 0.7. The MS1, MS2 and MS3 files were extracted from the raw files using RAW Converter (version 1.1.0.22; available at http://fields.scripps.edu/rawconv/), uploaded to Integrated Proteomics Pipeline (IP2), and searched using the ProLuCID algorithm (publicly available at http://fields.scripps.edu/downloads.php) using a reverse concatenated, non-redundant variant of the Human UniProt database (release-2016_07). Cysteine residues were searched with a static modification for carboxyamidomethylation (+57.02146 Da) and a differential modification accounting for the reaction with iodoacetamide desthiobiotin probe (+398.25292 Da; site-of-labeling experiments). *N*-termini and lysine residues were searched with a static modification corresponding to the TMT tag (+229.1629 Da for 10-plex, +304.2071 Da for 16-plex TMT experiments). Peptides were required to be at least 5 amino acids long, and to have at least one tryptic terminus. The same parameters were used during the RTS feature at the time of data acquisition. ProLuCID data was filtered through DTASelect (version 2.0) to achieve a peptide false-positive rate below 1%. The MS3- based peptide quantification was performed with reporter ion mass tolerance set to 20 ppm with Integrated Proteomics Pipeline (IP2).

###### Data processing and analysis

Two 10-plex experiments with primary human T cells from different biological donors were analyzed (5 experimental conditions in duplicate for each donor). Each experiment was processed individually using normalized protein intensities (peptide total intensity of each channel is normalized by the sum of signal intensities for all peptides in the same channel, normalization performed by Integrated Proteomics Pipeline). The following filters were applied: removal of non-unique peptides, removal of half-tryptic peptides, removal of peptides with more than one internal missed cleavage site, removal of peptides with low (< 5,000) average of reporter ion intensities for control channels, and peptides with high variation between the replicate control channels (coefficient of variation > 0.5). Peptide ratios (signal intensity / sum of signal intensities per peptide) were calculated and ratios of all peptides per labelled residue were averaged. The ratios of treatment groups (EV96_ns, EV96_stim, EV96_stim_MG132, stim_MG132) to control (DMSO_ns) were calculated and values are provided as percent to control. Cysteine aggregation was performed to aggregate signal intensities for multiple cysteines on the same peptide and multiple peptides (e.g., charge states, missed cleavage sites) for the same cysteine. Experimental data was combined with the requirement for residues to be quantified in at least 2 experiments (e.g., min 4 technical replicates per condition) and median values per condition were reported. Keratins were excluded. The median and SEM values were calculated based on 2 experiments (4 channels) and plotted as bar plots using PRISM. The processed data can be found in Supplementary Table S2.

##### Proteomic platforms: *In situ* TMT-ABPP competition with advanced probes (Related to figures 2A- 2C, S2C, Supplementary Tables S3, S4)

###### Sample preparation

Three 10-plex experiments with primary human T cells and two 10-plex experiments with Jurkat cells were processed according to this protocol (5 experimental conditions in duplicate). Expanded primary human T cells were pelleted (600 *g*, 5 min, 4 °C) and re-suspended in fresh T cell media at 3 x 10^6^ cells/mL for compound treatment. T cells were treated with compounds (EV-96 (10 µM), EV97 (10 µM), PF (5 µM)) or DMSO control in tissue culture flasks (15 mL/flask, 45 x 10^6^ cells/treatment) for 3 h, followed by treatment with EV96A-yne (5 µM) for 1 h. Following the treatment, cells were transferred into Falcon tubes (50 mL) and pelleted (600 *g*, 5 min, 4 °C). Cells were then transferred into new low binding 1.5 mL Eppendorf tubes in cold PBS (1 mL), pelleted (600 *g*, 5 min, 4 °C), flash frozen in liquid nitrogen, and stored at −80 °C until the day of analysis.

On the first day of the mass spectrometry protocol, cell pellets were thawed on ice for 10 min, and cells were lysed by sonication (3 x 8 pulses, 60% output) in ice-cold PBS (560 µL/sample). Protein concentration was measured using a standard DC assay (Bio-Rad) and normalized to 1-2 mg/mL. These samples were further used for mass spectrometry (figure 2A) or gel-based ABPP analysis (figure 2D).

Biotin was conjugated to the modified proteins using copper-catalyzed azide-alkyne cycloaddition reaction. Reagents for the reaction were pre-mixed prior to their addition. Briefly, biotin-azide (10 µL, 5 mM stock in water, 100 µM final concentration), copper sulfate (10 µL, 50 mM stock in water, 1 mM final concentration), tris(benzyltriazolylmethyl)amine ligand (TBTA; 30 µL, 1.7 mM stock in DMSO/*t*BuOH, 1:4, 100 µM final concentration), and tris(2-carboxyethyl)phosphine hydrochloride (TCEP; 10 µL, fresh 50 mM stock in water, 1 mM final concentration) were combined in an Eppendorf tube, vortexed and spun down. The reagent mixture (55 µL) was then added to each sample from the 10-plex experiment, and the samples were incubated at ambient temperature for 1 h. Following the incubation step, samples were transferred on ice and the proteins were precipitated using MeOH/CHCl_3_ precipitation as described above. The protein pellet was re-solubilized in buffer containing 8M urea in PBS (500µL; 4.4 g urea, 4 mL H_2_O, 8 mL final volume) and 10 µL of 10% SDS in PBS with sonication. DTT (25µL, fresh 400 mM solution in H_2_O, 10 mM final concentration) was added to each sample and the samples were heated at 65 °C for 15 min on a heating block. Following this step, the samples were allowed to cool down (5 min) before addition of iodoacetamide (25 µL, fresh 400 mM solution in H_2_O, 10 mM final concentration) and incubation at 37 °C for 30 min.

###### Streptavidin enrichment and trypsin digestion

Following the alkylation step, the samples were transferred to 15 mL Falcon tubes and diluted with SDS-containing PBS buffer (6 mL final volume, 0.2% final SDS concentration in PBS). Pre-washed streptavidin-agarose beads were then added to each sample (50 µL compact beads, 100 µL in PBS/sample) and the mixture was rotated for 2 h at ambient temperature. Briefly, for a 10-plex experiment, streptavidin-agarose beads (1100 µL, 50% slurry) were washed with PBS (2 times x 10 mL) and re-suspended in 1.1 mL of PBS. Following the enrichment step, the beads were pelleted (2000 *g*, 2 min) and washed extensively: 2 x 10 mL of 0.2% SDS in PBS, 1 x 5 mL PBS, then transferred to a new Eppendorf tube and washed 2 x 1 mL water, 1 x 1 mL 200 mM EPPS, pH 8. The final bead slurry was re-suspended in a buffer containing 2M urea in EPPS (200 µL, 200 mM, pH 8). Trypsin (6 μL of 0.33 μg/μL trypsin in trypsin buffer, containing 33 mM CaCl_2_) was added to each sample and the proteins were digested overnight at 37 °C. The following day, the beads were spun down (2000 *g*, 30 s) and transferred to a BioSpin column. The peptides were eluted into a new Eppendorf tube (800 *g*, 30 s) and the beads were washed with extra EPPS (100 µL, 200 mM, pH 8), followed by a second elution (800 *g*, 30 s).

###### TMT labeling

The resulting protein digest (200 µL/sample) was used for TMT labeling. Briefly, acetonitrile (85 µL) was added, followed by the addition of TMT tags (5 µL/sample, 20 µg/µL in dry CH_3_CN) to the corresponding samples. The samples were vortexed, spun down and incubated for 1 h at ambient temperature. The tags were quenched by the addition of NH_2_OH (5 µL, 5% in H_2_O), vortexed, and incubated for 15 min. The samples were acidified by the addition of formic acid (15 µL/sample) and combined in a new Eppendorf tube. The solvent was removed overnight using SpeedVac vacuum concentrator and the final sample was desalted on a SepPak (C18, 50 mg) column as described above. The solvent was removed overnight using SpeedVac vacuum concentrator and the sample was high pH fractionated using HPLC fractionator as described above.

###### TMT liquid chromatography mass-spectrometry

Samples were analyzed by liquid chromatography tandem mass spectrometry using standard 160 min gradient as described above without Real-Time Search feature. Additionally, methionine residues were searched with a differential modification for oxidation (+15.9949 Da).

###### Data processing and analysis

Competition experiments were performed as 10-plex experiments (5 experimental conditions in duplicate) using primary human T cells (3 biological donors) and Jurkat cells (2 biological replicates). Each experiment was processed individually using the following filters: removal of non-unique peptides, removal of half-tryptic peptides, removal of peptides with more than one internal missed cleavage site, removal of peptides with low (< 5,000) average of reporter ion intensities for control channels, and peptides with high variation between the replicate control channels (coefficient of variation > 0.5). Proteins were required to have at least two unique quantified peptides to pass into the final list. Peptide ratios (signal intensity / sum of signal intensities per peptide) were calculated and ratios of all peptides per protein were averaged. The ratios of DMSO/DMSO, EV96(10)/EV96A-yne, EV97(10)/EV96A-yne, PF(10)/EV96A-yne to DMSO/EV96A-yne were calculated and values are provided as percent to DMSO/EV96-yne. Proteins were required to be quantified in a minimum of 2 experiments to be reported in the final table and keratins were excluded. Experiments were combined and replicates were averaged (median) per condition. Proteins were sorted by the EV96(10)/EV96-yne condition, to obtain the ones that have <25% percent to control, which corresponds to >75% competition. This subset was plotted in a heat map (figures 2A-2C and S2C). The processed data can be found in Supplementary Tables S3 and S4.

##### Proteomic platforms: *In vitro* TMT-ABPP competition with scout fragments (related to figure 7, supplementary table S8)

###### Sample preparation

Eight 6-plex experiments with primary human T cells (DMSO/DMSO, DMSO/scout-alkyne, scout/scout- alkyne treatments in duplicate) were processed according to this protocol. Expanded or activated T cells (2 days) were pelleted (600 *g*, 5 min), washed with cold PBS (1 x 10 mL), flash frozen in liquid nitrogen, and stored at −80 °C until the day of analysis. On the first day of the mass spectrometry protocol, cell pellets were thawed on ice for 10 min, and the cells were lysed by sonication (3 x 8 pulses, 60% output) in ice-cold PBS. Protein concentration was measured using a standard DC assay (BioRad) and normalized to 1-2 mg/mL. The samples were treated with DMSO or scout fragment (500 µM) for 1 h at ambient temperature, followed by treatment with DMSO or scout fragment-based alkyne probe (50 µM) for 1 h at ambient temperature. Each 6-plex TMT-ABPP experiment contained the following treatment groups (2 replicates each): DMSO/DMSO, DMSO/50 µM scout fragment probe, 500 µM scout fragment / 50 µM scout fragment probe. Following the *in vitro* reaction with the scout fragment probes, the samples were processed the same way as the *in situ* TMT-ABPP competition experiment with advanced probes (starting at the copper-catalyzed azide-alkyne cycloaddition reaction step).

###### Data processing and analysis

Each experiment was performed with T cells from 2 different biological donors. In total, 8 unenriched proteomics experiments were processed, including KB02 and KB05 treated 6-plex samples in activated and quiescent T cells. Each experiment was processed individually using the following filters: removal of non-unique peptides, removal of half-tryptic peptides, removal of peptides with more than one internal missed cleavage site, removal of peptides with low (< 5,000) average of reporter ion intensities for control channels, and peptides with high variation between the replicate control channels (coefficient of variation > 0.5). Proteins were required to have at least two unique quantified peptides to be reported in the final list and keratins were excluded. Peptide ratios (signal intensity / sum of signal intensities per peptide) were calculated and ratios of all peptides per protein were averaged. The ratio of DMSO/DMSO, scout/scout-alkyne, and DMSO/scout-alkyne was calculated and values are provided as percent to DMSO/scout-alkyne. Each experimental setup was processed individually with the requirement of the proteins to be quantified in at least 2 experiments to be in the final table. Experiments were combined and median values across replicates were calculated per condition. Lower values represent higher levels of competition (e.g., <25% percent to control corresponds to >75% competition). The processed data can be found in Supplementary Table S8.

###### Data processing with SF1 isoform 6 (related to figures 7F-7H, supplementary table S9)

The same data processing and analysis were performed using RAW files from the original publication[1]. FASTA file for SF1 isoform 6 was used for ProLuCid search using Integrated Proteomics Pipeline. Peptide ratios (signal intensity / sum of signal intensities per peptide) were calculated and ratios of all peptides per identified residue were averaged. For each identified peptide, the mean of DMSO_act and DMSO_exp was calculated and if the difference between expanded and activated (exp/act) channels was more than 2-fold, peptides were excluded. For each condition group (exp and act), % to control was calculated (each channel divided by mean of corresponding control). The processed data can be found in Supplementary Table S9.

###### Comparison of scout fragment protein level TMT-ABPP to peptide level TMT-ABPP

Cysteine-directed scout fragment TMT-ABPP data (Master Table S6[1]) was used to calculate maximum competition values for any quantified peptide from a specific protein. Briefly, for each peptide, maximum R values for KB02 and KB05 fragments in activated and quiescent T cells were selected. For each protein, maximum value of all the quantified peptides in a specific experimental condition was selected and the value was converted to the corresponding value showing % signal intensity following competition experiment. This final value was used for comparison with newly acquired protein-directed TMT data. Total number of quantified and ligandable cysteines with KB02 and KB05 is reported. Total number of cysteines in each protein was calculated using FASTA database, containing canonical sequences, from the current study (release-2016_06). Scatterplots were created for each condition and the information on the coloring of scatterplots was included into the final table (Supplementary table S8).

##### Proteomic platforms: Abundance-based proteomics with TMT multiplexing, TMT-exp (related to figure 3, supplementary table S5)

###### Sample preparation

Three 16-plex experiments with primary human T cells from different biological donors were processed (8 experimental conditions in duplicate for each donor). Expanded T cells were pelleted (600 *g*, 5 min, 4°C) and re-suspended in fresh T cell media at 2 x 10^6^ cells/mL for compound treatment. T cells were treated with WX-02-23 (5 µM), WX-02-43 (5 µM), PladB (10 nM)) or DMSO control in 15 cm plates (20 mL/plate, 40 x 10^6^ cells/treatment) with or without αCD3/αCD28 stimulation for 8 h. For treatment conditions that required T cell activation, plates were pre-coated with 15 mL PBS containing αCD3 (5 µg/mL, BioXCell) and αCD28 (2 µg/mL, BioXCell) overnight at 4 °C or for 2 h at 37 °C and washed with PBS (7 mL, 3 times) prior to treatment. Following treatment, cells were transferred into Falcon tubes (50 mL) and pelleted (600 *g*, 5 min, 4 °C). Cells were then transferred into new low binding 1.5 mL Eppendorf tubes using 1 mL cold PBS, pelleted (600 *g*, 5 min, 4 °C), flash frozen in liquid nitrogen, and stored at −80 °C until the day of analysis. On the first day of the mass spectrometry protocol, cell pellets were thawed on ice for 10 min, ice cold PBS containing cOmplete™ EDTA-free Protease Inhibitor Cocktail (1 pellet/10 mL) was added and the cells were lysed by sonication (3 x 8 pulses, 60% output). Protein concentration was measured using a standard DC assay (BioRad) and normalized to 1-2 mg/mL. Each 16-plex TMT-ABPP experiment contained the following treatment groups (2 replicates each): expanded T cells without re-stimulation - DMSO, WX-02-23 (5 µM), WX-02-43 (5 µM), PladB (10 nM); expanded T cells with αCD3/αCD28 re-stimulation - DMSO, WX-02-23 (5 µM), WX-02-43 (5 µM), PladB (10 nM).

Normalized proteomes (100 µL) were added to new 1.5 mL low binding Eppendorf tubes containing pre-weighed urea (48 mg/tube) and vortexed to dissolve urea (8M final concentration). DTT (5 µL, 200 mM, 10 mM final concentration) was added and the samples were heated at 65 °C for 15 min. The samples were allowed to cool down (5 min) before addition of iodoacetamide (5 µL, 400 mM, 20 mM final concentration) and incubation at 37 °C for 30 min. Eppendorf tubes containing the samples were transferred on ice, water was added (400 µL/sample), and the proteins were precipitated using MeOH/CHCl_3_ precipitation as described above. If the protein disc is too small, lower organic layer (CHCl_3_) can be left until the second MeOH addition. The final protein pellet was re-suspended in EPPS (160 µL, 200 mM, pH 8.0, no urea) with sonication, Trypsin/LysC (6 μL of 0.42 μg/μL in Trypsin/LysC buffer, containing 15 mM CaCl_2_) were added, and the digestion was allowed to proceed with shaking at 37 °C overnight.

The following day, peptide concentration was measured using standard BCA assay according to manufacturer’s protocol. For each TMT labeling reaction, 25 µg of proteome was taken, and the total EPPS volume was brough up to 35 µL. Acetonitrile (9 µL/sample) was added, followed by the addition of the corresponding tags (5 µL/sample, 20 µg/µL in dry CH_3_CN). The final acetonitrile content was approximately 25%. The sample was incubated with the tags at ambient temperature for 1 h, followed by incubation with NH_2_OH (5 µL, 5% in H_2_O) for 15 min to quench the remainder of unreacted tags. The final sample was acidified by the addition of FA (2.5 µL/sample).

A ratio check (RC) sample was generated by combining 2 µL of each sample, removing the solvent using SpeedVac vacuum concentrator and desalting using stage tips (10 µL, Thermo Fisher). Briefly, the tips were washed and equilibrated (2 x 20 µL CH_3_CN, 3 x 20 µL buffer A), prior to sample loading. The dry sample was re-suspended in buffer A (20 µL) with sonication and loaded on the stage tip (1,000 *g*, 1 min). The stage tip was washed (3 x 20 µL buffer A; 1,000 *g*, 1 min) and the peptides were eluted (1 x 20 µL buffer B; 1,000 *g*, 1 min) and dried using SpeedVac vacuum concentrator. The ratio check sample was re-suspended in buffer A (10 µL) and analyzed by liquid chromatography mass spectrometry (2 µL injection) using Orbitrap Eclipse mass spectrometer. The following 70 min gradient was used: 5% LC-MS buffer B (CH_3_CN, 0.1% FA) in LC-MS buffer A (H_2_O, 0.1% FA) from 0-10 min, 5%–20% buffer B from 10-50 min, 20%–45% buffer B from 50-55 min, 45%–95% buffer B from 55-57 min, 95% buffer B from 57-59 min, 5% buffer B from 59-61 min, 95% buffer B from 61-63 min, 5% buffer B from 63-70 min). The RC sample allows for calculation of normalization values for the main sample. For a 16-plex experiment, N x 20 µL (N – normalization factor from RC) from each channel were combined and used for further analysis. The desalting step using SepPak columns, high pH HPLC fractionation and LC-MS/MS/MS analysis were performed as described above.

###### Data processing and analysis for TMT-exp experiments

Three 16-plex experiments with primary human T cells from different biological donors were processed (8 experimental conditions in duplicate for each donor). Each experiment was processed individually using the following filters: removal of non-unique peptides, removal of half-tryptic peptides, removal of peptides with more than one internal missed cleavage site, removal of peptides with low (< 5,000) average of reporter ion intensities for control channels, and peptides with high variation between the replicate control channels (coefficient of variation > 0.5). Proteins were required to have at least two unique quantified peptides to pass into the final list. Peptide ratios (signal intensity / sum of signal intensities per peptide) were calculated and ratios of all peptides per protein were averaged. Proteins needed to be quantified in at least 2 experiments to be included in the final table and keratins were excluded. Experiments were combined and median values of all replicates for each condition were reported. The following ratios were calculated and can be found in supplementary table S5: WX-02- 43/WX-02-23 in activated T cells, EV-02-43/PladB in activated T cells, DMSO-act/DMSO-exp (related to figure 3B). Principal component analysis (PCA) was performed using scikit-learn (Pedregosa et al., 2011) library (version v1.0.2) for Python 3.10.4 (or just 3.10). Protein ratio values were log2 transformed and only proteins quantified in all experiments were used (4166/4861) and PC1 and PC2 were plotted (related to figure 3C).

##### Proteomic platforms: SF3B1 co-immunoprecipitation-MS (Related to figure S5C, supplementary table S6)

###### Sample preparation

Expanded or activated T cells (2 days) were pelleted (600 *g*, 5 min) and re-suspended in fresh T cell media at 3 x 10^6^ cells/mL for compound treatment. T cells were treated with compounds (WX-02-23 (5 µM), WX-02-43 (5 µM), or PladB (10 nM)) or DMSO control in tissue culture flasks (15 mL/flask, 45 x 10^6^ cells/treatment) for 3 h. Following the treatment, cells were transferred into Falcon tubes (50 mL) and pelleted (600 *g*, 5 min). The cells were then transferred into new low binding 1.5 mL Eppendorf tubes in ice-cold PBS (1 mL), pelleted (600 *g*, 5 min), flash frozen in liquid nitrogen, and stored at −80 °C until the day of analysis. On the first day of the mass-spectrometry protocol, cell pellets were thawed on ice for 10 min and lysed using ice-cold lysis buffer (100 mM EPPS pH 8, 150 mM NaCl, 0.5% NP40, cOmplete™ EDTA-free Protease Inhibitor Cocktail (1 pellet/10 mL)) with sonication (3 x 8 pulses, 60% output). The Eppendorf tubes were further rotated in the cold room for 15 min to ensure proper lysis and spun down (16,000 *g*, 10 min) to remove remaining insoluble material. Protein concentration of the clarified lysate was measured using a standard BCA assay (Thermo Fisher) and normalized to 1-2 mg/mL. Normalized samples were incubated with SF3B1 antibody (5 µL/sample) or IgG control (5 µL/sample) for 2 h at 4 °C with rotation. Each 10-plex IP-MS experiment was designed to contain the following treatment groups (2 replicates each): expanded T cells – DMSO/IgG, DMSO/SF3B1, WX-02-23 (5 µM)/SF3B1, WX-02-43 (5 µM)/SF3B1, PladB (10 nM)/SF3B1; activated T cells – DMSO/IgG, DMSO/SF3B1, WX-02-23 (5 µM)/SF3B1, WX-02-43 (5 µM)/SF3B1, PladB (10 nM)/SF3B1.

Pre-washed Protein A beads were then added to each sample (25 µL compact beads, 50 µL in IP lysis buffer/sample) and the mixture was rotated for 1.5 h at 4 °C. Briefly, for a 10-plex experiment, Protein A bead slurry (550 µL, 50%) was washed by pelleting (2,000 *g*, 2 min) with IP lysis buffer (2 times x 10 mL) and re-suspended in 550 µL of PBS. Following the enrichment step, the beads were pelleted (2,000 *g*, 2 min) and washed extensively (3 x 1 mL IP lysis buffer, 2 x EPPS (200 mM, pH 8.0), transferring the beads to a new Eppendorf tube with the last wash). The beads were re-suspended in EPPS buffer containing 8M Urea (0.48 mg/mL EPPS, 200 mM, pH 8.0; 40 µL/sample) and heated for 10 min at 65 °C. The samples were transferred to BioSpin columns and the proteins were eluted into new LoBind tubes (2,000 *g*, 1 min). The proteins were reduced with DTT (3.75 µL/sample, 200 mM DTT, 12.5 mM final concentration) for 15 min at 65 °C, cooled down and alkylated with iodoacetamide (3.75 µL/sample, 400 mM IA, 25 mM final concentration) for 30 min at 37 °C. The urea was diluted to 2M with EPPS (202.5 µL, 200 mM, pH 8.0), followed by addition of trypsin (8 µL/sample, 0.25 µg/µL trypsin, 25 mM CaCl_2_) and overnight digest at 37 °C.

###### TMT labeling

The resulting protein digest was used for TMT labeling. Briefly, acetonitrile (90 µL) was added, followed by the addition of TMT tags (5 µL/sample, 20 µg/µL in dry CH_3_CN) to the corresponding samples. The samples were vortexed, spun down, and incubated for 1 h at ambient temperature. The tags were quenched by the addition of NH_2_OH (5 µL, 5% in H_2_O), vortexed, and incubated for 15 min. The samples were acidified by the addition of formic acid (12 µL/sample) and combined in a new Eppendorf tube. Following this step, the samples were processed the same way as the TMT-exp experiments.

###### Data processing and analysis

Six 10-plex experiments with primary human T cells from three different biological donors were processed (5 experimental conditions in duplicate for each donor, activated and expanded T cells). Each experiment was processed individually using the following filters: removal of non-unique peptides, removal of half-tryptic peptides, removal of peptides with more than one internal missed cleavage site, removal of peptides with low (< 5,000) average of reporter ion intensities for DMSO/SF3B1 channels, and peptides with high variation between the replicate DMSO/SF3B1 channels (coefficient of variation > 0.5). Proteins were required to have at least two unique quantified peptides to pass into the final list. Raw SI of peptides was calculated per channel for each protein. Data was median normalized to SF3B1 and technical replicates were averaged (median) per experiment. Proteins were required to show > 2-fold enrichment of DMSO/IgG to be included. The following ratios were calculated per experiment: DMSO/SF3B1 / DMSO/IgG; WX-02-23/SF3B1 / DMSO/SF3B1; WX-02-43/SF3B1 / DMSO/SF3B1; PladB/SF3B1 /

DMSO/SF3B1. Next, ratios were averaged (median) across all experiments per condition (activated and expanded). Proteins needed to be quantified in a minimum of 2 experiments in either condition (activated or expanded) to be included in the final table and keratins were excluded. The processed data can be found in Supplementary Table S6.

##### RNA sequencing and analysis of differentially expressed genes (related to figure 3, Supplementary table 2S)

###### Sample preparation

Small aliquots of T cells from TMT-exp unenriched proteomics experiments (8 x 10^6^ cells/condition) were collected, pelleted (600 *g*, 5 min, 4 °C), transferred to new Eppendorf tubes in cold PBS (1 mL), pelleted (600 *g*, 5 min, 4 °C), flash frozen and stored at −80 °C until the RNA isolation step. Total RNA isolation and on column DNA digestion was carried out from compound or DMSO treated cells using the RNeasy Mini Kit (QIAGEN) following cell homogenization with the QIAshredder kit (QIAGEN) according to the manufacturer’s protocol. RNA concentration was determined using NanoDrop and the samples were stored at −80 °C until sequencing analysis.

###### Data processing and analysis

The quality of initial total RNA was assessed using Agilent Bioanalyzer. Total RNA (100 ng) was used to generate RNA-Seq libraries using Illumina TruSeq stranded mRNA LT kit, following manufacturer’s protocol. The quality and quantity of the Multiplexed libraires was validated using Agilent Tapestation and ThermoFisher NanoDrop. Libraries prepared with unique dual indexes were pooled at equimolar ratios. The pool was denatured and sequenced on Illumina NovaSeq 6000 sequencer using V1.5 reagents, SP flowcell and NovaSeq Control Software V1.7.0 to generate 100 bp single end reads, following manufactures protocol. Raw FASTQ files were trimmed using Trim_Galore (v0.6.4) to remove sequencing adapters and any low quality (Q<15) reads. Trimmed sequencing reads were next aligned to the human Hg19 reference genome (GENCODE, GRCh37.p13) with STAR (v2.7.5) using a two-pass alignment mode with the following parameters: --twopassMode Basic --twopass1readsN -1. SAM files were subsequently converted to BAM files, sorted, and indexed using samtools (v1.9). For visualization of RNA-seq in Integrated Genomics Viewer (IGV; Broad Institute), BAM files were used to generate bigwig files using bamCoverage (part of the Deeptools package; v3.3.1). For differential gene expression analysis, read counting across genomic features (exons) was performed using featureCounts (part of the subread package; v1.5.0) using the following parameters: -p -T 20 -O -F GTF -t exon. Differential gene expression analysis was performed using the edgeR (v3.32.1), DESeq2 (v1.30.1), and limma voom (v3.46.0) R packages. Data visualization and figure generation was performed in Rstudio (v1.3.1073). For quantification of alternative RNA splicing, processed BAM files were used as the input for rMATS (v4.1.1) and against the GENCODE (v35) GTF annotation for GRCh37.p13 using the following parameters: -t paired –libType fr-unstranded –readLength 150 –novelSS. Enumeration of isoform counts was performed using only reads that span the splice junction directly. To identify high confidence AS events, events were considered significant if (i) the inclusion level difference was greater than 20% compared to DMSO, (ii) the False Discovery Rate (FDR) was smaller than 0.05. For comparison of splicing events between conditions, we further limited to highly covered events where at least an average of 50 junctional reads for all conditions. Data analysis and visualization was performed using custom scripts in Rstudio (v1.3.1073) using the following packages: ggplot2 (v3.3.5), ggrepel (v0.9.1), maser (v1.8.0). The processed data can be found in Supplementary Table 2S. The prcomp() and autoplot() functions were used to generate Principal Component Analysis (PCA) plots using gene expression counts as an input.

#### MOLECULAR MODELING

##### Covalent docking (related to figure S2F)

Compounds EV96 and WX-02-23 were prepared using LigPrep in Maestro v13.3.121. To generate tautomers at pH 7.4, we utilized the Ionizer algorithm in LigPrep. For the SF3B1 covalent docking study, we used the crystal structure of SF3B1 in complex with pladienolide D, which has the PDB IDs 7B91. We prepared the protein using the Protein Preparation Wizard in Maestro v13.3.121. EV96 was found to bind covalently to Cys1111 of SF3B1 (Fig. S2F)[5], so we selected Cys1111 as the reactive residue in the Schrodinger CovDock[6]. We defined the Michael addition as the reaction type in CovDock because all compounds contain an acrylamide warhead. We first docked EV96 in the site using CovDock and generated multiple docked poses. When selecting the representative pose, we considered not only a favorable docking/prime score but also the maximization of hydrogen bonds with the site and the minimization of the number of unsatisfied hydrogen bond donors. The Glide gscore of the representative pose in Fig. S2F is −5.8 kcal/mol. Next, we docked WX-02-23 into the site, restricting core docking to the representative pose of EV96 described above. The Glide gscore for WX-02-23 was −7.9 kcal/mol.

#### GENERATION OF REFERENCE PROTEIN TABLES

##### Generation of the Splicing Master List reference protein table (related to figures 6, S6, supplementary table S7)

The splicing master list reference table was consolidated by combining proteins reported to be components of the spliceosome, splicing factors, or splicing regulators (kinases, phosphatases, methyl transferases, and proteins involved in polyadenylation. Specifically, three lists were included: (1) A list of 221 splicing factors published by Blencowe *et al.*[7]; (2) A reference homo sapiens Uniprot list was downloaded (release 2023-01), and the following sections were queried for splicing-related terms (*splicing*, *spliceosome*, etc): protein names, Function [CC], and GO-term annotations (GO, MF, BP); (2) Corum dataset [http://mips.helmholtz-muenchen.de/corum/, download: 9. Dec 2022] was parsed for complexes with a role in splicing by querying for splicing-related descriptions (*splicing*, *spliceosome*, etc) and all members of the identified complexes were included. Three lists were combined, removing duplicate entries, and the final list was manually curated to (1) remove misannotated proteins, and (2) add missing members based on literature curation for splicing regulators (e.g., phosphatases). The final extended list contained 585 entries.

Manually curated splicing factor list was merged with the following resources: (1) Published T cell scout fragment ligandability data (Tables S6 and S2) [1]; (2) Predicted short IDrs (S2 Dataset)[3]; (3) Rbp2go prediction score [https://rbp2go.dkfz.de/, download: 26. Nov 2020]; (4) Corum complex composition and description [http://mips.helmholtz-muenchen.de/corum/, download: 9. Dec 2022]; (5) Interpro domain annotation [https://ftp.ebi.ac.uk/pub/databases/interpro/current_release/match_complete.xml.gz, download: 14. Aug 2023]. All the tables were merged by using Uniprot ID as the identifier. The final Splicing Master List can be found in Supplementary Table S7.

## References

1. Boike, L., N.J. Henning, and D.K. Nomura, Advances in covalent drug discovery. Nat Rev Drug Discov, 2022. 21(12): p. 881–898.

2. Lu, W., et al., Correction: Fragment-based covalent ligand discovery. RSC Chem Biol, 2021. 2(2): p. 670–671.

3. Peczka, N., et al., Electrophilic warheads in covalent drug discovery: an overview. Expert Opin Drug Discov, 2022. 17(4): p. 413–422.

4. Cravatt, B.F., K.L. Hsu, and E. Weerapana, Activity-Based Protein Profiling. 2019: Springer International Publishing.

5. Keller, L.J., et al., Activity-based protein profiling in bacteria: Applications for identification of therapeutic targets and characterization of microbial communities. Curr Opin Chem Biol, 2020. 54: p. 45–53.

6. Vinogradova, E.V., et al., An Activity-Guided Map of Electrophile-Cysteine Interactions in Primary Human T Cells. Cell, 2020. 182(4): p. 1009–1026 e29.

7. Kathman, S.G., et al., Remodeling oncogenic transcriptomes by small molecules targeting NONO. Nat Chem Biol, 2023. 19(7): p. 825–836.

8. Conway, L.P., W. Li, and C.G. Parker, Chemoproteomic-enabled phenotypic screening. Cell Chem Biol, 2021. 28(3): p. 371–393.

9. Bonnal, S., L. Vigevani, and J. Valcarcel, The spliceosome as a target of novel antitumour drugs. Nat Rev Drug Discov, 2012. 11(11): p. 847–59.

10. Jiang, M., et al., SF3B1 mutations in myelodysplastic syndromes: A potential therapeutic target for modulating the entire disease process. Front Oncol, 2023. 13: p. 1116438.

11. Han, C., et al., SF3B1 homeostasis is critical for survival and therapeutic response in T cell leukemia. Sci Adv, 2022. 8(3): p. eabj8357.

12. Hong, D.S., et al., A phase I, open-label, single-arm, dose-escalation study of E7107, a precursor messenger ribonucleic acid (pre-mRNA) splicesome inhibitor administered intravenously on days 1 and 8 every 21 days to patients with solid tumors. Invest New Drugs, 2014. 32(3): p. 436–44.

13. Seiler, M., et al., H3B-8800, an orally available small-molecule splicing modulator, induces lethality in spliceosome-mutant cancers. Nat Med, 2018. 24(4): p. 497–504.

14. Lazear, M.R., et al., Proteomic discovery of chemical probes that perturb protein complexes in human cells. Mol Cell, 2023. 83(10): p. 1725–1742 e12.

15. Lynch, K.W., Consequences of regulated pre-mRNA splicing in the immune system. Nat Rev Immunol, 2004. 4(12): p. 931–40.

16. Arechavala-Gomeza, V., B. Khoo, and A. Aartsma-Rus, Splicing modulation therapy in the treatment of genetic diseases. Appl Clin Genet, 2014. 7: p. 245–52.

17. Martinez, N.M. and K.W. Lynch, Control of alternative splicing in immune responses: many regulators, many predictions, much still to learn. Immunol Rev, 2013. 253(1): p. 216–36.

18. Li, R., et al., Virus usurps alternative splicing to clear the decks for infection. Virol J, 2023. 20(1): p. 131.

19. Boudreault, S., et al., Viral modulation of cellular RNA alternative splicing: A new key player in virus-host interactions? Wiley Interdiscip Rev RNA, 2019. 10(5): p. e1543.

20. Wang, J., et al., Multiple functions of heterogeneous nuclear ribonucleoproteins in the positive single-stranded RNA virus life cycle. Front Immunol, 2022. 13: p. 989298.

21. Li, J. and P. Yu, Genome-wide transcriptome analysis identifies alternative splicing regulatory network and key splicing factors in mouse and human psoriasis. Sci Rep, 2018. 8(1): p. 4124.

22. Ren, P., et al., Alternative Splicing: A New Cause and Potential Therapeutic Target in Autoimmune Disease. Front Immunol, 2021. 12: p. 713540.

23. Stanley, R.F. and O. Abdel-Wahab, Dysregulation and therapeutic targeting of RNA splicing in cancer. Nat Cancer, 2022. 3(5): p. 536–546.

24. Lechner, K.S., M.F. Neurath, and B. Weigmann, Role of the IL-2 inducible tyrosine kinase ITK and its inhibitors in disease pathogenesis. J Mol Med (Berl), 2020. 98(10): p. 1385–1395.

25. Cravatt, B.N.E.; Hayward, R.; DeMeester, K.; Ogasawara, D.; Dix, M.; Nguyen, T.; Ashby, P.; Simon, G.; Schreiber, S.; Melillo, B., Comprehensive Mapping of Electrophilic Small Molecule-Protein Interactions in Human Cells. ChemRxiv, 2023.

26. Zapf, C.W., et al., Covalent inhibitors of interleukin-2 inducible T cell kinase (itk) with nanomolar potency in a whole-blood assay. J Med Chem, 2012. 55(22): p. 10047–63.

27. Foy, A. and M.F. McMullin, Somatic SF3B1 mutations in myelodysplastic syndrome with ring sideroblasts and chronic lymphocytic leukaemia. J Clin Pathol, 2019. 72(11): p. 778–782.

28. Kotake, Y., et al., Splicing factor SF3b as a target of the antitumor natural product pladienolide. Nat Chem Biol, 2007. 3(9): p. 570–5.

29. Steensma, D.P., et al., Phase I First-in-Human Dose Escalation Study of the oral SF3B1 modulator H3B-8800 in myeloid neoplasms. Leukemia, 2021. 35(12): p. 3542–3550.

30. Choi, J.O., J.H. Ham, and S.S. Hwang, RNA Metabolism in T Lymphocytes. Immune Netw, 2022. 22(5): p. e39.

31. Bunnell, S.C., et al., Identification of Itk/Tsk Src homology 3 domain ligands. J Biol Chem, 1996. 271(41): p. 25646–56.

32. Paolino, M., et al., Essential role of E3 ubiquitin ligase activity in Cbl-b-regulated T cell functions. J Immunol, 2011. 186(4): p. 2138–47.

33. Chen, S., S. Benbarche, and O. Abdel-Wahab, Splicing factor mutations in hematologic malignancies. Blood, 2021. 138(8): p. 599–612.

34. Cretu, C., et al., Structural Basis of Splicing Modulation by Antitumor Macrolide Compounds. Mol Cell, 2018. 70(2): p. 265–273 e8.

35. Schneider-Poetsch, T., J.K. Chhipi-Shrestha, and M. Yoshida, Splicing modulators: on the way from nature to clinic. J Antibiot (Tokyo), 2021. 74(10): p. 603–616.

36. Choudhary, G.S., et al., Activation of targetable inflammatory immune signaling is seen in myelodysplastic syndromes with SF3B1 mutations. Elife, 2022. 11.

37. De Arras, L. and S. Alper, Limiting of the innate immune response by SF3A-dependent control of MyD88 alternative mRNA splicing. PLoS Genet, 2013. 9(10): p. e1003855.

38. De Arras, L., et al., Comparative genomics RNAi screen identifies Eftud2 as a novel regulator of innate immunity. Genetics, 2014. 197(2): p. 485–96.

39. Jiang, B., et al., ITK degradation to block T cell receptor signaling and overcome therapeutic resistance in T cell lymphomas. Cell Chem Biol, 2023. 30(4): p. 383–393 e6.

40. Kung, J.E. and N. Jura, Structural Basis for the Non-catalytic Functions of Protein Kinases. Structure, 2016. 24(1): p. 7–24.

41. Mace, P.D. and J.M. Murphy, There’s more to death than life: Noncatalytic functions in kinase and pseudokinase signaling. J Biol Chem, 2021. 296: p. 100705.

42. Dombroski, D., et al., Kinase-independent functions for Itk in TCR-induced regulation of Vav and the actin cytoskeleton. J Immunol, 2005. 174(3): p. 1385–92.

43. Ratni, H., et al., Discovery of Risdiplam, a Selective Survival of Motor Neuron-2 ( SMN2) Gene Splicing Modifier for the Treatment of Spinal Muscular Atrophy (SMA). J Med Chem, 2018. 61(15): p. 6501–6517.

44. Han, T., et al., Anticancer sulfonamides target splicing by inducing RBM39 degradation via recruitment to DCAF15. Science, 2017. 356(6336).

45. Bernard, A., R. Boidot, and F. Vegran, Alternative Splicing in Cancer and Immune Cells. Cancers (Basel), 2022. 14(7).

46. Peng, Q., et al., Impacts and mechanisms of alternative mRNA splicing in cancer metabolism, immune response, and therapeutics. Mol Ther, 2022. 30(3): p. 1018–1035.

47. Schaub, A. and E. Glasmacher, Splicing in immune cells-mechanistic insights and emerging topics. Int Immunol, 2017. 29(4): p. 173–181.

48. Scotti, M.M. and M.S. Swanson, RNA mis-splicing in disease. Nat Rev Genet, 2016. 17(1): p. 19–32.

49. Will, C.L. and R. Luhrmann, Spliceosome structure and function. Cold Spring Harb Perspect Biol, 2011. 3(7).

50. Papasaikas, P. and J. Valcarcel, The Spliceosome: The Ultimate RNA Chaperone and Sculptor. Trends Biochem Sci, 2016. 41(1): p. 33–45.

51. Tsitsiridis, G., et al., CORUM: the comprehensive resource of mammalian protein complexes-2022. Nucleic Acids Res, 2023. 51(D1): p. D539–D545.

52. Coudert, E., et al., Annotation of biologically relevant ligands in UniProtKB using ChEBI. Bioinformatics, 2023. 39(1).

53. Caudron-Herger, M., et al., RBP2GO: a comprehensive pan-species database on RNA-binding proteins, their interactions and functions. Nucleic Acids Res, 2021. 49(D1): p. D425–D436.

54. Paysan-Lafosse, T., et al., InterPro in 2022. Nucleic Acids Res, 2023. 51(D1): p. D418–D427.

55. Bludau, I., et al., The structural context of posttranslational modifications at a proteome-wide scale. PLoS Biol, 2022. 20(5): p. e3001636.

56. Backus, K.M., et al., Proteome-wide covalent ligand discovery in native biological systems. Nature, 2016. 534(7608): p. 570–4.

57. Bar-Peled, L., et al., Chemical Proteomics Identifies Druggable Vulnerabilities in a Genetically Defined Cancer. Cell, 2017. 171(3): p. 696–709 e23.

58. Oberdoerffer, S., et al., Regulation of CD45 alternative splicing by heterogeneous ribonucleoprotein, hnRNPLL. Science, 2008. 321(5889): p. 686–91.

59. Chang, X., B. Li, and A. Rao, RNA-binding protein hnRNPLL regulates mRNA splicing and stability during B-cell to plasma-cell differentiation. Proc Natl Acad Sci U S A, 2015. 112(15): p. E1888–97.

60. Meininger, I., et al., Alternative splicing of MALT1 controls signalling and activation of CD4(+) T cells. Nat Commun, 2016. 7: p. 11292.

61. Nozawa, R.S., et al., SAF-A Regulates Interphase Chromosome Structure through Oligomerization with Chromatin-Associated RNAs. Cell, 2017. 169(7): p. 1214–1227 e18.

62. Knott, G.J., C.S. Bond, and A.H. Fox, The DBHS proteins SFPQ, NONO and PSPC1: a multipurpose molecular scaffold. Nucleic Acids Res, 2016. 44(9): p. 3989–4004.

63. O’Connor, B.P., et al., Regulation of toll-like receptor signaling by the SF3a mRNA splicing complex. PLoS Genet, 2015. 11(2): p. e1004932.

64. Verma, S., et al., SNW1, a Novel Transcriptional Regulator of the NF-kappaB Pathway. Mol Cell Biol, 2019. 39(3).

65. Li, H.B., et al., *m(6)A mRNA methylation controls T cell homeostasis by targeting the IL-7/STAT5/SOCS pathways*. Nature, 2017. 548(7667): p. 338–342.

66. Sledz, P. and M. Jinek, Structural insights into the molecular mechanism of the m(6)A writer complex. Elife, 2016. 5.

67. Sun, Y., et al., Molecular basis for the recognition of the human AAUAAA polyadenylation signal. Proc Natl Acad Sci U S A, 2018. 115(7): p. E1419–E1428.

68. Crowley, V.M., M. Thielert, and B.F. Cravatt, Functionalized Scout Fragments for Site-Specific Covalent Ligand Discovery and Optimization. ACS Cent Sci, 2021. 7(4): p. 613–623.

69. Wang, X., et al., Phosphorylation of splicing factor SF1 on Ser20 by cGMP-dependent protein kinase regulates spliceosome assembly. EMBO J, 1999. 18(16): p. 4549–59.

70. Zhang, Y., et al., *Structure,* phosphorylation and U2AF65 binding of the N-terminal domain of splicing factor 1 during 3’-splice site recognition. Nucleic Acids Res, 2013. 41(2): p. 1343–54.

71. Yoshida, K., et al., Frequent pathway mutations of splicing machinery in myelodysplasia. Nature, 2011. 478(7367): p. 64–9.

72. Scott, K.A.Z., T. L.; Xi,S. Y.; Ngo, B.; Vinogradova, E. V., Protein State-Dependent Chemical Biology. Israel Journal of Chemistry, 2023. 63: p. e202200101.

73. Fernandes, J.A.L., et al., Cryptococcus neoformans Prp8 Intein: An In Vivo Target-Based Drug Screening System in Saccharomyces cerevisiae to Identify Protein Splicing Inhibitors and Explore Its Dynamics. J Fungi (Basel), 2022. 8(8).

74. De Kesel, J., et al., Splicing dysregulation in human hematologic malignancies: beyond splicing mutations. Trends Immunol, 2022. 43(8): p. 674–686.

75. Martinez, N.M., et al., Alternative splicing networks regulated by signaling in human T cells. RNA, 2012. 18(5): p. 1029–40.

76. Pan, Q., et al., Deep surveying of alternative splicing complexity in the human transcriptome by high-throughput sequencing. Nat Genet, 2008. 40(12): p. 1413–5.

## References

1. Vinogradova, E.V., et al., An Activity-Guided Map of Electrophile-Cysteine Interactions in Primary Human T Cells. Cell, 2020. 182(4): p. 1009–1026 e29.

2. Zapf, C.W., et al., Covalent inhibitors of interleukin-2 inducible T cell kinase (itk) with nanomolar potency in a whole-blood assay. J Med Chem, 2012. 55(22): p. 10047–63.

3. Bludau, I., et al., The structural context of posttranslational modifications at a proteome-wide scale. PLoS Biol, 2022. 20(5): p. e3001636.

4. Caudron-Herger, M., et al., RBP2GO: a comprehensive pan-species database on RNA-binding proteins, their interactions and functions. Nucleic Acids Res, 2021. 49(D1): p. D425–D436.

5. Lazear, M.R., et al., Proteomic discovery of chemical probes that perturb protein complexes in human cells. Mol Cell, 2023. 83(10): p. 1725–1742 e12.

6. Toledo Warshaviak, D., et al., Structure-based virtual screening approach for discovery of covalently bound ligands. J Chem Inf Model, 2014. 54(7): p. 1941–50.

7. Han, H., et al., MBNL proteins repress ES-cell-specific alternative splicing and reprogramming. Nature, 2013. 498(7453): p. 241–5.

