## Supplementary material for "Covalent Targeting of Splicing in T Cells": Chemistry SI

#### Supplementary Dataset 3

##### Synthetic procedures and characterization.

##### List of Abbreviations

**$Et_3N$** triethylamine
**DIPEA** *N,N*-diisopropylethylamine
**COMU** (1-cyano-2-ethoxy-2-oxoethylidenaminooxy)dimethylamino-morpholino-carbenium hexafluorophosphate
**DCM** dichloromethane
**THF** tetrahydrofuran
**MeOH** methanol
**FA** formic acid
**TFA** trifluoroacetic acid
**TLC** thin-layer chromatography

##### General considerations

**MY-7A (EV96), WX-01-10, WX-02-23, WX-02-43, WX-02-16, WX-02-36, WX-02-46, WX-03-58** were previously described[1-3].

$^{1}H$ NMR spectra were recorded on Bruker Avance III 400, Avance III HD 400, Avance Neo 400 spectrometers ($^{1}H$, 400 MHz) at 300 K, or a Bruker UltraShield Plus 600 ($^{1}H$, 600 MHz) at 297 K. $^{1}H$ NMR data are reported as follows: chemical shift (δ), multiplicity (s = singlet, d = doublet, t = triplet, m = multiplet; br = broad), coupling constants, and integration. Chemical shifts are reported in parts per million (ppm) using the appropriate solvent as reference[4]. Tandem liquid chromatography/mass spectrometry (LC-MS) was performed on an Agilent 1200 series LC/MSD system equipped with an Agilent G6110A mass detector, alternatively a Waters H-Class LC equipped with diode array and QDa mass detector.

#### General Procedures

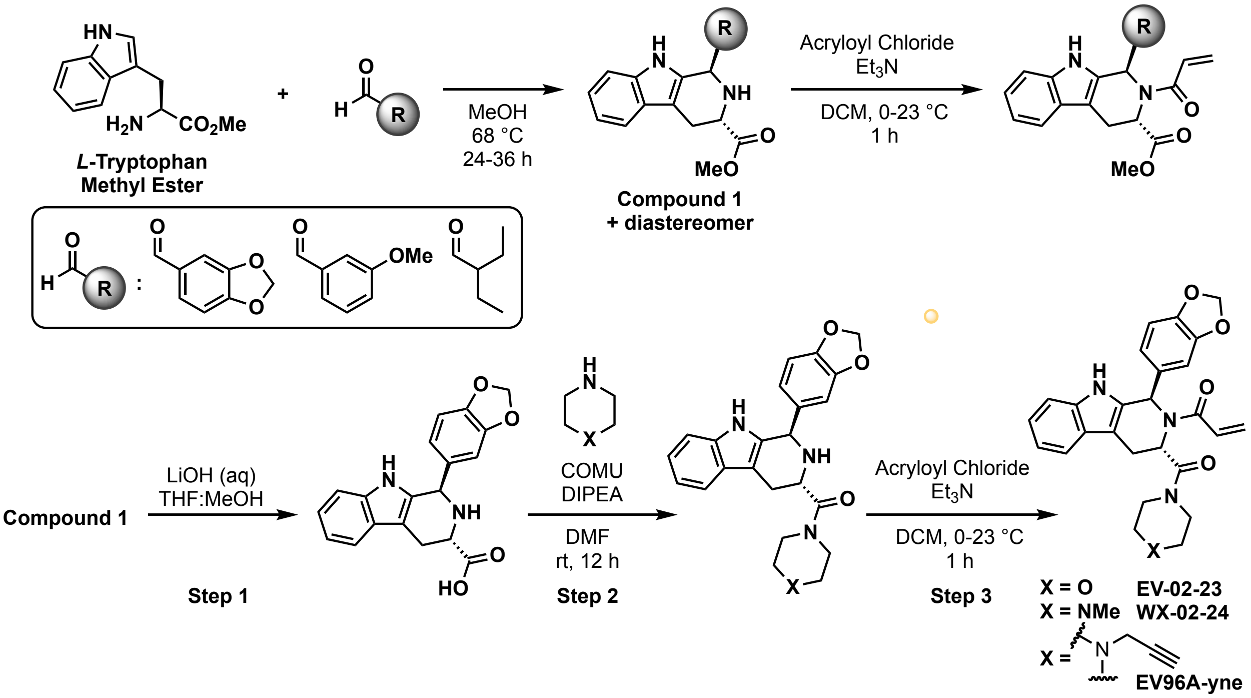


The β-carboline core scaffolds were generated as previously described[1-3]. Briefly, Pictet-Spengler reaction was carried out by combining the appropriate tryptophan methyl ester (1 eq) and the appropriate aldehyde (1.05 eq). The reaction was performed in dry methanol at 68 °C. Reaction progress was monitored by TLC and LCMS to ensure consumption of tryptophan (usually 12-36 h depending on scale). Methanol was removed under reduced pressure, and the crude reaction mixture was taken into ethyl acetate, washed twice with saturated sodium bicarbonate, once with brine, and dried over sodium sulfate. Ethyl acetate was removed under reduced pressure and the crude residue was purified by silica gel chromatography.

**General Procedure 1: Hydrolysis of tryptoline ester.** The product of the Pictet-Spengler reaction (compound 1) was re-suspended in a 3:1 THF:MeOH solvent mixture (0.1 M) and an aqueous solution of LiOH (1 M, 5.0 equiv) was slowly added with stirring. After two hours (or when the reaction was complete by LCMS), the reaction was acidified with 1 M aqueous HCl, diluted with 3 volumes of ethyl acetate and washed with water. Combined organic layers were dried over sodium sulfate and concentrated. Crude residue was purified by column chromatography or taken forward without further purification after the workup.

**General Procedure 2: Amide coupling.** Purified β-carboline free acid was re-suspended in DMF (0.1 M), DIPEA (5.0 eq) and COMU (5.0 eq) were added, followed by the addition of the corresponding amine (5.0 eq). If the piperazine was a TFA salt, an extra equivalent of DIPEA was used for every equivalent of TFA. The reaction was stirred at ambient temperature and the reaction progress was monitored by TLC and LCMS. Upon completion, the reaction was diluted with three volumes of ethyl acetate and washed three times with water and once with brine. The organic layer was dried over sodium sulfate, filtered, and the solvent was removed under reduced pressure. The crude product was purified by column chromatography.

**General Procedure 3: Acrylamide synthesis.** Acryloyl chloride (5.0 eq) was added dropwise to a stirring solution of β-carboline amide in dry DCM (1 M) at 0 °C. After 30 minutes, the reaction was warmed to room temperature and allowed to stir for another 10 min before being quenched with brine. The mixture was diluted with three volumes of ethyl acetate, washed once with saturated sodium bicarbonate solution, and once with brine. The organic layer was dried over sodium sulfate, filtered, and the solvent was removed under reduced pressure. The crude product was purified by preparative TLC.

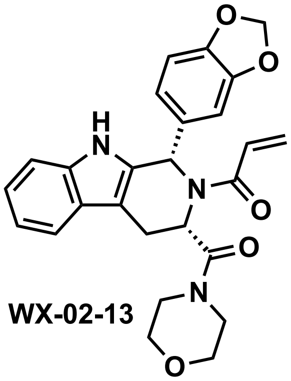

**WX-02-13**

**1-((1S,3S)-1-(benzo[d][1,3]dioxol-5-yl)-3-(morpholine-4-carbonyl)-1,3,4,9-tetrahydro-2H-pyrido[3,4-b]indol-2-yl)prop-2-en-1-one (WX-02-13)**

LC-MS m/z calcd for $C_{26}H_{26}N_3O_5$ $[M+H]^+$ 460.2. Found 460.2.

$^1$H NMR (400 MHz, $CD_3OD$) δ 7.53 (d, *J* = 7.8 Hz, 1H), 7.29 (d, *J* = 8.0 Hz, 1H), 7.25 – 7.07 (m, 2H), 7.07 – 7.00 (m, 1H), 6.95 – 6.81 (m, 2H), 6.81 – 6.71 (m, 1H), 6.64 – 6.44 (m, 0.5H), 6.39 (d, *J* = 16.6 Hz, 1H), 6.03 – 5.89 (m, 3H), 5.88 – 5.73 (m, 0.5H), 3.59 – 3.20 (m, 8H + solvent residual peak), 3.19 – 3.04 (m, 1H), 3.04 – 2.94 (m, 1H), 2.48 (m, 1H), 1 exchangeable proton not observed. [1:1 rotamers]

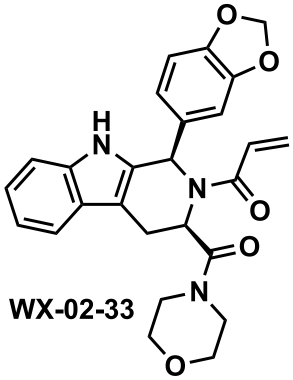

**WX-02-33**

**1-((1*R*,3*R*)-1-(benzo[*d*][1,3]dioxol-5-yl)-3-(morpholine-4-carbonyl)-1,3,4,9-tetrahydro-2*H*-pyrido[3,4-*b*]indol-2-yl)prop-2-en-1-one (WX-02-33)**

LC-MS m/z calcd for $C_{26}H_{26}N_3O_5$ $[M+H]^+$ 460.2. Found 460.2.

$^1$H NMR (400 MHz, $CD_3OD$) δ 7.53 (d, *J* = 7.8, 1H), 7.29 (d, *J* = 8.0 Hz, 1H), 7.23 – 7.07 (m, 2H), 7.03 (ddd, *J* = 8.0, 7.1, 1.1 Hz, 1H), 6.91 – 6.80 (m, 2H), 6.80 – 6.73 (m, 1H), 6.57 – 6.43 (m, 0.5H), 6.39 (d, *J* = 16.6 Hz, 1H), 5.98 – 5.89 (m, 3H), 5.88 – 5.72 (m,

0.5H), 3.66 – 3.20 (m, 8H + solvent residual peak), 3.18 – 3.05 (m, 1H), 2.99 (ddd, *J* = 15.4, 6.2, 1.5 Hz, 1H), 2.67 – 2.28 (s, br, 1H), 1 exchangeable proton not observed. [1:1 rotamers]

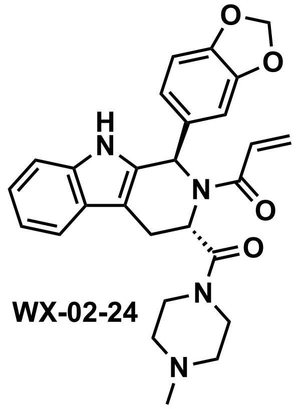

WX-02-24

**1-((1R,3S)-1-(benzo[d][1,3]dioxol-5-yl)-3-(4-methylpiperazine-1-carbonyl)-1,3,4,9-tetrahydro-2H-pyrido[3,4-b]indol-2-yl)prop-2-en-1-one (WX-02-24)**

LC-MS m/z calcd for $C_{27}H_{29}N_4O_4$ $[M+H]^+$ 473.2. Found 473.2.

$^1$H NMR (400 MHz, MeOD) δ 7.47 (d, *J* = 7.7 Hz, 1H), 7.27 (d, *J* = 8.0 Hz, 1H), 7.07 (t, *J* = 7.8 Hz, 1H), 7.01 (t, *J* = 7.4 Hz, 1H), 6.98 – 6.60 (m, 4H), 6.44 – 6.30 (m, 1H), 6.23 (d, *J* = 16.6 Hz, 1H), 5.92 (br. s, 2H), 5.74 (d, *J* = 10.6 Hz, 1H), 5.12 – 4.95 (m, 1H), 3.61 (s, 1H), 3.45 – 3.00 (m, 5H + solvent residual peak), 2.47 – 2.09 (m, 7H), 1 exchangeable proton not observed.

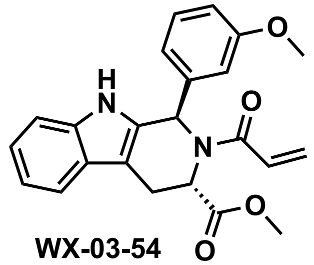

WX-03-54

**methyl (1*R*,3*S*)-2-acryloyl-1-(3-methoxyphenyl)-2,3,4,9-tetrahydro-1*H*-pyrido[3,4-*b*]indole-3-carboxylate (WX-03-54)**

LC-MS m/z calcd for $C_{23}H_{23}N_2O_4$ $[M+H]^+$ 391.2. Found 391.2.

$^1$H NMR (400 MHz, $CD_3OD$) δ 7.44 (d, *J* = 7.8 Hz, 1H), 7.35 – 6.93 (m, 6H), 6.88 – 6.68 (m, 2H), 6.36 – 6.06 (m, 2H), 5.69 (d, *J* = 10.6 Hz, 1H), 5.59 – 5.41 (m, 0.3H), 5.17 – 5.02 (m, 0.7H), 3.76 (s, 3H), 3.70 – 3.39 (m, 4.3H), 3.28 – 3.15 (m, 0.7H), 1 exchangeable proton not observed. [7:3 rotamers]

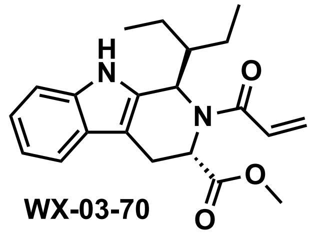

WX-03-70

**methyl (1*R*,3*S*)-2-acryloyl-1-(pentan-3-yl)-2,3,4,9-tetrahydro-1*H*-pyrido[3,4-*b*]indole-3-carboxylate (WX-03-70)**

LC-MS m/z calcd for $C_{21}H_{27}N_2O_3$ $[M+H]^+$ 355.2. Found 355.2.

$^1$H NMR (400 MHz, $CD_3OD$) δ 7.43 (d, *J* = 7.9 Hz, 1H), 7.32 (d, *J* = 8.0 Hz, 1H), 7.12 – 7.04 (m, 1H), 7.03 – 6.97 (m, 1H), 6.85 – 6.68 (m, 1H), 6.27 – 6.09 (m, 1H), 5.77 (d, *J* = 10.5 Hz, 0.7H), 5.69 (d, *J* = 10.6 Hz, 0.3H), 5.48 (s, 0.3H), 5.28 (s, 0.3H), 5.05 (d, *J* = 8.0 Hz, 0.7H), 4.63 (dd, *J* = 9.2, 5.1 Hz, 0.7H), 3.72 (s, 2H), 3.57 – 3.42 (m, 1.4H), 3.30 – 3.24 (m, 1H + solvent residual peak), 3.07 (dd, *J* = 15.8, 5.2 Hz, 0.6H), 2.21 – 2.07 (m, 0.3H), 2.05 – 1.88 (m, 0.7H), 1.85 – 1.29 (m, 4H), 1.20 (t, *J* = 7.5 Hz, 1H), 1.04 (t, *J* = 7.4 Hz, 2H), 0.94 (t, *J* = 7.4 Hz, 2H), 0.84 – 0.71 (m, 1H), 1 exchangeable proton not observed. [7:3 rotamers]

**EV96A-yne**

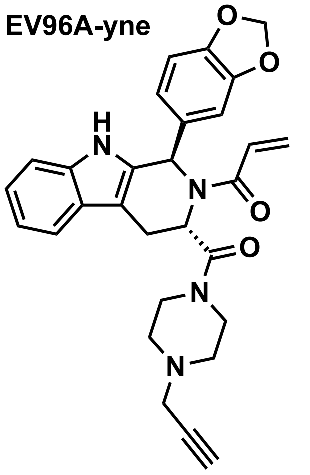


**1-((1*R*,3*S*)-1-(benzo[*d*][1,3]dioxol-5-yl)-3-(4-(prop-2-yn-1-yl)piperazine-1-carbonyl)-1,3,4,9-tetrahydro-2*H*-pyrido[3,4-*b*]indol-2-yl)prop-2-en-1-one (EV96A-yne)**

LC-MS m/z calcd for $C_{29}H_{29}N_4O_4$ $[M+H]^+$ 497.2. Found 497.3.

1H NMR (600 MHz, DMSO-*d*$_6$) δ 10.91 (s, 1H), 7.43 (d, *J* = 7.6 Hz, 1H), 7.27 (d, *J* = 7.9 Hz, 1H), 7.14 – 6.65 (m, 6H), 6.47 – 6.23 (m, 1H), 6.09 (br. s, 1H), 5.97 (d, *J* = 9.9 Hz, 2H), 5.65 (d, *J* = 10.8 Hz, 1H), 5.49 (s, 1H), 3.58 – 3.38 (m, 2H), 3.32 – 3.10 (m, 5H), 2.48 – 2.15 (m, 3H), 1.24 (s, 3H).

#### References


1. Cravatt, B.N., E.; Hayward, R.; DeMeester, K.; Ogasawara, D.; Dix, M.; Nguyen, T.; Ashby, P.; Simon, G.; Schreiber, S.; Melillo, B. , *Comprehensive Mapping of Electrophilic Small Molecule-Protein Interactions in Human Cells.* ChemRxiv, 2023.
2. Lazear, M.R., et al., *Proteomic discovery of chemical probes that perturb protein complexes in human cells.* Mol Cell, 2023. **83**(10): p. 1725-1742 e12.
3. Vinogradova, E.V., et al., *An Activity-Guided Map of Electrophile-Cysteine Interactions in Primary Human T Cells.* Cell, 2020. **182**(4): p. 1009-1026 e29.

4. Fulmer, G.R., et al., *NMR Chemical Shifts of Trace Impurities: Common Laboratory Solvents, Organics, and Gases in Deuterated Solvents Relevant to the Organometallic Chemist.* Organometallics, 2010. **29**(9): p. 2176-2179.

**Spectra**

**WX-02-13 $^1$H NMR ($CD_3OD$)**

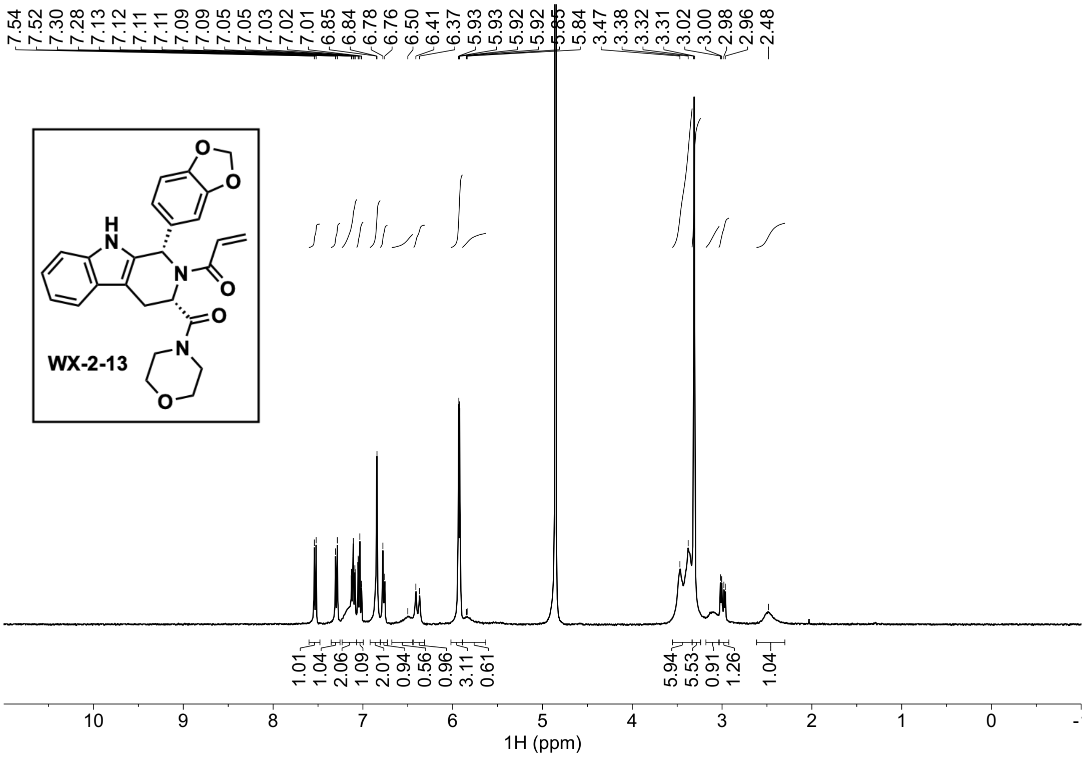

#### WX-02-33 $^1H$ NMR ($CD_3OD$)

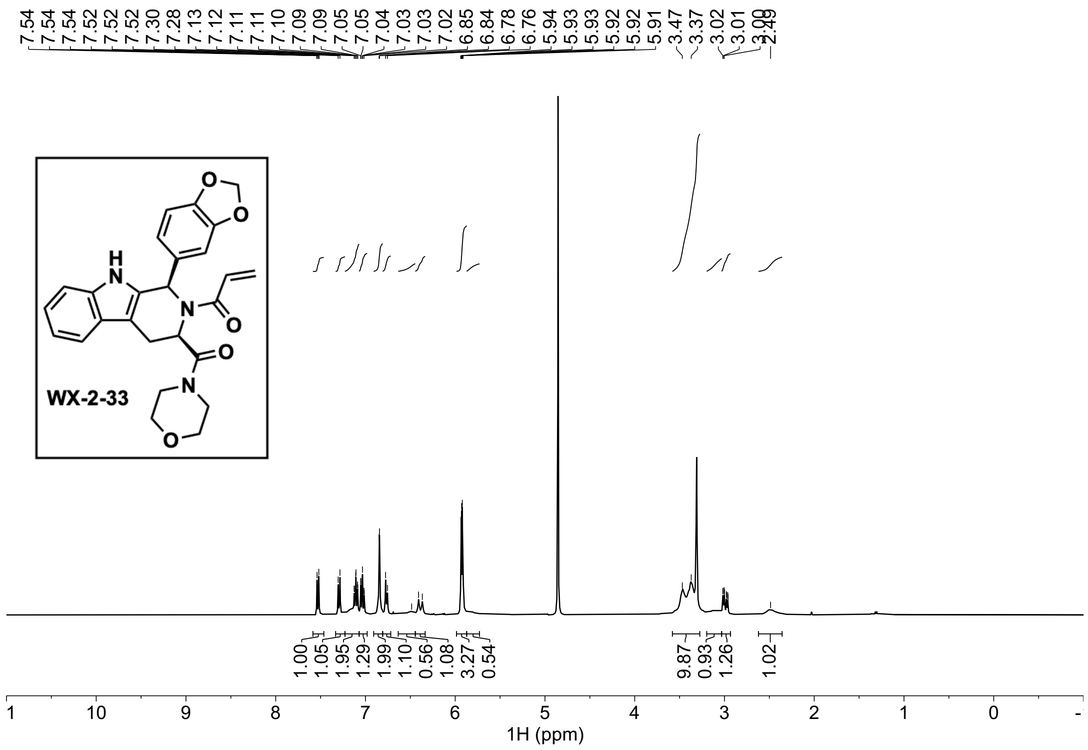


#### WX-02-24 $^1H$ NMR ($CD_3OD$)

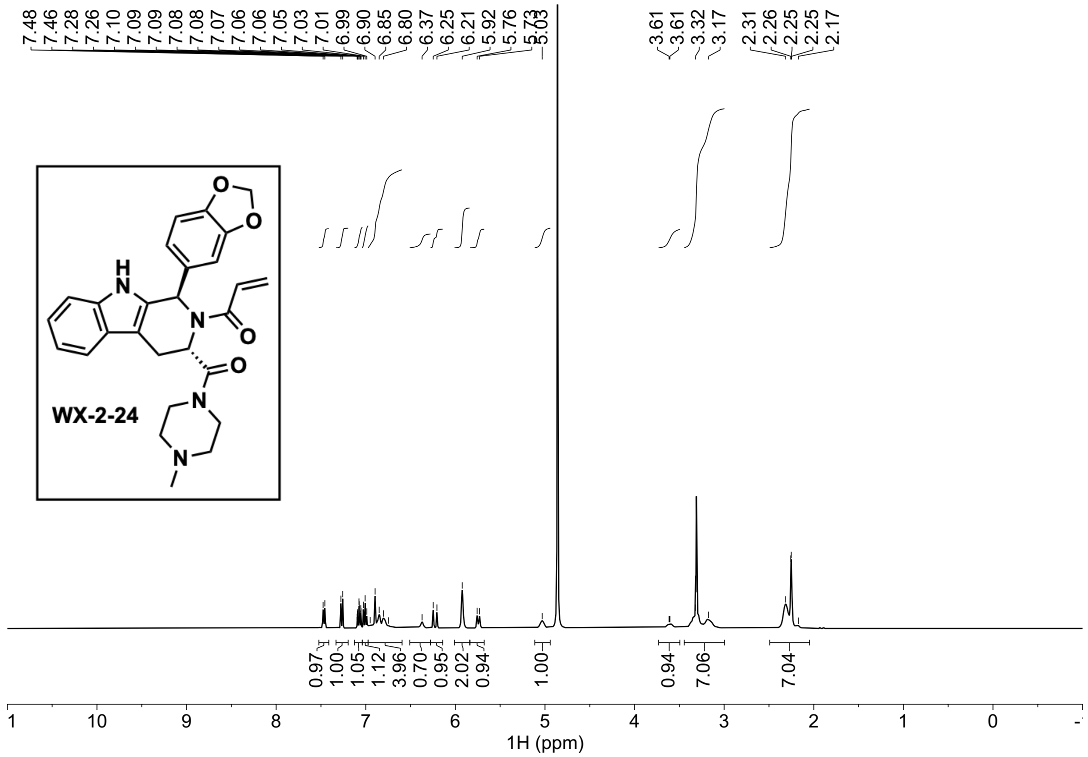

#### WX-03-54 $^1$H NMR ($CD_3OD$)

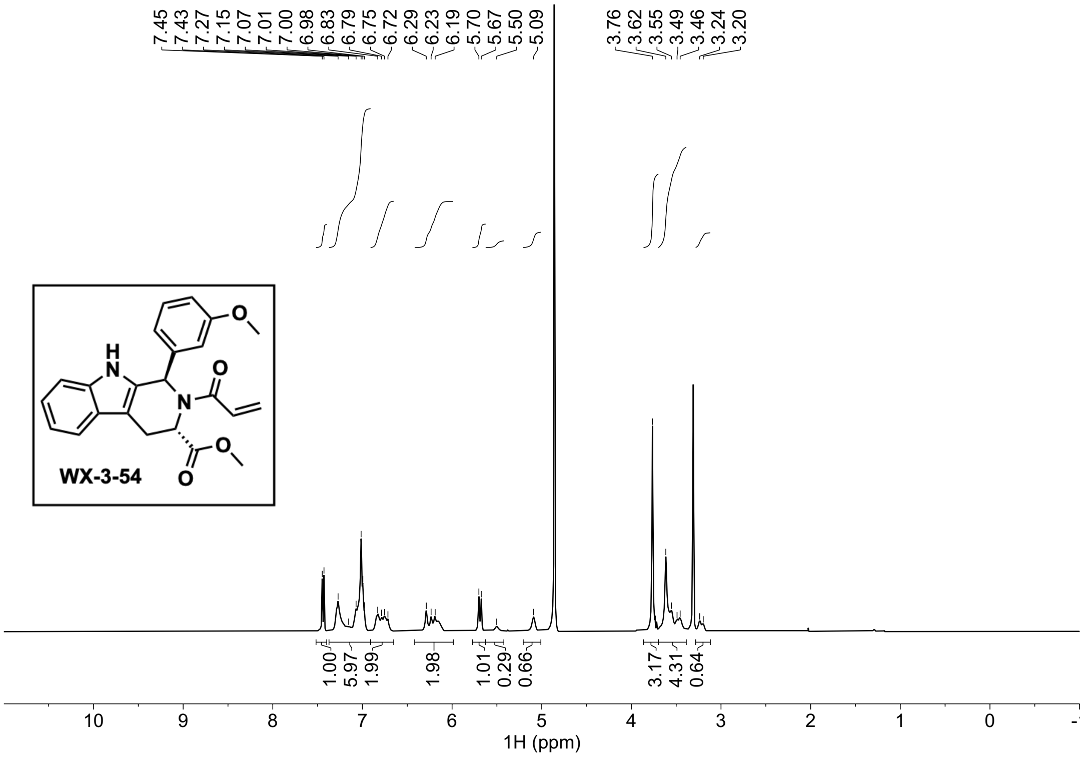


#### WX-03-70 $^1$H NMR ($CD_3OD$)

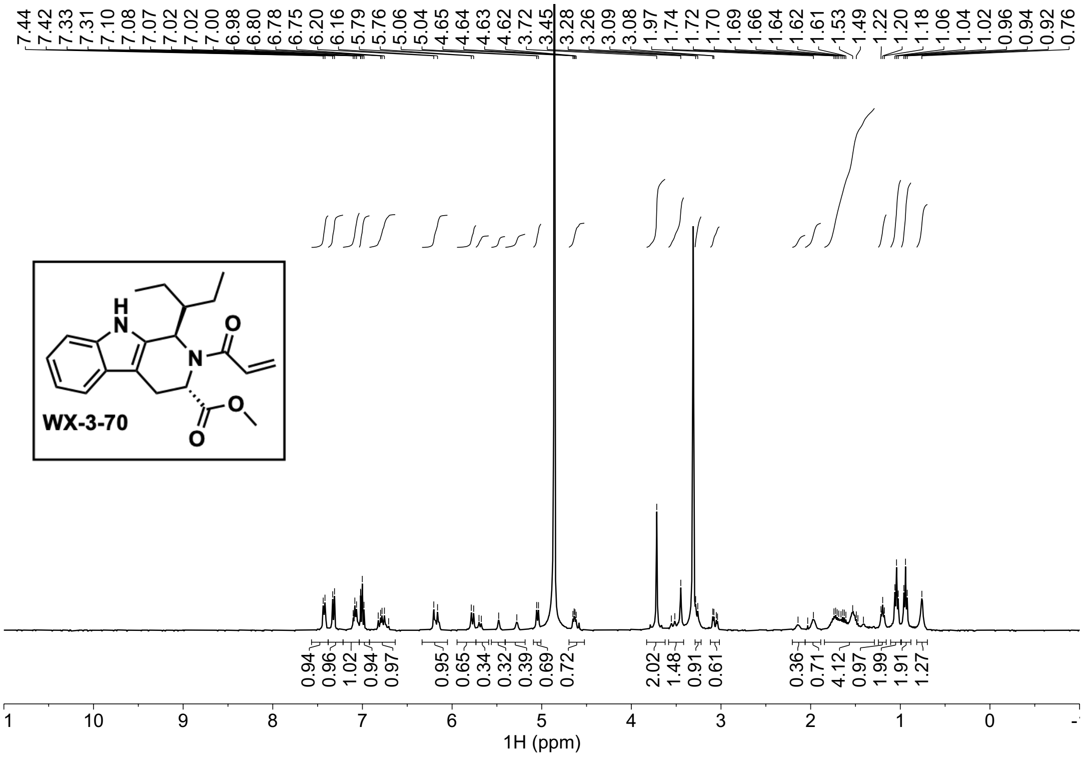

### EV96A-yne $^1H$ NMR (DMSO-$d_6$)

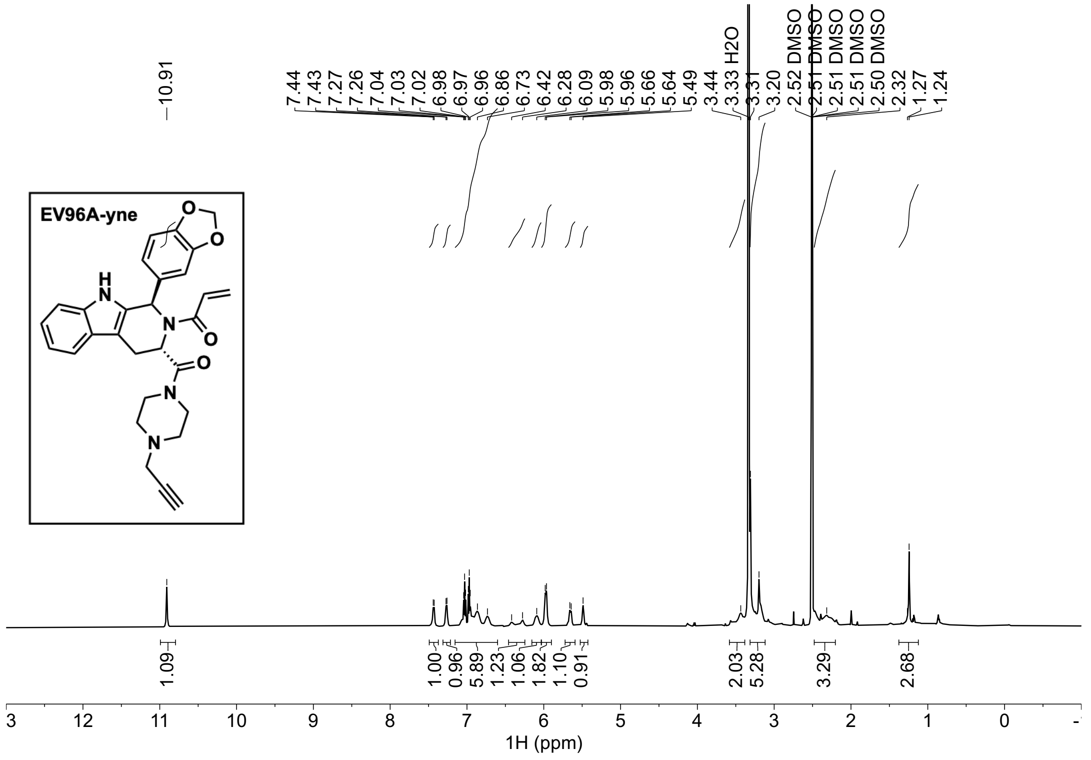
